# Instability of the pseudoautosomal boundary in house mice

**DOI:** 10.1101/561951

**Authors:** Andrew P Morgan, Timothy A Bell, James J Crowley, Fernando Pardo-Manuel de Villena

## Abstract

Faithful segregation of homologous chromosomes at meiosis requires pairing and recombination. In taxa with dimorphic sex chromosomes, pairing between them in the heterogametic sex is limited to a narrow interval of residual sequence homology known as the pseudoautosomal region (PAR). Failure to form the obligate crossover in the PAR is associated with male infertility in house mice (*Mus musculus*) and humans. Yet despite this apparent functional constraint, the boundary and organization of the PAR is highly variable in mammals, and even between subspecies of mice. Here we estimate the genetic map in a previously-documented expansion of the PAR in the *Mus musculus castaneus* subspecies and show that the local recombination rate is 100-fold higher than the autosomal background. We identify an independent shift in the PAR boundary in the *Mus musculus musculus* subspecies and show that it involves a complex rearrangement but still recombines in heterozygous males. Finally, we demonstrate pervasive copy-number variation at the PAR boundary in wild populations of *M. m. domesticus, M. m. musculus* and *M. m. castaneus*. Our results suggest that the intensity of recombination activity in the PAR, coupled with relatively weak constraints on its sequence, permit the generation and maintenance in the population of unusual levels of polymorphism of unknown functional significance.

## Introduction

Pairing, synapsis and crossing-over between homologous chromosomes is a requirement for orderly progression through meiosis that is broadly conserved in eukaryotes (Mather 1938; Mirzaghaderi and Hörandl 2016). In most organisms, pairing is facilitated by sequence homology. The sex chromosomes of mammals – which are divergent in sequence and structure over most of their length (Graves 2006) – thus pose a unique challenge for males, the heterogametic sex. Homology between the X and Y is limited to remnant “pseudoautosomal regions” (PARs) that are a vestige of the sex chromosomes’ origin as an ancestral autosome pair (Ohno 1966). Synapsis and crossing-over in this relatively small territory appears to be necessary for faithful segregation of the sex chromosomes in both humans and mice (Rouyer *et al.* 1986; Matsuda *et al.* 1991; Hassold *et al.* 1991; Burgoyne *et al.* 1992) – although not in some other mammals (Ashley and Moses 1980; Wolf *et al.* 1988; Carnero *et al.* 1991). We limit our discussion here to the placental mammals; the sex chromosomes of marsupials and monotremes are distinct and have more complex meiotic behavior (Sharman *et al.* 1950; Sharp 1982; Graves and Watson 1991; Page *et al.* 2005).

Despite the putative functional importance of the PAR, its boundary and structure appear to evolve rapidly in eutherian mammals (Raudsepp and Chowdhary 2015). PARs have been defined at the sequence level for only a handful of organisms, due in large part to the challenges of assembling repeat- and GC-rich sequences. The human X chromosome has two PARs, PAR1 on Xp and PAR2 on Xq (Figure 1A). Only PAR1 represents the remnant of the ancestral autosome, while PAR2 arose via several rearrangements within the primate lineage (Charchar *et al.* 2003); PAR1 recombines avidly in male meiosis (Hinch *et al.* 2014) while PAR2 recombines infrequently and has sequence properties similar to the rest of the X chromosome (Cotter *et al.* 2016). At least one well-described structural polymorphism of PAR1 segregates in human populations (Mensah *et al.* 2014). The allele is a translocation of X-linked sequence onto the Y chromosome and arose via non-homologous recombination between repetitive elements present on the X and the male-specific portion of the Y chromosome. The duplicated segment recombines in male meiosis at rates similar to single-copy PAR1 (Poriswanish *et al.* 2018). The PARs of simian primates such as the rhesus macaque have sequence homology and conserved synteny with human PAR1 on Xp (Ellis *et al.* 1990) and are on the order of 3 Mb in physical size, while those of ruminants extend further into the ancestral X chromosome and are closer to 10 Mb in size (Das *et al.* 2009).

**Figure 1:**
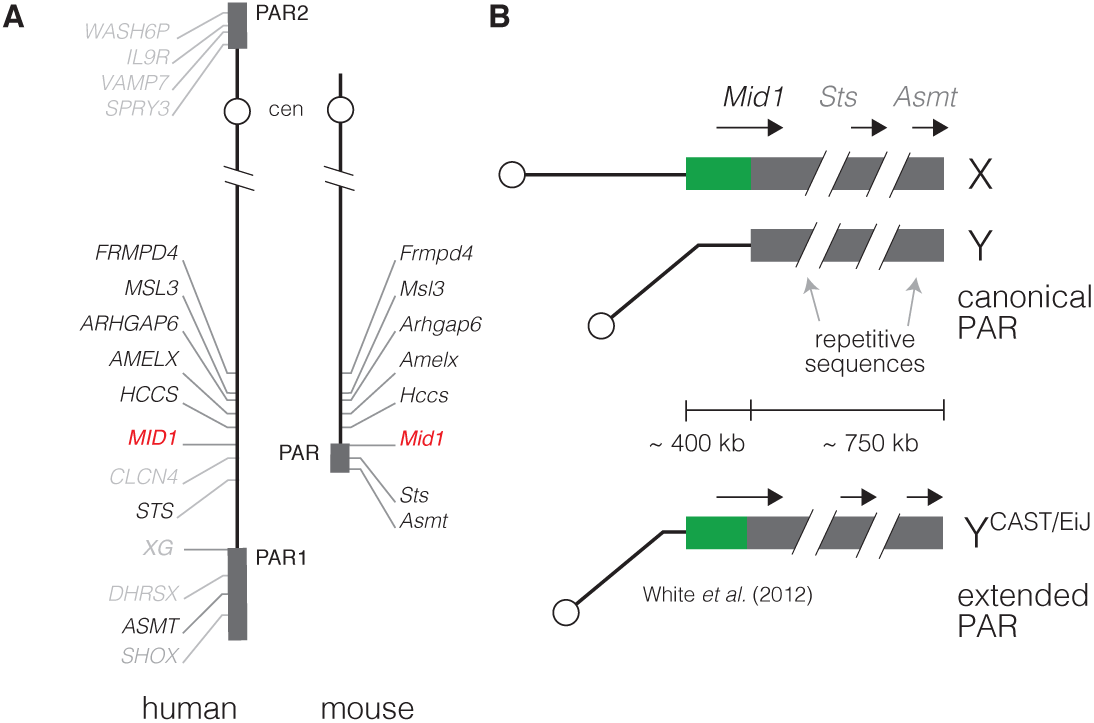
Schematic organization of the pseudoautosomal region in human and the house mouse. (**A**) Synteny map of the human and mouse X chromosomes. Orthologous genes shown in black, non-orthologous genes shown in grey, and the gene marking the end of the human-mouse synteny block (*MID1*) shown in red. PARs shown as grey boxes. Not all human genes are shown. (**B**) The mouse PAR. The “canonical PAR” defined in the C57BL/6J reference strain (top) encompasses at least two genetically-mapped protein-coding genes (*Sts, Asmt*) and the 3’ fragment of *Mid1*. Its physical size is approximately 750 kb, most of which consists of unassembled repetitive sequence. The “extended PAR” of CAST/EiJ (bottom) involves the duplication of approximately 400 kb of ancestrally X-unique sequence onto the Y, including the 5’ fragment of *Mid1* (White *et al.* 2012a).

The mouse PAR, however, is dissimilar to those of humans or other mammals. Its basic organization is shown in Figure 1B. Its boundary falls in the third intron of the mouse ortholog of the human Opitz syndrome (OMIM #300000) gene *MID1* (Palmer *et al.* 1997). Restriction mapping has shown that the mouse PAR is approximately 750 kb in size in the C57BL/6J inbred strain, most of which consists of arrays of GC-rich satellite and other repetitive sequences (Perry *et al.* 2001), and is exquisitely prone to internal rearrangement in male meiosis (Kipling *et al.* 1996a,b). Two genes, *Sts* (Kipling *et al.* 1996a) and *Asmt* (Kasahara *et al.* 2010), have been genetically mapped distal to *Mid1*. Two further genes (*Ngln4x* and *Sfrs17a*) have been localized to the PAR in comparative studies (Bellott *et al.* 2014) but lie in an assembly gap whose sequence is awaiting finishing (https://www.ncbi.nlm.nih.gov/grc/mouse/issues/MG-3246). The representation of the PAR in the current version of the reference genome (mm10/GRCm38) comprises 961, 540 bases (of which 800, 000 lie in two gaps whose size represents a rough estimate by the Genome Reference Consortium) and includes the sequence of *Mid1* and *Asmt*.

The PAR boundary is variable even within the *Mus* genus. It falls between the eighth and ninth exons of *Mid1* in the *Mus spretus*-derived strain SPRET/Ei (Perry *et al.* 2001). White *et al.* (2012a) showed that in the *Mus musculus castaneus*-derived strain CAST/EiJ, the PAR boundary is shifted ∼ 400 kb proximally so that the entire sequence of *Mid1* is present on both sex chromosomes. This implies translocation of previously X-unique sequence into the Y chromosome at the PAR boundary. Recombinants in this extended PAR were recovered from F2 crosses with the *M. m. domesticus*-derived WSB/EiJ strain, and heterozygosity in the PAR was associated with spermatogenic failure (White *et al.* 2012b). Hereafter we refer to the PAR defined in C57BL/6J as the “canonical PAR” and the CAST/EiJ allele as an “extended PAR” (Figure 1B). Unlike the canonical PAR, the extended portion of the PAR in CAST/EiJ consists mostly of unique sequence with GC content and repeat density similar to the genomic background.

Gain or loss of pseudoautosomal identity is predicted to have several important consequences for molecular evolution. First, since pseudoautosomal genes are diploid in both sexes, they are subject to a different dosage-compensation regime than the rest of the X chromosome and are more likely to escape X-inactivation (Graves *et al.* 1998). Second, to the extent that recombination is mutagenic (Hellmann *et al.* 2003), the intensity of pseudoautosomal recombination in male meiosis means pseudoautosomal sequences are expected to diverge more quickly across species than strictly X- or Y-linked sequences. Diversity within populations should also be higher than at sex-linked sites, due to both higher mutational input and mitigation of hitchhiking and background-selection effects by the high local recombination rate in males. Finally, the spectrum of mutation in pseudoautosomal sequence should be biased by the locally increased effects of GC-biased gene conversion (Lamb 1984; Eyre-Walker Adam 1993). These expectations have generally been borne out in empirical studies of the human (Schiebel *et al.* 2000; Filatov and Gerrard 2003; Cotter *et al.* 2016) and mouse (Perry and Ashworth 1999; Huang *et al*. 2005; White *et al*. 2012a) pseudoautosomal regions.

In this work we use crosses between inbred strains from the three major subspecies of house mice to create a genetic map for the both the extended PAR of CAST/EiJ and the proximal portion of the canonical PAR. We use whole-genome sequencing to confirm the presence of a 14 kb extension of the PAR in the *M. m. musculus*-derived PWK/PhJ strain and show that, although it involves a complex structural rearrangement, it too recombines in heterozygous males. Finally, we uncover a remarkable degree of polymorphism at and likely across the PAR boundary in natural populations of all three subspecies. These findings add to a growing body of evidence that the degree of homology required for male fertility is relatively weak (Dumont 2017; Acquaviva *et al.* 2019). Relaxed constraint on PAR structure, combined with the intensity of double-strand break activity in male meiosis, permit the generation and maintenance of unusual levels of diversity in this peculiar region of the sex chromosomes.

## Results

### Recombination in the extended pseudoautosomal region of CAST/EiJ

To estimate a recombination map in the CAST/EiJ extended pseudoautosomal region, we used genotype data from two crosses segregating for the CAST/EiJ Y chromosome: 219 N2 progeny of (PWK/PhJ×CAST/EiJ)F1 males and 48 N2 progeny of (WSB/EiJ×CAST/EiJ)F1 males. The experimental design is shown in Figure 2. For notational convenience we will refer to these crosses by the sire genotype throughout. Progeny were genotyped on one of two SNP array platforms (one with approximately 77, 000 markers, hereafter “77K”, and one with approximately 11, 000 markers, hereafter “11K”) whose content is optimized for populations derived from 8 inbred strains including those used here (Morgan *et al.* 2016). The nominal position of informative markers in the CAST/EiJ extended PAR in the mm10 reference genome in is shown in Figure 2A. To capture recombination in the canonical PAR, we identified 2 additional informative markers beyond the PAR boundary (see **Materials and Methods**).

**Figure 2:**
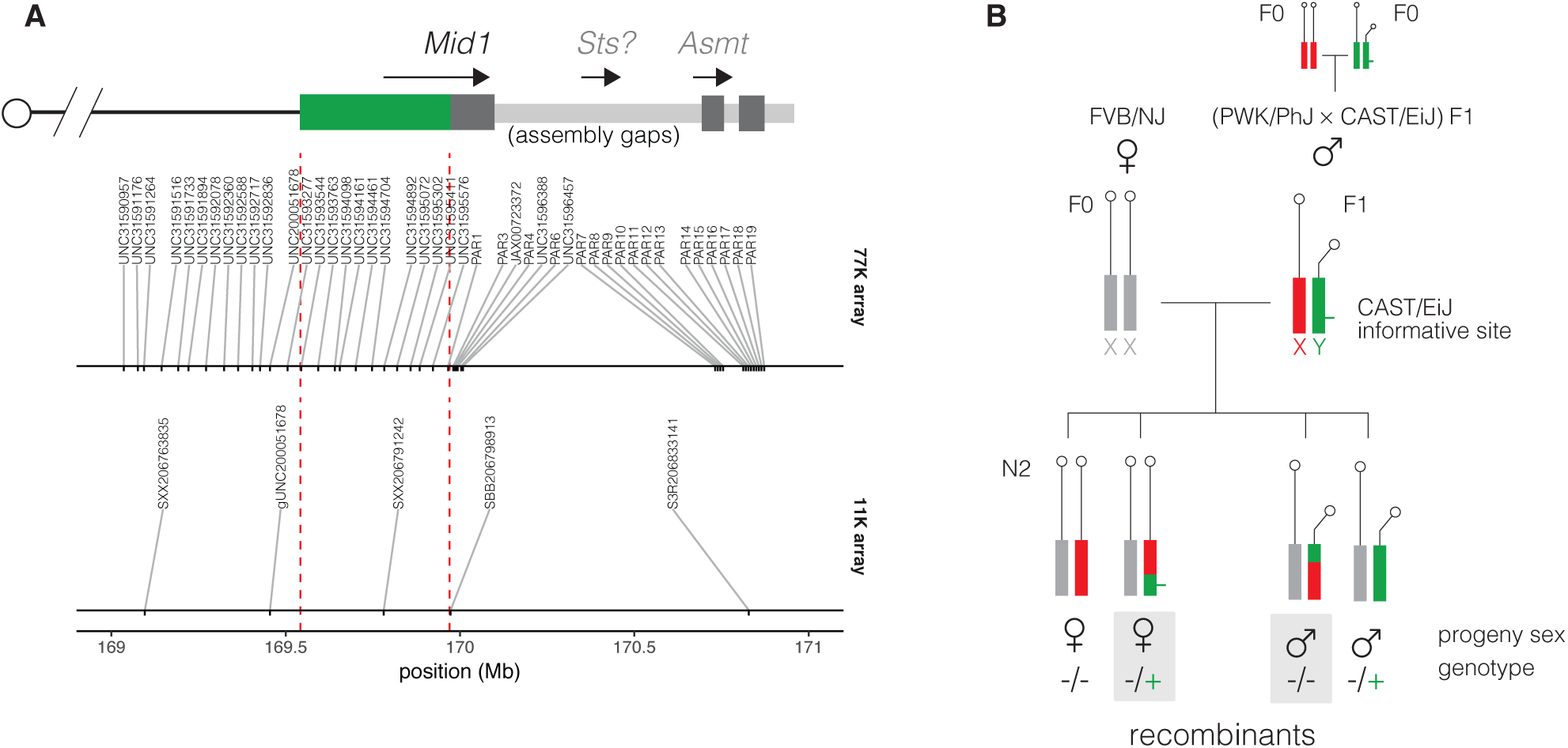
Experimental design for detecting recombination in the extended PAR of CAST/EiJ. (**A**) Marker map near the pseudoautosomal boundary. A schematic representation of the CAST/EiJ extended PAR (green) and canonical PAR (dark grey) with its internal assembly gaps (light grey) and protein-coding genes, is shown in the top panel. Nominal physical positions (in the mm10 assembly) of SNP markers from the 77K and 11K arrays are shown in the bottom panel. CAST/EiJ and canonical PAR boundaries are indicated by red dashed lines. (B) Multiple (PWK/PhJ×CAST/EiJ)F1 males were bred with FVB/NJ females, and N2 progeny genotyped with commercial SNP arrays. In the absence of recombination, female progeny inherit an maternal FVB/NJ allele and a paternal PWK/PhJ allele at X chromosome markers; non-recombinant male progeny inherit a maternal FVB/NJ allele and a paternal CAST/EiJ allele. Other combinations require recombination between the X and Y chromosomes.

Recombinants in the PAR were then identified in curated genotypes using the rules illustrated in Figure 2B. Sex (ie. presence or absence the male-specific region of the Y chromosome) anchors the proximal end of the genetic map of the PAR by definition. Aggregating across both array platforms, 26 of 219 progeny of (PWK/PhJ×CAST/EiJ)F1 males and 8 of 48 of progeny of (WSB/EiJ×CAST/EiJ)F1 males have evidence of recombination. Of these, at least 4 (all from the PWK/PhJ×CAST/EiJ cross) are double recombinants (Table 1). Example recombinant chromosomes from the PWK/PhJ×CAST/EiJ cross are shown in Figure 3A, and the estimated genetic maps for both crosses in Figure 3B (and in tabular form in File S1). The genetic length of the portion of the PAR spanned by our array markers is 13.7±2.3 cM (point estimate ± standard error) and 16.7±5.4 cM in the PWK/PhJ×CAST/EiJ and WSB/EiJ×CAST/EiJ crosses, respectively. (Most of this map, 10.0 ± 2.0 cM and 8.3 ± 4.0 cM, respectively, lies in the “extended” portion of the PAR due to limited marker coverage in the canonical PAR.) Given the region’s physical size of 466 kb (spanning chrX:169,542,082–169,969,759 in the mm10 assembly), this translates to recombination rates of 29.4 ± 5.0 cM/Mb and 35.8 ± 11.5 cM/Mb, respectively, assuming negligible crossover interference (Soriano *et al.* 1987). By contrast, male-specific autosomal rates in these crosses have been estimated at approximately 0.4 cM/Mb (Dumont and Payseur 2011a,b). This 100-fold elevation of the recombination rate in the CAST/EiJ extended PAR over the autosomal background is consistent with obligatory crossing-over in the PAR at every meiosis (Burgoyne 1982) despite its relatively small physical size (Perry *et al.* 2001). We find no evidence for transmission bias against recombinants (pooled OR from both crosses = 0.89 by logistic regression, 95% CI 0.44 - 1.82, *p* = 0.75 by likelihood-ratio test), and a rate of sex-chromosome nondisjunction (XOs: 1/267, 0.37%) similar to prior estimates for these F1s (Dumont 2017).

**Table 1:**
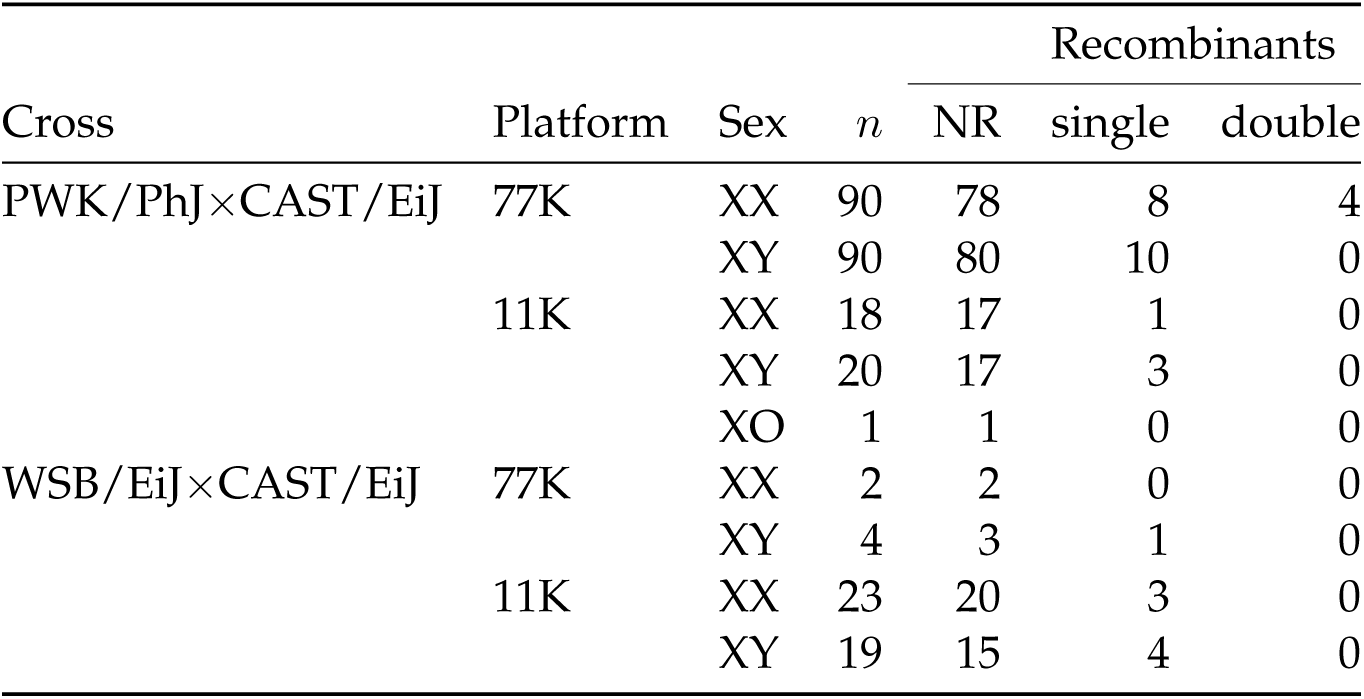
Summary of crossovers detected in the PAR in N2 progeny.

| Cross | Platform | Sex | *n* | Recombinants |  |  |
| --- | --- | --- | --- | --- | --- | --- |
|  |  |  |  | NR | single | double |
| PWK/PhJ×CAST/EiJ | 77K | XX | 90 | 78 | 8 | 4 |
|  |  | XY | 90 | 80 | 10 | 0 |
|  | 11K | XX | 18 | 17 | 1 | 0 |
|  |  | XY | 20 | 17 | 3 | 0 |
|  |  | XO | 1 | 1 | 0 | 0 |
| WSB/EiJ×CAST/EiJ | 77K | XX | 2 | 2 | 0 | 0 |
|  |  | XY | 4 | 3 | 1 | 0 |
|  | 11K | XX | 23 | 20 | 3 | 0 |
|  |  | XY | 19 | 15 | 4 | 0 |

**Figure 3:**
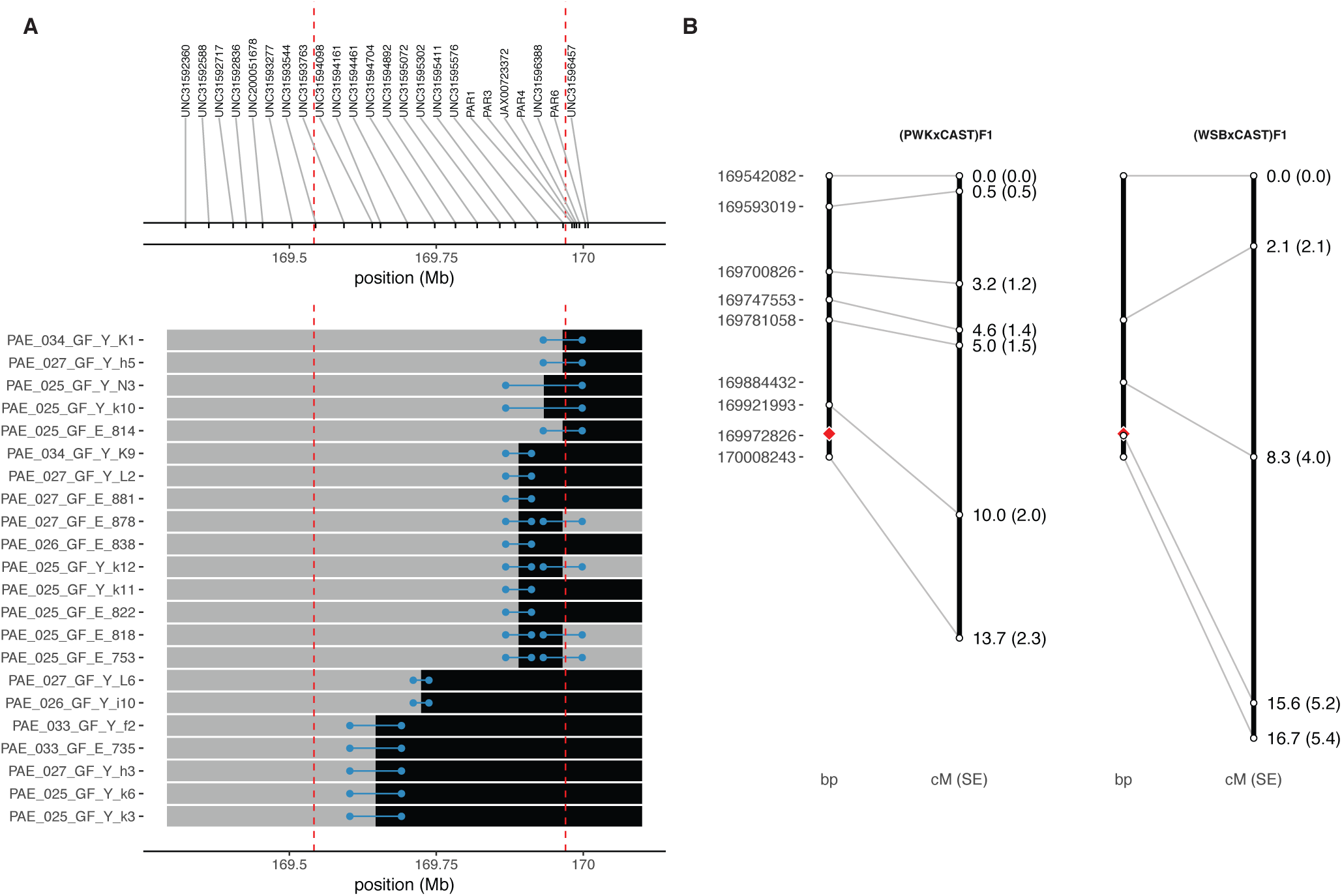
Genetic map of the CAST/EiJ extended PAR. (**A**) Recombinant chromosomes in the extended PAR in the PWK/PhJ×CAST/EiJ cross. The expected non-recombinant haplotype for sex is shown in grey and the recombinant haplotype in black. Blue bars show the interval between flanking markers (with points nudged to avoid overplotting). CAST/EiJ and canonical PAR boundaries are indicated by red dashed lines. Physical positions of markers from the 77K array are shown in the top panel. (**B**) Physical (left) and genetic (right) positions are shown for informative markers in the PWK/PhJ×CAST/EiJ and WSB/EiJ×CAST/EiJ crosses, pooling across both array platforms. Red diamond indicates boundary of canonical PAR.

Two biochemical surrogates of meiotic recombination – histone 3 lysine 4 trimethylation (H3K4me3), which marks sites for DSB formation, and DMC1 protein, which binds free ends of DSBs – both accumulate on the CAST/EiJ Y haplotype in spermatocytes. We re-analyzed ChIPseq performed against H3K4me3 and DMC1 in testes of reciprocal F1 hybrids between C57BL/6J and CAST/EiJ (Baker *et al.* 2015; Smagulova *et al.* 2016). Reads were assigned to parental haplotypes using known SNVs informative between the parental strains. In the extended PAR, H3K4me3 and DMC1 reads are attributed almost exclusively to the X chromosome in CAST/EiJ×C57BL/6J males, but to both haplotypes in C57BL/6J×CAST/EiJ males (Figure S1). This suggests that recombination is initiated from both haplotypes in the extended PAR when the CAST/EiJ Y chromosome is present, and proceeds through the normal homologous recombination pathway.

### Extension of the pseudoautosomal region in PWK/PhJ

We next attempted to estimate the rate of recombination in the canonical PAR. For this we turned to low-coverage whole-genome sequence data from 86 progeny of (WSB/EiJ×PWK/PhJ)F1 sires and FVB/NJ dams (hereafter referred to as the N2 generation). It was immediately apparent that the complex repetitive sequence in the assembled portion of the PAR – to say nothing of the unassembled portion – would preclude accurate genotyping of SNVs from short reads. However, we noticed a copy-number variant (CNV) at the PAR boundary segregating among the N2s that is not present in published whole-genome sequence data from the founders of the cross (FVB/NJ, male; WSB/EiJ, female; PWK/PhJ, female) (Keane *et al.* 2011) but is present in a Collaborative Cross strain carrying a PWK/PhJ Y chromosome (CC044/Unc, (Srivastava *et al.* 2017)) (Figure 4A). The copy gain segregates almost perfectly with male sex: 39 of 40 males (97.5%) have the expected copy number 1 over the adjacent segment of chrX but 2 copies of the CNV segment, while 46 of 46 females (100%) have 2 copies across the entire region (Figure 4B). This pattern is consistent with duplication of X-linked sequence onto the Y chromosome in PWK/PhJ. The segment is thus a candidate for an extension of the PAR. This agrees with the observation of (White *et al.* 2012a) that primers in the second intron of *Mid1* amplify from both X and Y chromosomes in PWD/PhJ, a strain closely-related to PWK/PhJ (Yang *et al.* 2011).

**Figure 4:**
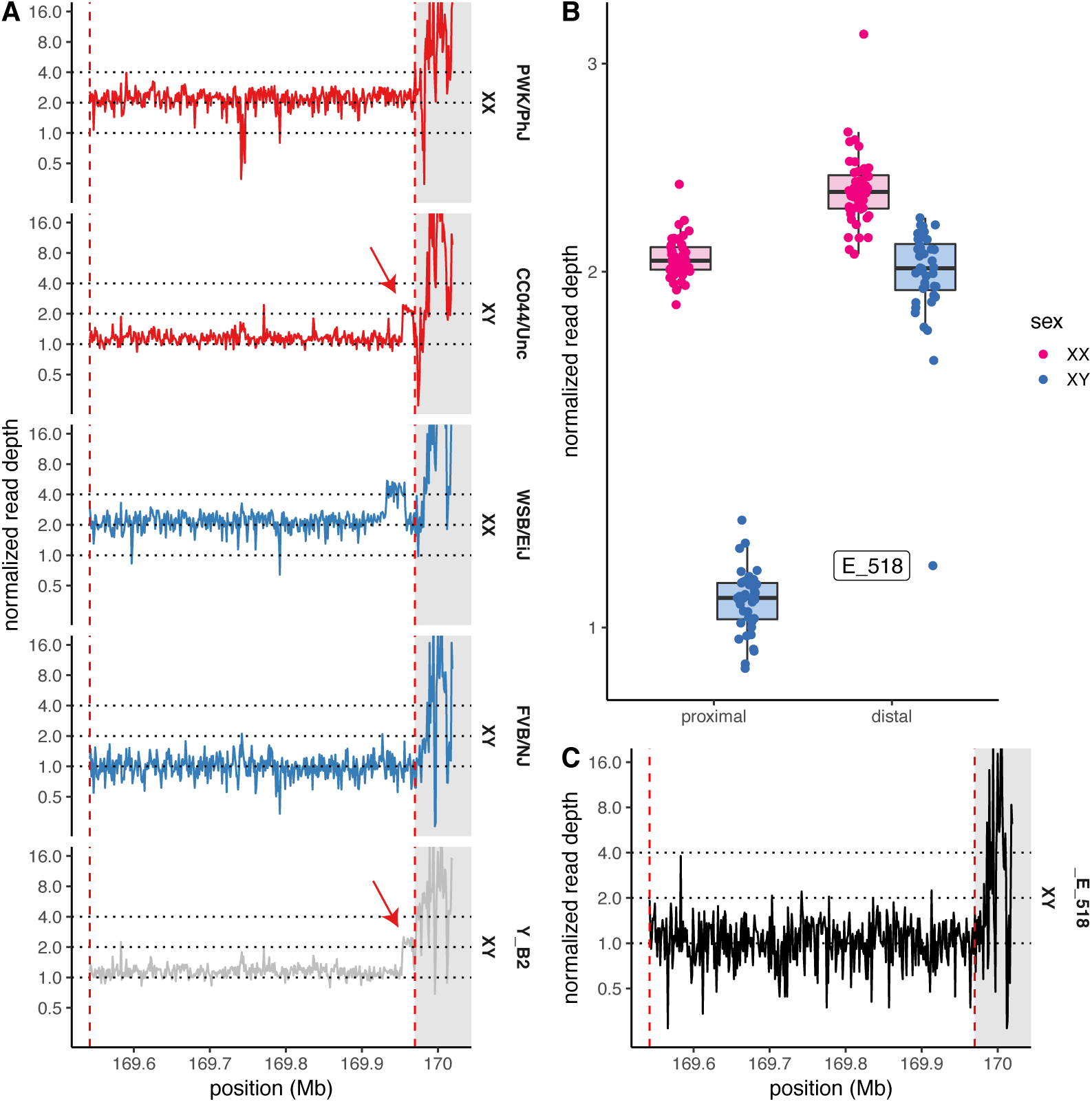
A translocation of X-linked sequence onto the Y in PWK/PhJ. (**A**) Copy number in the distal X chromosome, estimated from whole-genome sequencing read depth in 1 kb bins, for the parental inbred strains (with sex-chromosome karyotypes as shown) of the WSB/EiJ ×CAST/EiJ cross and a representative N2 male (PAE_041_HG_Y_B2). Duplicated segment indicated by red arrow. CAST/EiJ and canonical PAR boundaries are indicated by red dashed lines. Canonical PAR shaded in grey. (**B**) Copy number over the proximal X chromosome (“proximal”) and chrX:169.96 Mb (“distal”, at red arrow in panel A) among N2 males (blue) and females (pink). (**C**) Copy number profile of N2 male PAE_033_HG_E_518, demonstrating *de novo* loss of the duplication.

Closer examination of the duplication shows that it lies near but not immediately adjacent to the PAR boundary in the mm10 reference assembly, and that it is not contiguous (Figure S2). The boundaries of the duplication correspond to clusters of read alignments with aberrant mate-pair orientation. We manually inspected read alignments near the putative duplication boundaries in `IGV` and obtained base-pair resolution of the boundaries by identifying multiple of reads with partial alignments that terminated at the same position. The coordinates of the duplicated segments in mm10 are chrX:169,953,707–169,966,684 and chrX:169,967,172– 169,968,235; the nominal size of the duplicated region is 14,040 bp. Reads from fragments overlapping the duplication boundary were pooled and *de novo* assembled with `SPAdes`. This yielded 3 contigs (File S3) whose alignments back to the reference (with `blat`), shown in Figure S2, define the breakpoints of the duplication. The distal boundary coincides with a cluster of ancient LINE L1 elements, but other boundaries do not appear to correspond to either specific repetitive elements or exons of the *Mid1* gene.

We used the copy number of sub-segments of the CNV and the constraints imposed by the breakpoint alignments to reconstruct the organization of the PAR boundary on the PWK/PhJ Y chromosome (Figure 5). In this proposed configuration, the previously X-linked sequence lies internal to the PWK/PhJ Y PAR; this is supported by direct evidence from single reads whose alignments cross the breakpoint. The duplication also contains 460 bp of novel sequence (in breakpoint contig region_3) that is absent from the mm10 assembly – including unplaced contigs – but is found in *Mid1* introns in *de novo* assemblies for *Mus spretus, Mus caroli* and *Mus pahari*. Recovery of this sequence from multiple outgroup species suggests that it was present in the common ancestor of the genus *Mus* (5 Mya, (Thybert *et al.* 2018)) and has since been lost in some lineages of *M. musculus*. Note that the duplication may be much larger than its nominal size in the mm10 reference, as it may contain other novel sequence that lies too far from the breakpoints to be captured by our assembly strategy.

**Figure 5:**
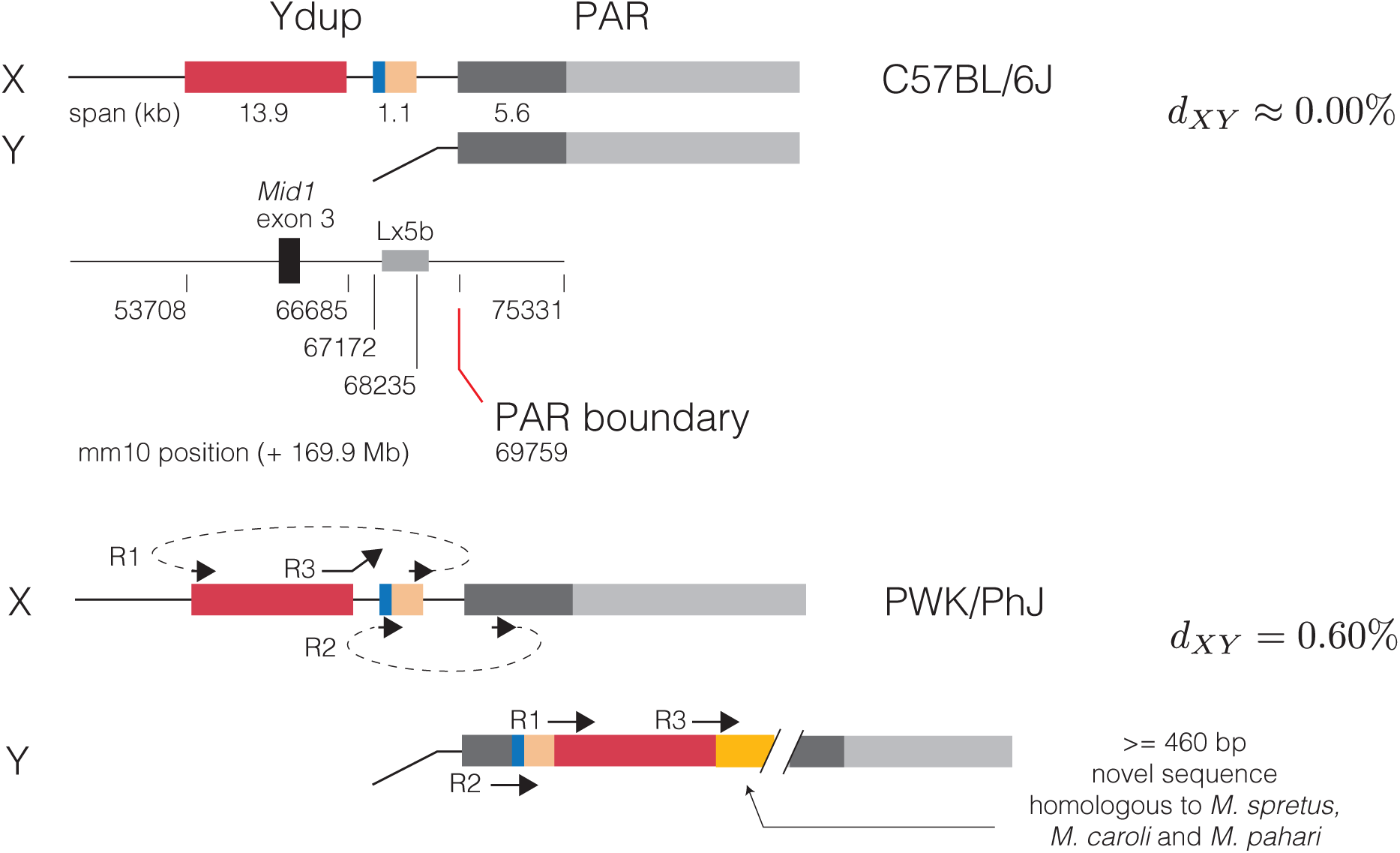
Proposed organization of the PWK/PhJ Y-linked duplication. X-linked segments involved in the duplication are shown in their position in C57BL/6J (mm10 assembly) in the top panel. Breakpoint coordinates and selected annotations are shown in the middle panel. The bottom panel shows alignments of breakpoint-spanning contigs to the PWK/PhJ X chromosome (which is near-identical to C57BL/6J), along with one possible reconstruction of the Y chromosome consistent with copy-number and alignment constraints. *d*_*XY*_, absolute sequence divergence between X- and Y-linked copies of PAR.

Having established that the duplication is genetically and physically linked to the PAR on the PWK/PhJ Y chromosome, we next tested whether it is truly pseudoautosomal in the sense that it recombines in male meiosis. Biallelic SNVs in the duplicated region were ascertained in the N2 progeny, the parental strains of the cross, and a collection of wild mice with publicly-available whole-genome sequence data. Only sites segregating at moderate frequency (minor allele count > 10) in the N2s and heterozygous in a CC male carrying the PWK/PhJ Y chromosome were retained. This conservative ascertainment strategy yielded 84 high-confidence sites informative for recombination in (WSB/EiJ×PWK/PhJ)F1 males. To our surprise, the PWK/PhJ X and Y chromosomes carry different alleles at all 84 sites; we revisit this finding below. Manual review of genotype calls in the N2s revealed 2 likely recombinants (Figure 6): 1 male (PAE_041_HG_Y_B3) with a run of homozygous genotypes, and 1 female (PAE_033_HG_Y_p2) with a run of heterozygous genotypes. The male with copy number 1 (PAE_033_HG_E_518) has reverted to homozygosity across the entire region. We conclude that this individual represents a *de novo* loss of the PWK/PhJ Y-linked copy of the duplicated sequence. Further support for the *de novo* loss is provided by the absence of aberrantly-oriented read alignments at the boundaries of the allele in PAE_033_HG_E_518 but not the male recombinant (Figure S3). The recombination fraction in the duplicated region in this cross, excluding the individual with the *de novo* loss, is 2*/*85 (2.4%) in 28 kb, for an estimated recombination rate of 84 cM/Mb. The duplication is thus a bona fide extension of the PAR.

**Figure 6:**
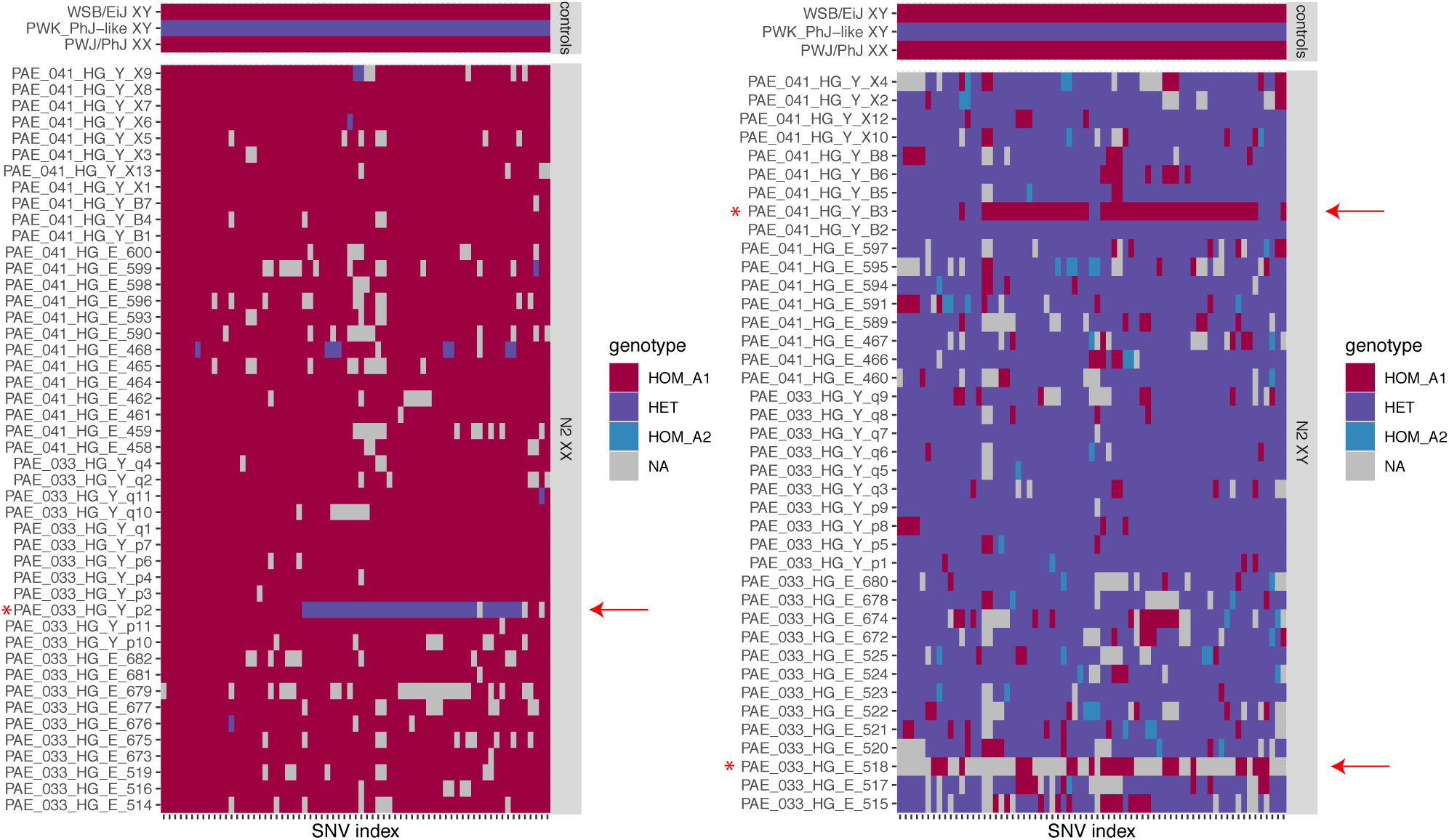
Genotypes of N2 females (left) and males (right) at informative sites between the PWK/PhJ Y chromosome and the PWK/PhJ X or WSB/EiJ X haplotypes under the PWK/PhJ Y-linked duplication. The same set of parental inbred strain controls is shown in both top panels. Genotype calls are color-coded as homozygous for the mm10 reference allele (HOM_A1), heterozygous (HET) or homozygous for the alternate allele (HOM_A2). Missing calls are shown in grey. Recombinant individuals shown with red arrowheads.

### Origin of the extended pseudoautosomal region in PWK/PhJ

Free recombination is difficult to reconcile with the divergence between the X- and Y-linked copies in PWK/PhJ. The two haplotypes differ at 84 sites in 14 kb (0.60%), within the range expected for autosomal sequences from different mouse subspecies (Geraldes *et al.* 2008). PWK/PhJ is a wild-derived strain whose founders were trapped in the Czech Republic (Gregorová and Forejt 2000), has > 90% *M. m. musculus* ancestry (Yang *et al.* (2011) and Figure 7A) and a *M. m. musculus* haplotype in the male-specific portion of the Y chromosome (Morgan and Pardo-Manuel de Villena 2017). However, both PWK/PhJ and the related strain PWD/PhJ are known to carry a *M. m. domesticus* haplotype in the distal portion of the X chromosome (Yang *et al.* 2011). PCA on SNV genotypes both at autosomal sites and within the Y-linked duplication clearly separates wild and wild-derived *M. m. domesticus, M. m. castaneus* and *M. m. musculus* individuals by subspecies (Figure 7A,B). A pseudo-diploid PWK/PhJ Y PAR haplotype, constructed from the consensus PWK/PhJ paternal allele in N2 progeny, clusters with wild *M. m. musculus*, while the PWK/PhJ X chromosome clusters with wild *M. m. domesticus* (Figure 7B). N2 female progeny, expected to be homozygous for two *domesticus* haplotypes (WSB/EiJ and FVB/NJ), cluster with *M. m. domesticus*. N2 male progeny and the CC044/Unc inbred male, expected to be heterozygous between a *musculus-* and a *domesticus-*like haplotype, fall in between the two subspecies-specific clusters. Recombinant N2 progeny lie between the heterozygote and homozygote clusters. The sequence in the PWK/PhJ extended PAR appears to be of *M. m. musculus* origin.

**Figure 7:**
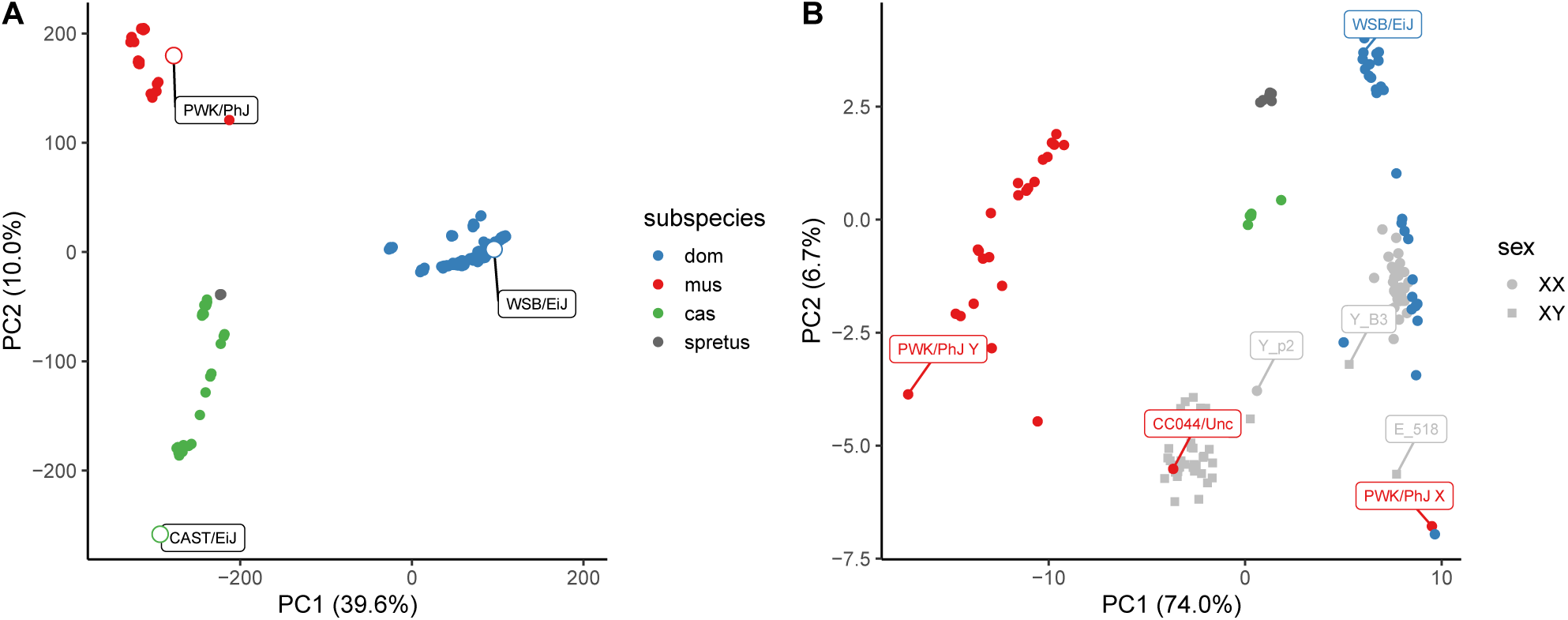
Determining the ancestry of the PWK/PhJ Y-linked duplication. (A) PCA on genotype matrix at 65, 977 autosomal SNPs from the 77K array platform, in a collection of 346 wild mice of *M. m. domesticus, M. m. musculus, M. m. castaneus* and *M. spretus* ancestry, plus the three wild-derived inbred strains used in this study (open points with labels). Data from Didion *et al.* (2016). (B) PCA on 84 sites informative between the PWK/PhJ X and Y chromosomes in wild mice (colored by subspecies) and (WSB/EiJ×PWK/PhJ)N2 progeny.

When did the duplication occur? If it occurred during the establishment of the PWK/PhJ line, it must have been early in the inbreeding process, before the distal X chromosome was fixed for a *domesticus* haplotype. We examined read alignments from wild-caught *M. m. musculus* and identified a male from the Czech Republic (CR16) with clusters of aberrantly-oriented read pairs coinciding with the boundaries of the PWK/PhJ duplication (Figure S4). Reads with partial alignments from this male supported precisely the same breakpoints as in PWK/PhJ. Although we cannot exclude the possibility of recurrent mutation, the most parsimonious explanation is that the duplication in PWK/PhJ is segregating at moderate frequencies in wild populations of *M. m. musculus*.

### Gene expression across the pseudoautosomal boundary

The gene that spans the pseudoautosomal boundary, *Mid1*, is broadly expressed in cell lineages of all three germ layers during embryonic development (Dal Zotto *et al.* 1998) and knockouts have abnormal migration and branching of axons (Lu *et al.* 2013). We therefore investigated the effects of shifts in the PAR boundary on transcriptional regulation of *Mid1*. Our group has previously assayed gene expression by RNAseq in adult brain in male and female progeny of a full diallel cross between CAST/EiJ, PWK/PhJ and WSB/EiJ (Crowley *et al.* 2015). We re-quantified expression in a total of 90 individuals from nine genotypes with `Kallisto` (Bray *et al.* 2016), using the Ensembl version 96 annotation after removal of redundant gene models present on both the X- and Y-encoded copies of the PAR. To characterize patterns of differential splicing, we also aligned reads to the mm10 reference genome after masking the Y-encoded copy of the PAR to avoid mapping artifacts (since the PAR is the only locus that deviates from the conventional haploid representation in the reference genome). Expression of *Mid1* is 11.2-fold higher when at least one CAST/EiJ haplotype is present (95% CI 7.5 - 16.7, adjusted *p* = 9 × 10^-20^), and there is no difference between males and females (1.2-fold higher expression in females, 95% CI 0.8 - 1.6, adjusted *p* = 0.94) (Figure 8A). In reciprocal F1 hybrids between CAST/EiJ and either PWK/PhJ or WSB/EiJ, there is no apparent effect of maternal versus paternal inheritance of the CAST/EiJ haplotype on *Mid1* expression (1.3-fold higher when maternally-inherited, 95% CI 0.8 - 2.1, adjusted *p* = 0.49).

Next we used known sequence variants to assign reads to parental haplotypes and examined coverage on the maternal and paternal alleles in males and females from reciprocal F1 hybrids with a CAST/EiJ parent (Figure 8B). Reads from biological replicates were pooled for improved visualization. In females, *Mid1* is transcribed from both X chromosomes. In males, transcription of the exons in the canonical PAR (exons 4-10, grey boxes) occurs on both X and Y. This 3’ fragment of the gene does not encode an open reading frame, so the role of the Y-linked transcripts is unclear. Transcription of the full-length transcript, including the 5’ fragment proximal to the PAR boundary (exons 1-3, black boxes), occurs only from the X – unless a CAST/EiJ Y chromosome is present.

**Figure 8:**
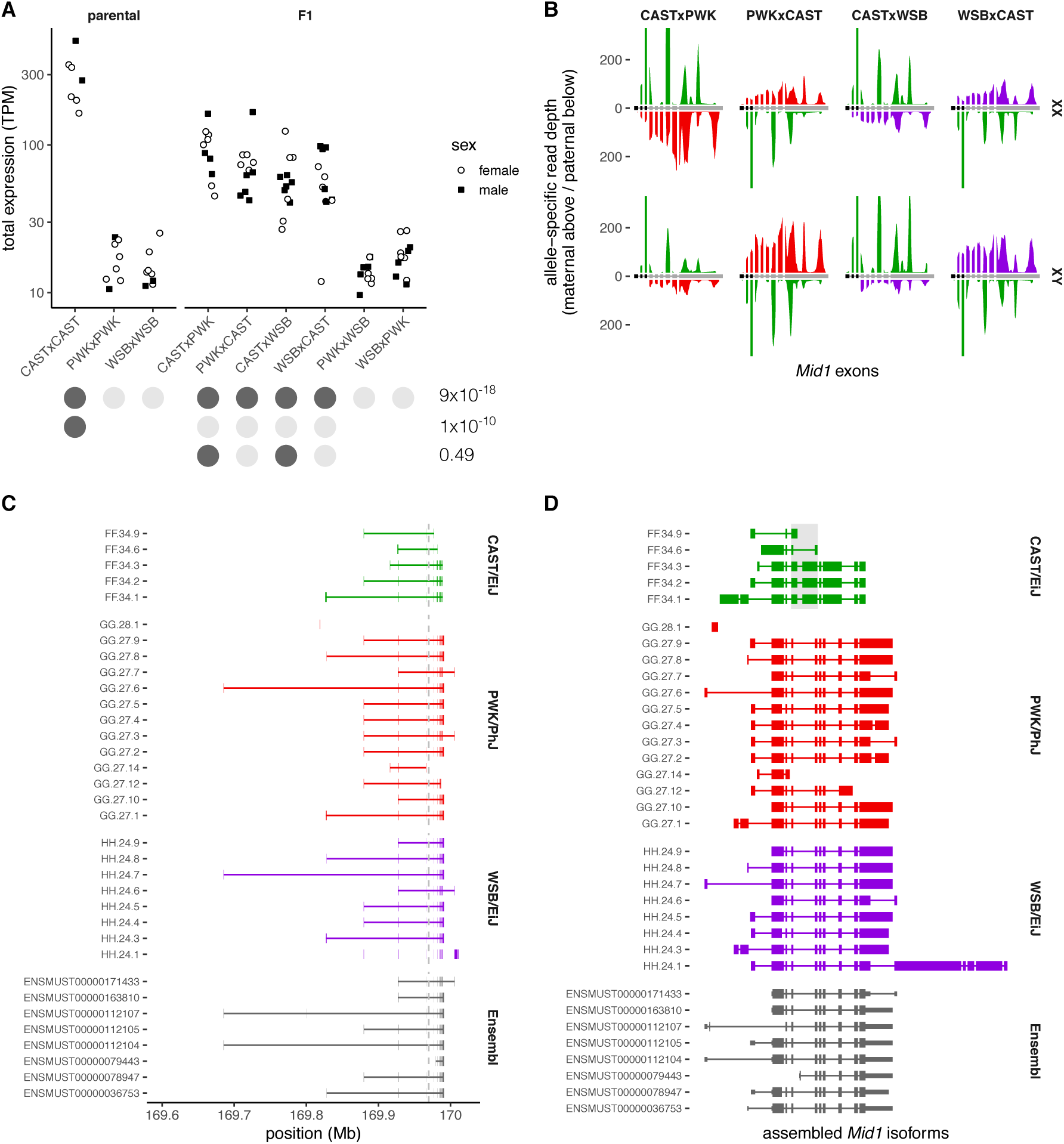
Expression of psueodoautosomal gene *Mid1* in F1 hybrids. (**A**) Total expression in adult whole brain, quantified as transcripts per million (TPM), in all 9 possible reciprocal crosses between CAST/EiJ, PWK/PhJ and WSB/EiJ. Group assignments for several tests of differential expression are shown as color-coded dots beneath *x*-axis, with corresponding adjusted *p*-values at right. (**B**) Aggregate allele-specific read depth over Ensembl-annotated exons of *Mid1* (transcript ENSMUST00000112107) in F1 females (top row) and males (bottom row) with a CAST/EiJ X or Y chromosome. Maternal reads are shown above the horizontal axis and paternal reads below, shaded according to strain origin (green, CAST/EiJ; red, PWK/PhJ; purple, WSB/EiJ). (**C**) Transcript isoforms assembled from RNAseq read alignments in the parental strains compared to *Mid1* transcript models from Ensembl, plotted in genomic coordinates. Grey dashed line shows canonical PAR boundary. (**D**) Same transcripts as panel C, but drawn with introns compressed, exons to scale and vertically aligned to facilitate comparison of splice sites. Coding region of Ensembl transcripts shown as tall boxes. CAST/EiJ splicing differences in exons corresponding to canonical exons 4 and 5 emphasized by light grey shaded box.

Finally we learned transcript models from read alignments in the parental strains and compared them to existing annotations in Ensembl (Figure 8C-D). Transcripts containing the full-length coding sequence of the MID1 protein were recovered from PWK/PhJ and WSB/EiJ but not from CAST/EiJ, despite having nearly 100, 000 reads aligned in the *Mid1* locus in that strain. Although we observed hundreds of reads originating from exons 6-10 in CAST/EiJ (Figure 8B), none had spliced alignments across the junction between exons 5 and 6, nor between 7 and 8. Several CAST/EiJ isoforms have a 3’ extension of exon 4 and a 5’ extension of exon 5 (grey highlighted box in Figure 8D); the coding potential of these isoforms is uncertain. The difficulty of assembling full-length transcripts in CAST/EiJ may be due to mapping artifacts if the organization of this portion of the PAR is divergent from the reference genome. Moreover, it is difficult to assess the accuracy of our transcript assembly in the presence of abundant copy-number variation of segments that include exons 4-10 (next section).

### Extreme structural diversity at the pseudoautosomal boundary

Inspection of sequencing data from 67 wild mice revealed abundant copy-number variation in the 1 Mb adjacent to the PAR boundary as well as within the canonical PAR itself (Figure 9A, Figure S5). Nominal copy number varies from zero to more than 20 for some segments, and individual mice may have 4 or more distinct copy-number states across the region. As for the PWK/PhJ allele, we sought to identify breakpoints of copy gains and losses by manual review of read alignments in `IGV` for 38 mice with evidence of CNVs in the distal X chromosome (excluding the canonical PAR). Clusters of 3 or more reads with partial alignments terminating at the same base pair in the mm10 reference were deemed sufficient evidence for a breakpoint. This analysis yielded 32 distinct breakpoints within 200 kb of the PAR boundary, none of which are shared across subspecies. Their spatial distribution appears non-random: their density seems to decrease moving outward from the PAR boundary (Figure 9B, File S9). However, tests for enrichment of breakpoints in proximity to exons and repetitive elements in the reference genome, and known recombination hotspots, all returned null results (cf. Figure 9C-E). The density of repetitive elements in the 500 kb of sequence proximal to the pseudoautosomal boundary (42.8%) is not different than the subtelomeric regions of the autosomes (mean 42.0%, standard deviation 10.2%). The previously-described ∼ 466 kb expansion of the PAR in the CAST/EiJ strain (White *et al.* 2012a), of predominantly *M. m. castaneus* ancestry and derived from founders trapped in Thailand, is not found in wild *M. m. castaneus* from northern India. The allele may be confined to southeast Asia. Several large deletions adjacent to and possibly involving the canonical PAR boundary that were also identified in CAST/EiJ appear fixed in the Indian population.

**Figure 9:**
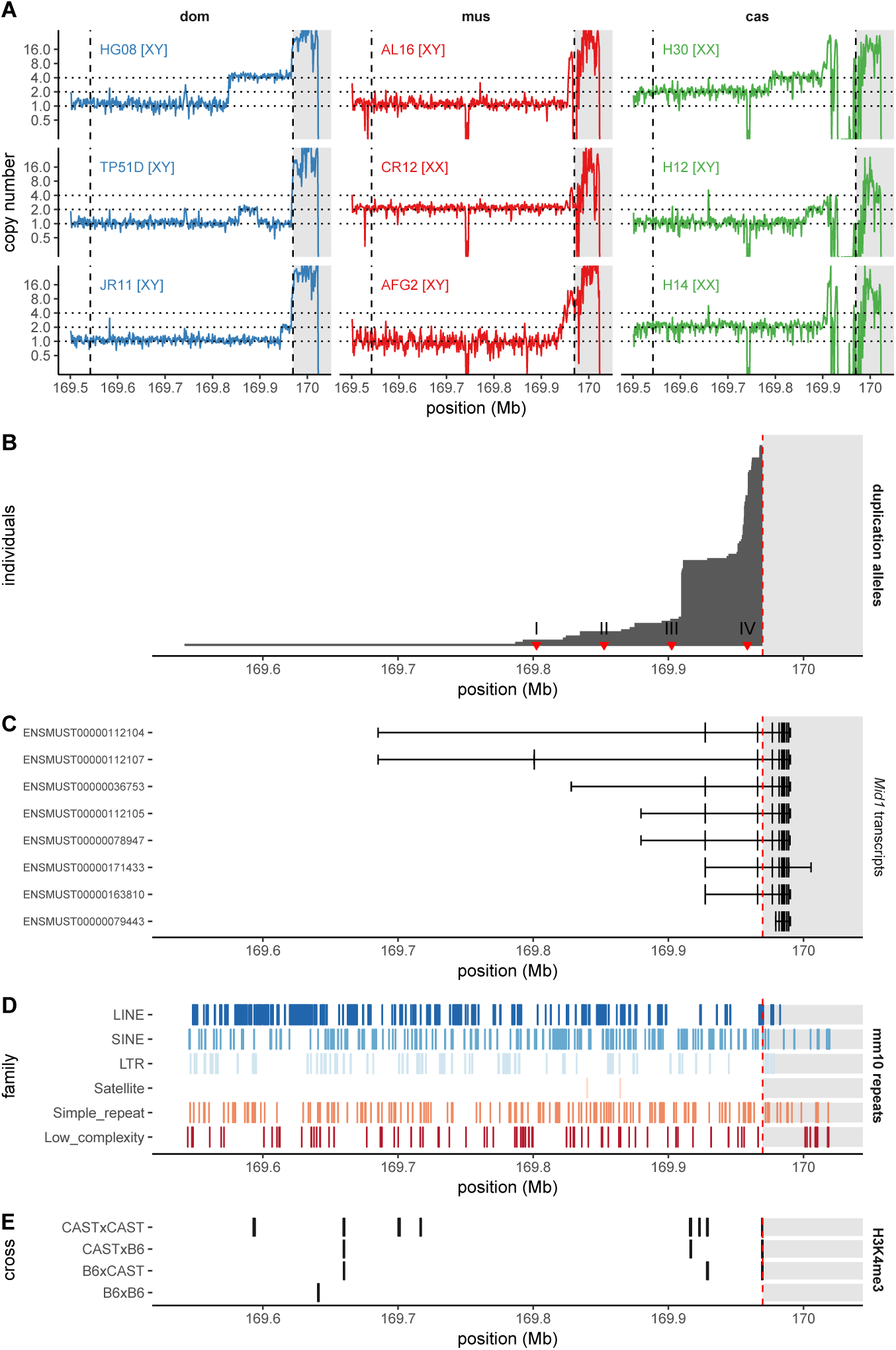
Abundant copy-number variation at the pseudoautosomal boundary in wild house mice. (**A**) Copy number in the distal X chromosome, estimated from whole-genome sequencing read depth in 1 kb bins, for representative individuals of *M. m. domesticus* (blue), *M. m. musculus* (red) and *M. m. castaneus* (green). Inferred sex-chromosome karyotype is shown for each individual. Dashed lines show the boundaries of the CAST/EiJ extended and canonical PARs, respectively. Canonical PAR is shaded grey. (**B**) Distinct copy-number alleles ascertained from breakpoint-spanning reads in 67 wild mice. Each allele is represented as a rectangle extending from the proximal breakpoint up to the pseudoautosomal boundary, and is shown in proportion to the number of times it is observed in our sample. (**C**) Transcripts of the *Mid1* gene from Ensembl release 96. (**D**) Repeat elements annotated in the mm10 reference by the UCSC Genome Browser. (**E**) Recombination hotspots ascertained in C57BL/6J (B6), and CAST/EiJ (CAST), and F1 males (Baker *et al.* 2015).

To gain some insight into the evolutionary trajectory of the PAR in the genus *Mus*, we estimated copy number in the distal X chromosome and the canonical PAR in 8 *M. spretus*, 1 *M. spicilegus* and 1 *M. caroli*. Both males and females have the expected copy number for the X chromosome in the vicinity of the PAR boundary (Figure S6). Within the most proximal region of the canonical PAR, from the PAR boundary to just past the eight exon of *Mid1* (chrX: 169986134), outgroup samples of both sexes have the copy number expected for the X chromosome – suggesting that the region is *not* pseudoautosomal in these species. (Similarly, no evidence of heterozygosity in the proximal exons of *Mid1* was found in whole-exome data from wild-caught male representatives of more distant outgroups *Mus cookii* and *Nannomys minutoides* (Sarver *et al.* 2017); not shown.) This agrees with previous findings that the PCR marker *DXYCbl1*, whose sequence aligns at chrX: 169974684-169974964, is X-unique in offspring of (C57BL/6JxSPRET/Ei)F1 males (Breen *et al.* 1994; Perry *et al.* 2001). Given the conservation of copy number on either side of the PAR boundary in three outgroup species, we conclude that the duplications observed in each *M. musculus* subspecies are derived alleles. Without pedigree data we cannot be certain of X-, Y- or pseudoautosomal linkage for these duplications, but similarity to the pattern we observe for the PWK/PhJ allele – which itself segregates in the wild – strongly suggests that at least some involve the pseudoautosomal boundary.

## Discussion

Here we combine crosses involving inter-subspecific hybrid males with whole-genome sequencing data from wild mice to clarify and extend previous observations on the lability of the mouse pseudoautosomal region. We show that the previously-described extension of the PAR in *M. m. castaneus*-derived CAST/EiJ – that is, translocation of X-unique sequence onto the Y chromosome – undergoes crossing-over in heterozygous males at nearly 100-fold higher rate than autosomal sequence. Our estimate of the recombination rate in the extended PAR (18 - 23 cM/Mb) agrees with a recent estimate from whole-genome sequencing of single recombinant sperm (24 cM/Mb) from a C57BL/6J×CAST/EiJ cross (Hinch *et al.* 2019) male. The nominal marker order in the reference genome is supported by haplotypes reconstructed for recombinant progeny. Nonetheless, we observe several double-recombinants within < 500 kb, consistent with prior observations that double-recombinants are common in the canonical PAR (Soriano *et al.* 1987; Pardo-Manuel de Villena and Sapienza 1996).

We further confirm that a shorter extension of the PAR has occurred in PWK/PhJ (White *et al.* 2012a). Unlike the CAST/EiJ allele, the X- and Y-linked copies in PWK/PhJ are divergent in sequence and structure. This suggests that they do not recombine freely in the inbred strain, although we find that they do recombine in inter-subspecific hybrid males. The X-linked copy is of *M. m. domesticus* ancestry – evidence of either introgression in the wild or contamination in a laboratory colony – while the Y-linked copy comes from *M. m. musculus*, is segregating in the wild in the Czech Republic.

In fact the PWK/PhJ Y-linked allele is just one of numerous structural variants at the PAR boundary that segregate in natural populations of all three subspecies. All of these are derived alleles: both the PAR boundary and a proximal segment of the canonical PAR are single-copy in *M. spretus, M. spicilegus* and *M. caroli* and therefore most likely were single-copy in the common *Mus* ancestor. Using breakpoint-spanning reads to uniquely identify structural alleles, we find at least 32 alleles in a sample of 67 mice. None of them are shared across sub-species. The unusual level of standing variation suggests that the mutation rate near the PAR boundary is quite high. Indeed, we recover a *de novo* deletion on the PWK/PhJ Y haplotype in a sample of only 86 mice, for an estimated mutation rate of 1.2% per generation (95% Poisson CI 0.066% - 5.1%). To our surprise, none of these rearrangements seem to prevent robust expression of an essential gene (*Mid1*) that flanks the pseudoautosomal boundary (Dal Zotto *et al.* (1998) and Figure 8).

Taken together, our results indicate that the peculiar properties of the PAR in male meiosis – intense recombination activity, apparent absence of crossover interference, markedly elevated rates of structural rearrangement (Kipling *et al.* 1996a,b) – can “leak” across the PAR boundary. This is in spite of an abrupt and complete loss of sequence homology between the X and Y, at least in the reference assembly. Extension of the PAR requires transposition of sequence from the X onto the Y chromosome. Once this occurs, the region seems immediately subject to PAR-like levels of recombination despite not sharing the sequence features of the canonical PAR. It follows that these sequence features (such as elevated GC content) are indeed a consequence rather than a cause of the PAR’s high recombination rate in males. A summary of the major evolutionary events at the pseudoautosomal boundary in *Mus* is provided in Figure 10.

**Figure 10:**
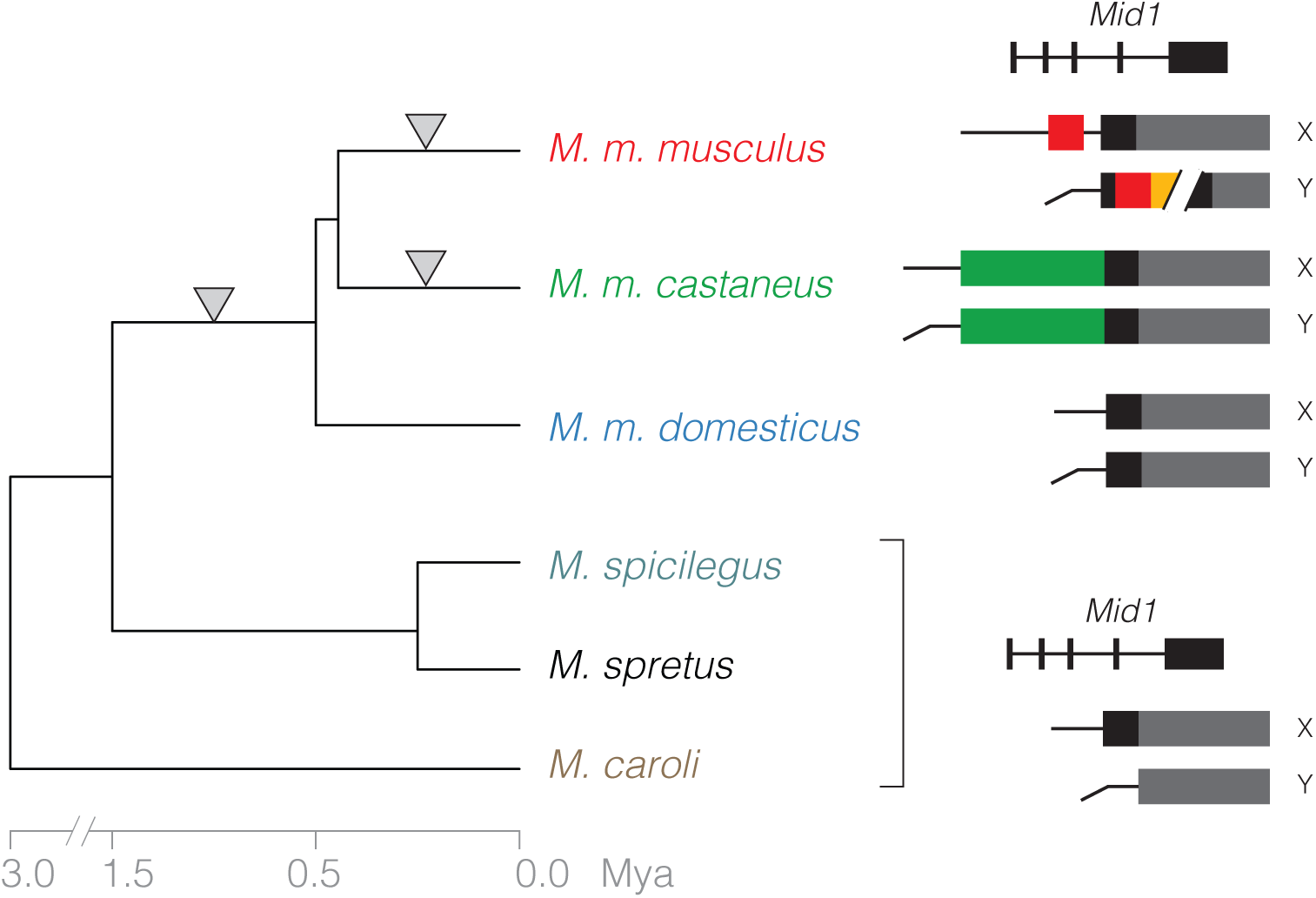
Changes in the pseudoautosomal boundary mapped onto the *Mus* phylogeny. Ancestral PAR, including unassembled repetitive regions, represented in grey. X-unique sequence translocated to the Y, expanding the PAR, shown as colored blocks. There are now at least three independent alterations of the boundary and/or organization of the PAR (inverted triangles): one in the ancestor of *M. musculus* (after the divergence from the *M. spretus*–*M. spicilegus* clade), expanding the PAR to include exons 4–8 of *Mid1*; the previously-documented expansion in CAST/EiJ; and the rearrangement we describe in PWK/PhJ. Approximate divergence times are shown below the tree, and schematics of the organization of each allele at right.

What could account for *de novo* X-Y exchanges in the absence of homology to promote pairing? One possibility is that the duplications happen through a mechanism such as non-homologous end joining. Another is ectopic recombination mediated by repetitive sequences such as transposon fragments (see Figure 5) or satellites. This is the mechanism for the human PAR1 extension present in some populations of European ancestry (Mensah *et al.* 2014). It has recently been shown that DSBs in the canonical PAR are formed preferentially at minisatellite arrays, and that it is the chromatin configuration of the repeat-rich sequence – rather than the sequence *per se* – that is essential for X-Y recombination (Acquaviva *et al.* 2019). It seems plausible that any nearby sequence drawn into the protein-DNA complex at the chromosomal axis could be susceptible to unequal exchange. Once present on both sex chromosomes, it would be prone to copy-number changes by further unequal exchange events.

Our findings support the notion that while the requirement for X-Y pairing in the first meiotic prophase is strictly enforced, the degree of structural homology in the PAR necessary to achieve pairing is relatively weak in mice. It has been known for several decades that failure to pair the sex chromosomes is associated with meiotic arrest in inter-specific (Matsuda *et al.* 1992; Hale *et al.* 1993) and inter-subspecific mouse hybrids (Forejt 1996). The unpaired sex chromosomes do not undergo meiotic sex-chromosome inactivation, and misexpression of X- and Y-linked genes is apparently toxic to germ cells (Good *et al.* 2010; Vernet *et al.* 2011; Campbell *et al.* 2013). However, absence of X-Y pairing in sterile hybrids occurs in the context of widespread asynapsis of both autosomes and sex chromosomes (Bhattacharyya *et al.* 2013). More direct evidence of the requirement for recombination in the PAR comes from comparison of genetically-engineered mice deficient for isoforms of *Spo11*, which encodes the enzyme that executes programmed meiotic DSBs (Keeney *et al.* 1997). The SPO11-*β* isoform is required for DSB formation and synapsis on autosomes and (in females) the X-chromosome pair; SPO11-*α* is required for DSB formation in the PAR (Bellani *et al.* 2010). Male mice deficient in SPO11-*α* alone have high rates of X-Y asynapsis and are sterile or subfertile, depending on genetic background, despite having normal crossover frequency and synapsis on autosomes (Kauppi *et al.* 2011; Faisal and Kauppi 2016). Faisal and Kauppi (2016) speculate that partial rescue of the SPO11-*α* knockout may be due to a PAR variant in segregating in their colony. The best evidence for a specific role of PAR divergence in infertility comes from an experiment by White *et al.* (2012b): F2 males heterozygous for a WSB/EiJ and a CAST/EiJ haplotype in the PAR have a variety of spermatogenic defects, independent of genotypes at other loci.

However, Dumont (2017) has shown that although the number of X-Y crossovers (assayed by immunostaining against the MLH1 protein) in inter-subspecific F1 hybrid males is reduced relative to the parental strains, absence of stable X-Y pairing is pervasive in both inbreds and F1s. Failure to pair the sex chromosomes is associated with meiotic arrest and subsequent apoptosis of up to 80% of spermatocytes in some inter-subspecific hybrids (Matsuda *et al.* 1992) – but also up to 25% of spermatocytes in some inbred strains (Imai *et al.* 1981; Krzanowska 1989). Even this level of X-Y dissociation is compatible with fertility, at least under laboratory conditions. Two Y-chromosome variants with large extensions of the PAR – the *Yaa* allele, a 4 Mb proximal extension of the PAR on the SB/Le background (Hudgins *et al.* 1985; Subramanian *et al.* 2006); and the *Y\** allele, a PAR-to-PAR fusion of a Y and an X chromosome (Eicher *et al.* 1991) – are readily propagated in laboratory colonies. Given these observations and the number of segregating PAR-boundary alleles we observe in natural populations, it is reasonable to conclude that the organization of the region is not under strong purifying selection in house mice. However, we stress that our study provides limited direct information on the fitness consequences of X-Y exchanges in the PAR in inter-subspecific hybrids. These events are compatible with viability through at least the neonatal period (when progeny were sacrificed for genotyping) but details of their effects on male and female fertility are unknown.

Our study has several further limitations. First, all of our analyses – whether of array genotypes or whole-genome sequence data – are predicated on the representation of the PAR in the mm10 reference genome. The assembly of the PAR has proven difficult because of its repetitive content, and is thus fragmented and incomplete. The PAR of C57BL/6J is approximately 750 kb long by restriction mapping (Perry *et al.* 2001) but has only 161 kb of non-gap bases in mm10. We are blind to any sequence that does not have at least partial representation in the reference assembly. It is extremely difficult, and in many cases impossible, to accurately reconstruct complex structural variants from short-read data alone. The noisy alignment patterns in the canonical PAR strongly suggest that its organization is much more complex than the reference assembly would suggest, but we can say little more than that. Detailed characterization of repetitive loci in mammalian genomes (eg. Soh *et al.* (2014); Cantsilieris *et al.* (2018); Lilue *et al.* (2018)) with orthogonal technologies including long-read single-molecule sequencing and high-throughput physical mapping has shown that the view from short reads may be a faint shadow of the truth. Finally, despite these limitations, our study provides a useful foundation for future efforts to characterize the sequence, structure and function of this enigmatic region of the genome.

## Materials and Methods

### Mouse breeding

The crosses described herein were bred as part of ongoing studies of aging, recombination and infertility in hybrid males. Standard mouse strains (CAST/EiJ, PWK/PhJ, WSB/EiJ, FVB/NJ) were purchased from the Jackson Laboratory (Bar Harbor, ME) between 2008 and 2011. Reciprocal F1 hybrid males were generated in all pairwise combinations between CAST/EiJ (abbreviated with strain code F) PWK/PhJ (G), and WSB/EiJ (H) in 2008-2009. F1 males were bred with multiple FVB/NJ females to generate N2 progeny which were sacrificed at birth by cervical dislocation. N2 progeny were named as follows: sire identifier (PAE_xxx), 2-letter encoding of sire genotype (eg. HF for WSB/EiJ×CAST/EiJ), 1-letter encoding of approximate sire age at conception (E=elderly, Y=young), and 2- or 3-letter pup identifier. All breeding was completed by 2011. The study was approved by the Institutional Animal Care and Use Committee of the University of North Carolina at Chapel Hill and all animal husbandry was conducted in the university’s AAALAC-accredited facility (Animal Welfare Assurance #A3410-01) accordance with institutional and federal regulations.

### DNA extraction, genotyping and quality control

Genomic DNA was extracted from tail clips of adult representatives of parental inbred strains or whole tails or heads of N2 progeny using either a standard phenol-chloroform method (Green and Sambrook 2012) or Qiagen DNeasy Blood & Tissue Kits (catalog no. 69506; Qiagen, Hilden, Germany). Approximately 1.5 micrograms of DNA per sample was shipped to Neogen Inc (Lincoln, NE) for hybridization to one of two Mouse Universal Genotyping Arrays, MegaMUGA (77K probes) or MiniMUGA (11K probes). Both arrays are custom-designed on the Illumina Infinium HD platform (Steemers *et al.* 2006), with content optimized for information content in classical inbred strains and the 8 founder strains of the Collaborative Cross (A/J, C57BL/6J, 129S1/SvImJ, NOD/ShiLtJ, NZO/HlLtJ, CAST/EiJ, PWK/PhJ, WSB/EiJ), as described in (Morgan *et al.* 2016). Genotypes were called by the vendor from hybridization intensity signals using the semi-supervised clustering algorithm implemented in the Illumina BeadStudio software (Illumina Inc, San Diego, CA).

Genotype data was processed in `R` v3.3.2 (R Foundation for Statistical Computing, http://www.r-project.org/) using the `argyle` package v0.2.2 (Morgan 2016). Samples with < 10% missing genotype calls were retained for analysis; there were no genotyping failures. Samples were genetically sexed by comparing average hybridization intensities for X- and Y-linked markers. We identified a single XO individual among the PWK/PhJ×CAST/EiJ N2s and none in the WSB/EiJ×CAST/EiJ or WSB/EiJ×PWK/PhJ N2s. To confirm correct assignment of N2s to F1 sires, we examined N2 genotypes at sites on the X chromosome informative between the parent strains. We identified 24 progeny representing 2 consecutive litters from the same sire (PAE_018) initially labelled as WSB/EiJ×CAST/EiJ. Males were homozygous for WSB/EiJ genotypes on the X and females heterozygous at sites informative between CAST/EiJ and FVB/NJ. These pattern was consistent with a (CAST/EiJxWSB/EiJ)F1 sire, and we updated our pedigrees accordingly.

A complete list of N2 progeny, their parents, genetically-confirmed sex, and platform on which they were genotyped is provided in File S4.

### Refinement of genotypes on the X chromosome

Because the aim of this study is to identify crossovers in the PAR, we focused particular attention on the quality of genotype calls at markers in this region. The extended PAR of CAST/EiJ begins (in the mm10 assembly) at chrX:169542082, and the canonical PAR at chrX:169969759. We augmented MegaMUGA genotypes and hybridization intensitnes (*n* = 186 N2s +4 parental strains) with 111 representatives of the 8 Collaborative Cross founder strains and the 26 viable reciprocal F1 hybrids between them. (These control genotypes are available for public download from http://csbio.unc.edu/CCstatus/index.py.) Of 19 markers whose probe sequence aligns to the PAR in mm10, only one (JAX00723372) yields automated calls in the heterozygous and both homozygous states among the 111 control samples. We manually inspected hybridization intensities at each of the remaining 18 markers to find any with cluster patterns from which reliable genotypes might be recovered. Three clusters – putatively corresponding to the two homozygous and one heterozygous states – were observed for at a single marker, UNC31596457. No heterozygotes are called at this marker by the automated algorithm in BeadStudio. However, it is clear that CAST/EiJ has a private allele at this marker, and that one of the clusters corresponds to CAST/(non-CAST) heterozygotes in the control samples (Figure S7). We therefore re-called genotypes at this marker by fitting a 3-component bivariate Gaussian mixture model to hybridization intensities of the control samples and N2s using the `R` package `mclust` v5.2.1 (Fraley and Raftery 2002). Samples with cluster assignment probability < 0.9 were marked as missing. The result is shown in Figure S8.

Following this genotype refinement for MegaMUGA, informative markers in or beyond the extended PAR of CAST/EiJ were identified separately on the two array platforms by comparing genotypes between single representatives of the relevant inbred strains. This left 7 and 6 markers on MegaMUGA for the PWK/PhJ×CAST/EiJ and WSB/EiJ×CAST/EiJ crosses respectively; and 1 and 2 markers on MiniMUGA for the same crosses. A list of informative markers for each array-cross combination is provided in File S2. Refined genotype calls for all N2 progeny and controls in the study are provided in File S5.

Crossovers in the PAR were identified as illustrated in Figure 2B. Genetic lengths of inter-marker intervals were calculated assuming an identity map function (ie. no crossover interference) and standard errors obtained from formula 2.12 in (Xu 2013).

### Whole-genome sequencing

New whole-genome sequence data was generated for 87 N2 progeny of the WSB/EiJ×PWK/PhJ cross at the University of North Carolina High-Throughput Sequencing Facility. Genomic DNA was fragmented by ultrasonication on a Covaris instrument (Covaris Inc, Woburn, MA) and enriched for fragments approximately 300 bp in size on a PippinPrep system (Sage Science, Beverly, MA). Paired-end libraries were prepared using the Illumina TruSeq PE v1 chemistry. Samples were multiplexed in two pools (with 45 and 42 samples, respectively) and sequenced 2×100bp on an Illumina HiSeq 2500 instrument to a depth of approximately 5×. Demultiplexing and postprocessing were performed with Illumina CASAVA v1.8.2. Data has been submitted to the European Nucleotide Archive (accession PRJEB32247).

Reads for inbred strains PWK/PhJ, FVB/NJ and WSB/EiJ was obtained from the public FTP site of the Sanger Mouse Genomes Project, February 2015 release (ftp://ftp-mouse.sanger.ac.uk/REL-1502-BAM) (Keane *et al.* 2011). Reads for Collaborative Cross strains CC001/Unc, CC022/GeniUnc, CC032/GeniUnc, CC044/GeniUNC and CC045/GeniUnc were generated by our research group and have been previously published (European Nucleotide Archive accession PRJEB14673) (Srivastava *et al.* 2017). Reads for wild *M. musculus* and *M. spretus* have been previously published (Harr *et al.* 2016) and were obtained from the European Nucleotide Archive (accessions PRJEB9450, PRJEB11742, PRJEB14167, PRJEB2176). Reads for *M. spicilegus* have been previously published (Neme and Tautz 2016) and were obtained from the European Nucleotide Archive (accession PRJEB11513). Reads for *M. caroli* have been previously published (Thybert *et al.* 2018) and were obtained from the European Nucleotide Archive (accession PRJEB14895). A complete list of samples used in whole-genome sequencing analysis is provided in File S6.

The standard mouse reference assembly is a haploid representation of the genome – except in the PAR, whose sequence appears on both sex chromosomes. This induces ambiguous sequence alignments and effectively halves the observed read depth over each copy of the PAR. To avoid these artifacts, we masked the sequence of the Y PAR (chrY:90745845-91644698) in the mm10/GRCm38 assembly with Ns. Reads were aligned to this masked reference with `bwa mem` 0.7.15-r1140 (Li 2013) with flags `-t 16 -YK100000000` and default settings otherwise. Optical duplicates were marked with `samblaster` v0.1.24 (Faust and Hall 2014).

### Ascertainment of CNVs and their boundaries

Copy-number variants in the distal X chromosome were identified by examining read depth in windows of varying sizes, as described in the main text. Paired-end reads are expected to align with a characteristic orientation: one read on the forward strand and its mate on the reverse (configuration “FR”). Clusters of pairs with other orientations (RR, FF or RF) may be a signal of underlying structural variation. Summaries of total and FR read depth were calculated with `pysamstats` v1.1.0 (https://github.com/alimanfoo/pysamstats) and normalized against read depth in single-copy autosomal sequence. CNV boundaries were refined by manual inspection of read alignments in `IGV` v2.4.16 (Robinson *et al.* 2011). Clusters of 3 or more reads with soft-clipped alignments (that is, alignments of less than the full read length, indicated by the presence of `S` in the CIGAR string) all terminating at exactly the same base pair, and coinciding with an abrupt change in both total read depth and the proportion of FR reads, were deemed candidate CNV breakpoints. Such reads, as well as their mates (whether the mate was mapped or not) were extracted and converted to fastq format with `samtools` v1.9 (Li *et al.* 2009). For the breakpoints of the PWK/PhJ Y CNV, reads crossing each of 3 candidate breakpoints were assembled *de novo* with `SPAdes` v3.10.1 (Bankevich *et al.* 2012) with flags `--12` (for paired-end reads) and `--only-assembler` (skipping error correction, which requires higher coverage). The longest contig assembled for each breakpoint was aligned back to the mm10 reference with `blat` (Kent 2002) at the UCSC Genome Browser (https://genome.ucsc.edu/cgi-bin/hgTracks?db=mm10). Contigs were screened for possible open reading frames with the ExPASy Translate tool (https://web.expasy.org/translate/). Their sequences are provided in File S3.

To identify the source of breakpoint sequence in region_3 (Figure 5), we performed `blat` searches against recent *de novo* assemblies of *M. spretus* (GenBank accession GCA_001624865.1; (Lilue *et al.* 2018)), *M. caroli* (GCA_900094665.2; (Thybert *et al.* 2018)) and *M. pahari* (GCA_900095145.2; (Thybert *et al.* 2018)). This contig has gapped alignments to these outgroups genomes that resemble alignments of a spliced transcript (Figure S9, File S7), but the aligned segments do not correspond to annotated exons. The longest ORF it encodes is 66 aa long, and `BLAST` searches of 6-frame translations (`blastx`) against the NCBI non-redundant protein database yield no hits with *e*-value < 1 (search performed 31 January 2019). We conclude that despite having the appearance of a retrotransposition event, the sequence is unlikely to represent an ancestral coding sequence.

### SNV calling and filtering

For analyses of recombination within the PWK/PhJ extended PAR, we used bcftools v1.9 to ascertain biallelic SNVs in the distal X chromosome (chrX:169400000-169969759) with call rate > 90% in 169 samples including N2 progeny, inbred parental strains, wild mice, and representative CC strains with a PWK/PhJ Y chromosome. We retained 84 sites homozygous in the PWK/PhJ female and heterozygous in the CC044/Unc strain (carrying a PWK/PhJ Y chromosome) and segregating at absolute allele count > 10 in the N2s. A single N2 individual (PAE_033_HG_E_463) had an outlying number of heterozygous calls and appeared to be cross-contaminated with another individual; it was excluded from this and all other analyses. Genotype calls at these sites are provided in VCF format in File S8.

Principal components analysis (PCA) on genotypes in the distal X chromosome was performed with `akt` v3beb346 (Arthur *et al.* 2017), treating all samples as nominally diploid (ie. substituting homozygous for hemizygous genotypes.) Because we did not have access to whole-genome sequence data for an inbred PWK/PhJ male, we created a pseudo-diploid individual homozygous for PWK/PhJ Y-linked alleles in order to perform ancestry assignment for the PWK/PhJ Y PAR.

### Re-analysis of ChIPseq data

Raw reads from published ChIPseq against H3K4me3 in 12 days post-partum testes from reciprocal F1 hybrid males between C57BL/6J and CAST/EiJ (Baker *et al.* 2015) were downloaded from the NCBI Short Read Archive (accession PRJNA259788) and aligned to the mm10 reference with bwa mem as described above. Post-processed read alignments from ChIPseq against DMC1 from the same genotypes (but different laboratories), representing single-stranded DNA fragments at DSB free ends (Smagulova *et al.* 2016), were downloaded from NCBI GEO (accession GSE75419).

Tag reads were assigned to the CAST/EiJ or C57BL/6J haplotype according to the base called at known informative sites between these strains (using biallelic SNVs from the Mouse Genomes Project VCF, May 2015 release; ftp://ftp-mouse.sanger.ac.uk/REL-1505-SNPs_Indels/). Given *n* informative sites spanned by a read, the log-likelihood of the read originating from each parental haplotype was calculated as the binomial probability of observing *k* bases consistent with that haplotype and *n - k* bases inconsistent. A read was deemed assignable if log-likelihood of maternal versus paternal assignments differed by > 2 units. Tag density was calculated with the `bamCoverage` tool from the `bamTools` suite (Barnett *et al.* 2011) using 100 bp bins and a 500 bp smoothing window. No ChIP background subtraction or peak-calling was performed.

We note that, since reads were aligned to the standard mm10 reference, reads arising from the CAST/EiJ haplotype are somewhat less likely to be aligned, and so the count of CAST/EiJ reads will be an underestimate. This “reference bias” causes our analyses to be conservative.

### Analysis of gene expression

Assay of gene expression by paired-end, unstranded RNAseq in adult brain from 90 progeny of a complete diallel cross (eg. all possible reciprocal crosses) between CAST/EiJ, PWK/PhJ and WSB/EiJ has been described previously (Crowley *et al.* 2015). We obtained the raw reads and quantified expression using `kallisto` v0.44.0 (Bray *et al.* 2016) with default options. Similar to the approach described for whole-genome sequencing, we removed 8 redundant transcripts annotated in the Y-encoded copy of the PAR (retaining X-encoded copies) and performed quantification against the remaining 141, 854 transcripts in the Ensembl version 96 annotation. Transcript-level estimates were aggregated to gene level with the `R` package `tximport` (Soneson *et al.* 2015) and analyses of differential expression were performed with `limma-voom` (Ritchie *et al.* 2015).

To investigate allele-specific expression and splicing patterns we also aligned reads to our masked version of the reference genome and masked transcript annotation with `STAR` v2.7.0e (Dobin *et al.* 2013). Read pairs were aligned to parental haplotypes as described above for the ChIPseq data. Coverage along annotated exons of *Mid1* was calculated with `bamCoverage` using 5 bp bins and a 25 bp smoothing window. Reads were pooled across biological replicates of the same genotype and sex.

*Mid1* transcript isoforms were assembled from read alignments in the parental strains (CAST/EiJ, PWK/PhJ and WSB/EiJ), pooling reads from both male and female samples, using `StringTie` v1.3.5 (Pertea *et al.* 2015) with options `-f 0.1 -a 4 -c 0.1 -g 500` and the Ensembl version 96 annotation as a guide.

### Data availability

Whole-genome sequence data from N2 progeny has been submitted to the European Nucleotide Archive (accession PRJEB32247). Supplementary files have been submitted to FigShare. Inferred genetic map for the extended PAR is in File S1; informative markers near PAR boundary in File S2; assembled breakpoint sequences of PWK/PhJ PAR variant in File S3; list of N2 progeny used for recombination analysis in File S4; array genotype and intensity data, with marker locations, in File S5; and genotype calls in PWK/PhJ PAR variant from whole-genome sequence in File S8. Supporting code is available at https://github.com/andrewparkermorgan/mouse_pseudoautosomal_region.

## Acknowledgments

The authors thank Grace Clark, Justin Gooch, the late Mark Calaway, and Darla Miller for assistance with DNA preparation and submission of samples for genotyping; Randy Nonneman for assistance with mouse husbandry; and Chris Baker, Kevin Brick and Fatima Smagulova for their generous technical assistance in reanalyzing their published ChIPseq data. The authors benefited from fruitful correspondence with Beth Dumont while preparing the manuscript. This work was supported in part by National Institutes of Health grants F30MH103925 (APM), R01HD065024 (FPMdV), R21MH096261 (FPMdV), K01MH094406 (JJC), P50MH090338 (FPMdV), P50HG006582 (FPMdV) and U42OD010924 (Terry Magnuson).

## Supplementary material

### Tables and data files

**Figure S1:**
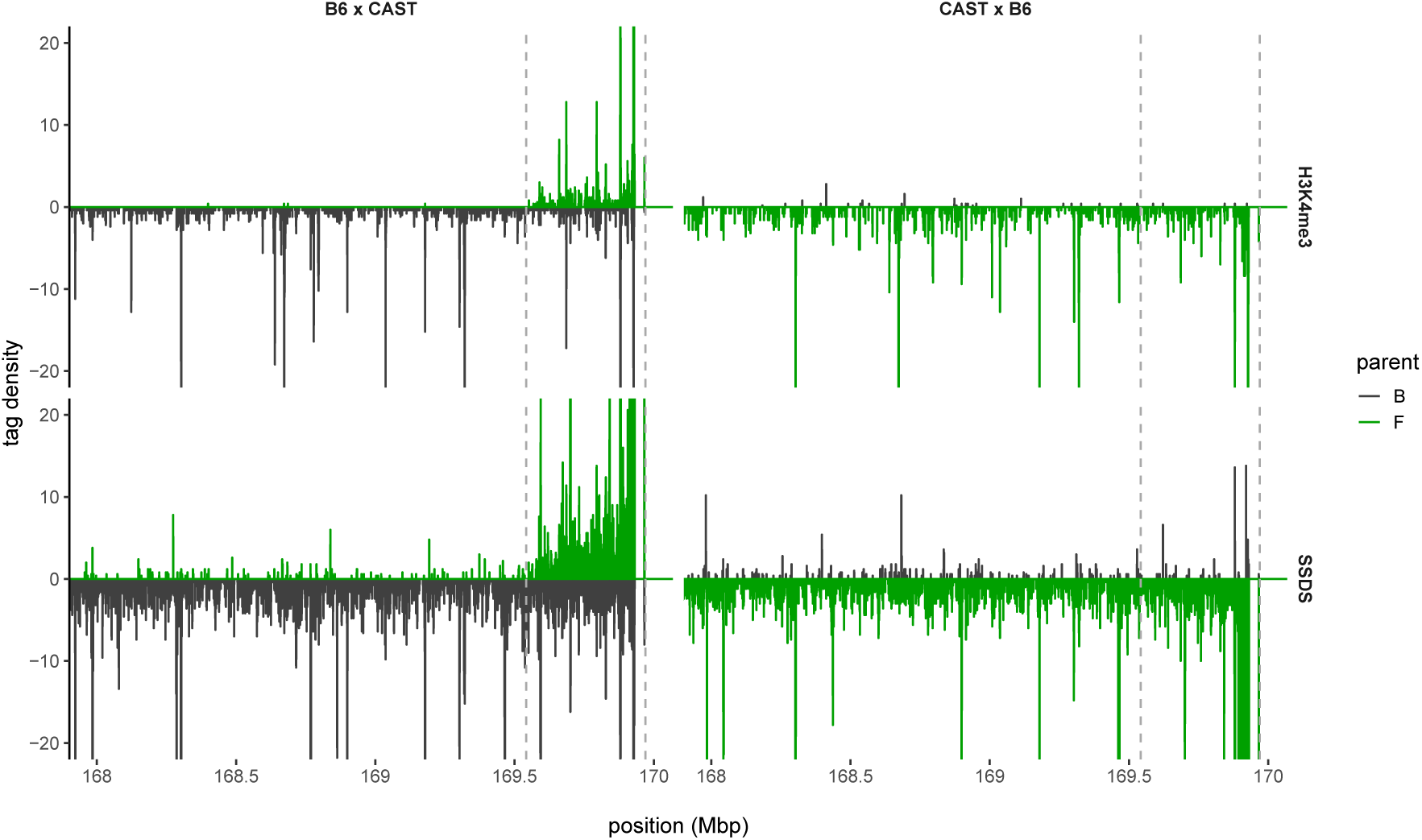
Accumulation of meiotic recombination precursors on the extended PAR of CAST/EiJ. Raw density of tags recovered from immunoprecipitation of H3K4me3 (top panels) or DMC1-bound single-stranded DNA fragments (bottom panels) in reciprocal hybrid males between C57BL/6J (B6, black) and CAST/EiJ (CAST, green). Reads were assigned to a parental haplotype on the basis of base calls at informative SNVs; signal on the paternal or maternal haplotype is plotted above or below the *x*-axis, respectively. Dashed lines indicate boundaries of the CAST/EiJ and canonical PARs.

**Figure S2:**
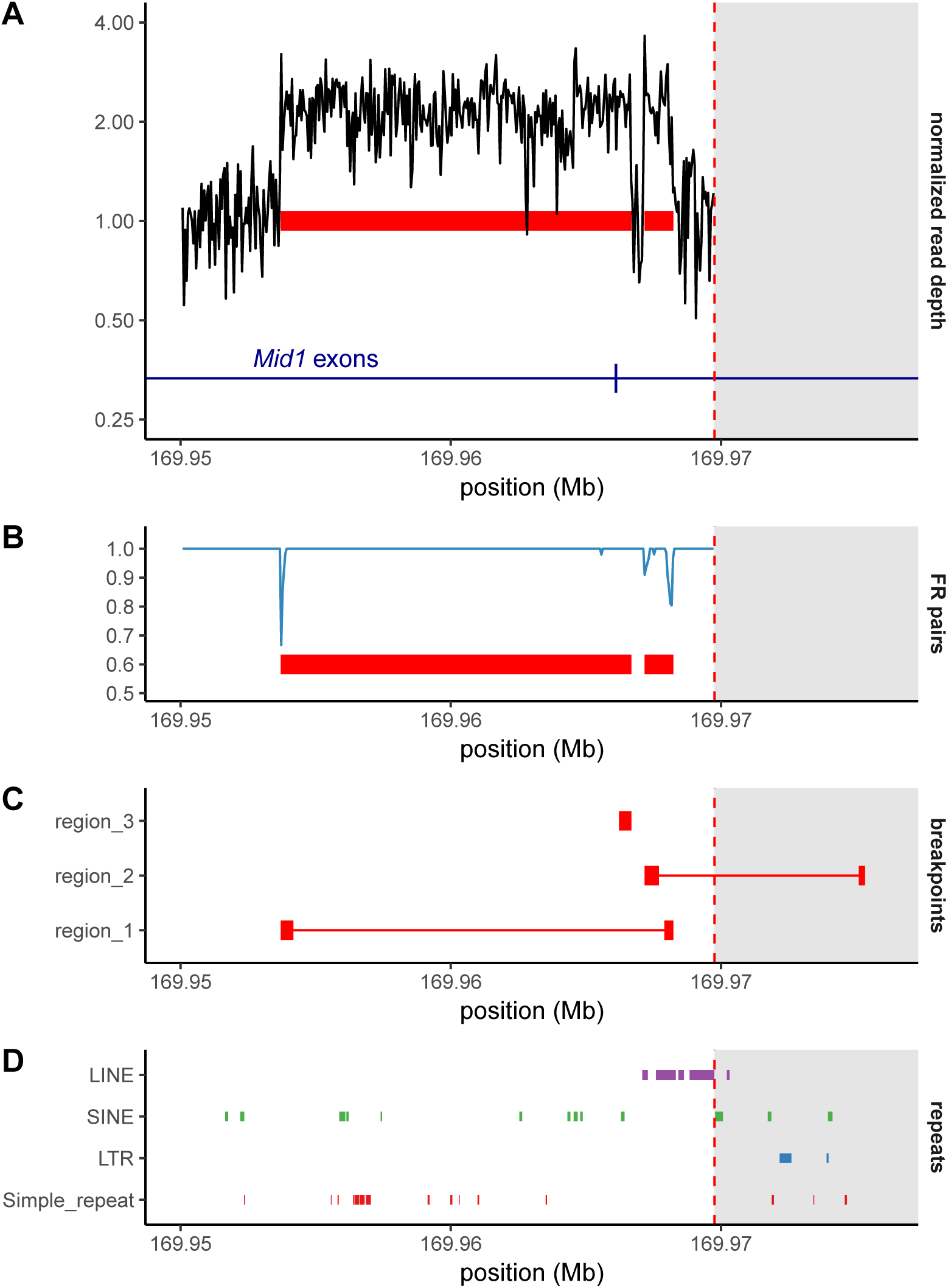
Detailed view of the Y-linked duplication identified in PWK/PhJ. (**A**) Copy number in the distal X chromosome, estimated from read depth in whole-genome sequencing in 200 bp bins and averaged over 4 individuals carrying a PWK/PhJ Y chromosome. Duplicated segments identified by red bars. Red dashed line indicates canonical PAR boundary. Copy number in the canonical PAR is not shown because of noisy read alignment to repetitive sequence. (**B**) Proportion of read pairs aligning in the expected orientation (FR), binned and average as above. (**C**) Alignments of *de novo* assembled contigs spanning duplication breakpoints. Segments of gapped alignments are connected with red bars. (**D**) Selected classes of repetitive elements annotated in the mm10 reference sequence.

**Figure S3:**
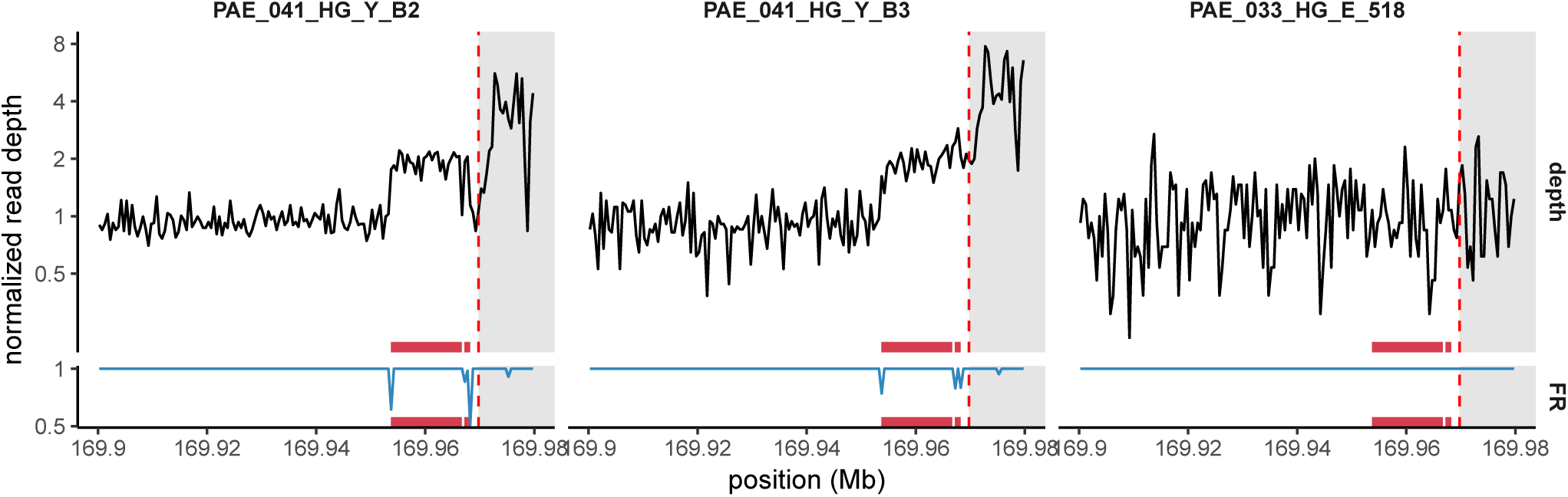
Discordant mate-pair orientation of reads at the boundary of the PWK/PhJ Y-linked duplication. Copy number (black) and proportion of read pairs aligning in the expected orientation (blue) are shown for 3 cases among N2 males: a non-recombinant PAR (PAE_041_HG_Y_B2); a recombinant PAR (PAE_041_HG_Y_B3); and the individual with *de novo* copy-number loss (PAE_041_HG_E_518). The *de novo* deletion is accompanied by loss of the discordant read pairs.

**Figure S4:**
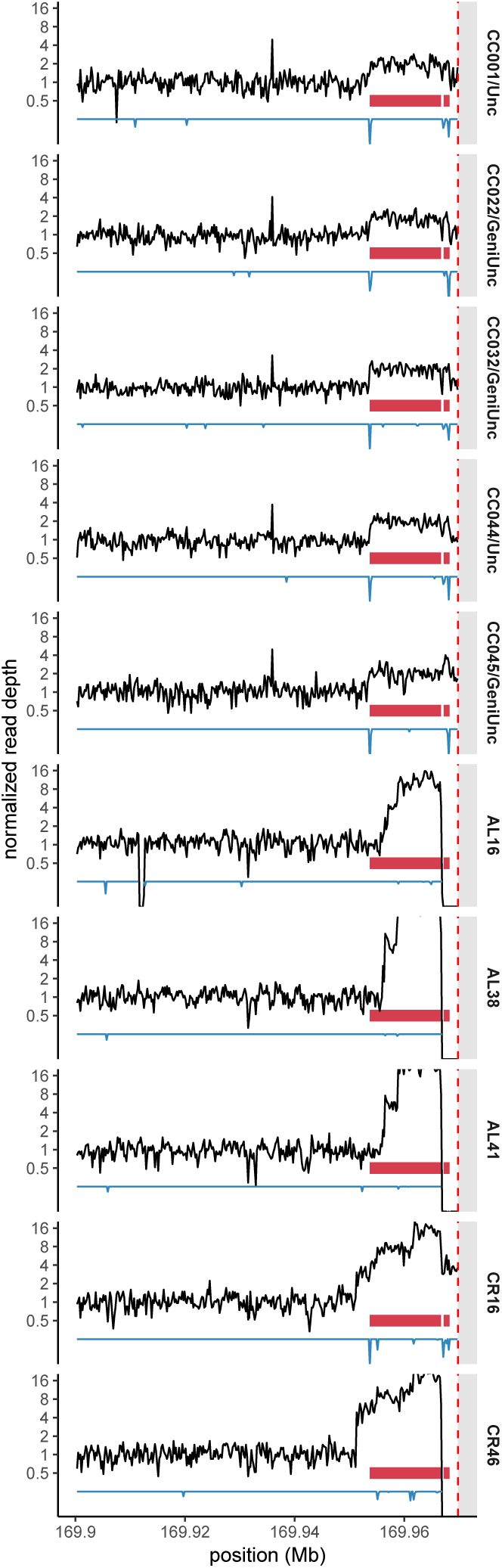
Identifying wild mice sharing the PWK/PhJ Y-linked duplication. Copy number in the distal X chromosome, estimated from read depth in whole-genome sequencing in 200 bp bins, for males from 5 Collaborative Cross strains with a PWK/PhJ Y chromosome (CCxxx), 2 wild males from Kazakhstan (ALxx) and 2 wild males from the Czech Republic (CRxx). Red dashed line shows position of the canonical PAR boundary. Dark red blocks identify the segment duplicated in PWK/PhJ. Blue lines show proportion of read pairs aligning in the expected orientation on an arbitrary scale.

[*provided as separate file*]

Figure S5: Normalized read depth on distal X chromosome in mice analyzed by whole-genome sequencing in this study. One individual per page, two panels per individual (one at relative broad scale, chrX:168-170 Mb; the other zoomed-in on PAR boundary.) Page title indicates sample type (wild-caught or wild-derived inbred strain), nominal subspecies of origin, country of origin (as ISO 2-letter code), sample ID and chromosomal sex.

**Figure S6:**
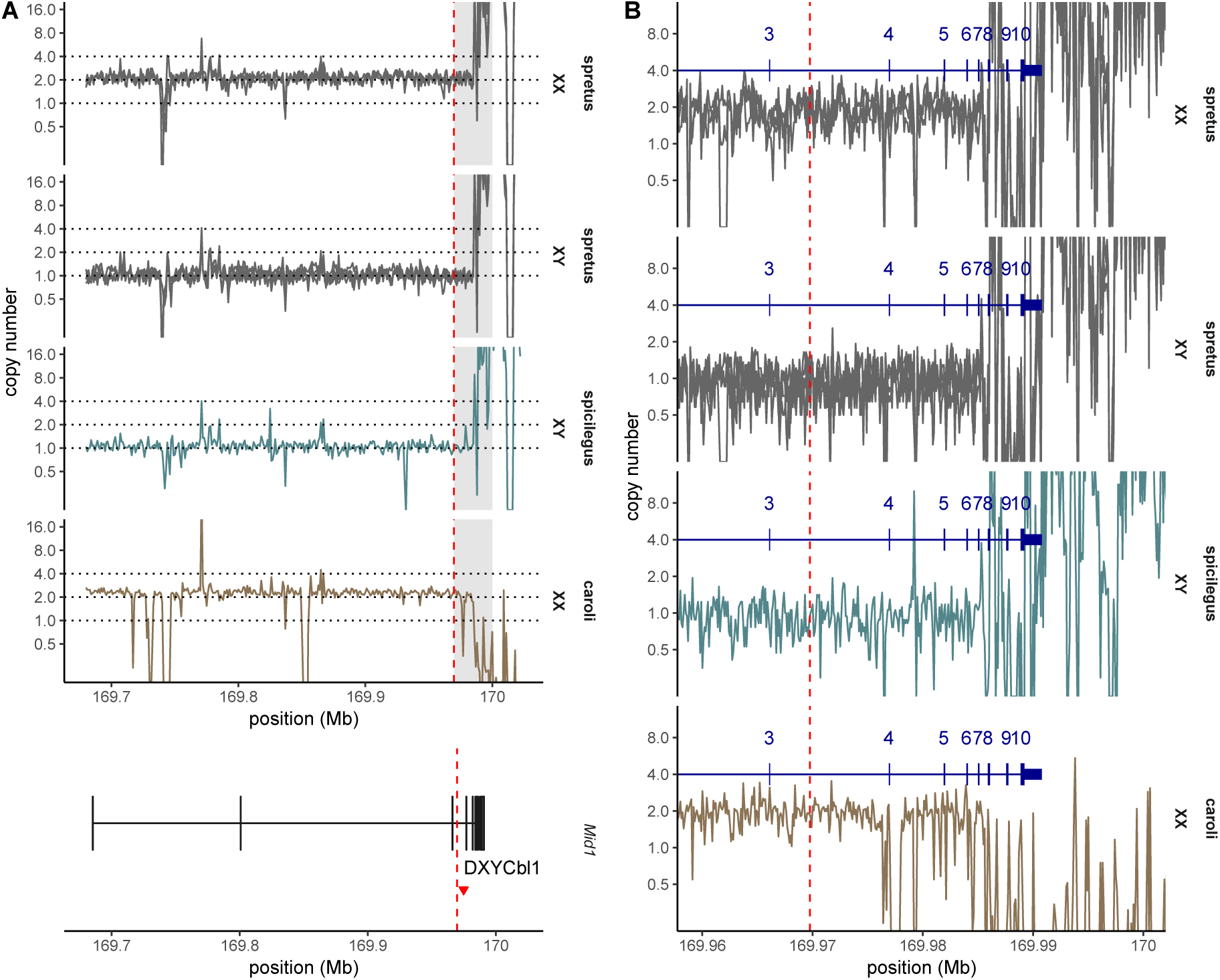
Defining the ancestral pseudoautosomal boundary in the common *Mus* ancestor. (**A**) Copy number, estimated from whole-genome sequencing read depth in 1 kb bins, in males and females from *M. spretus, M. spicilegus* and *M. caroli*. Red dashed line shows nominal PAR boundary in C57BL/6J with relation to exons of *Mid1*. Red triangle shows position of STS marker *DXYCbl1*. (**B**) Magnified view of the region just distal to the nominal PAR boundary. Read depth calculated in 100 bp windows. Exons of the longest coding transcript of *Mid1* are numbered in dark blue.

**Figure S7:**
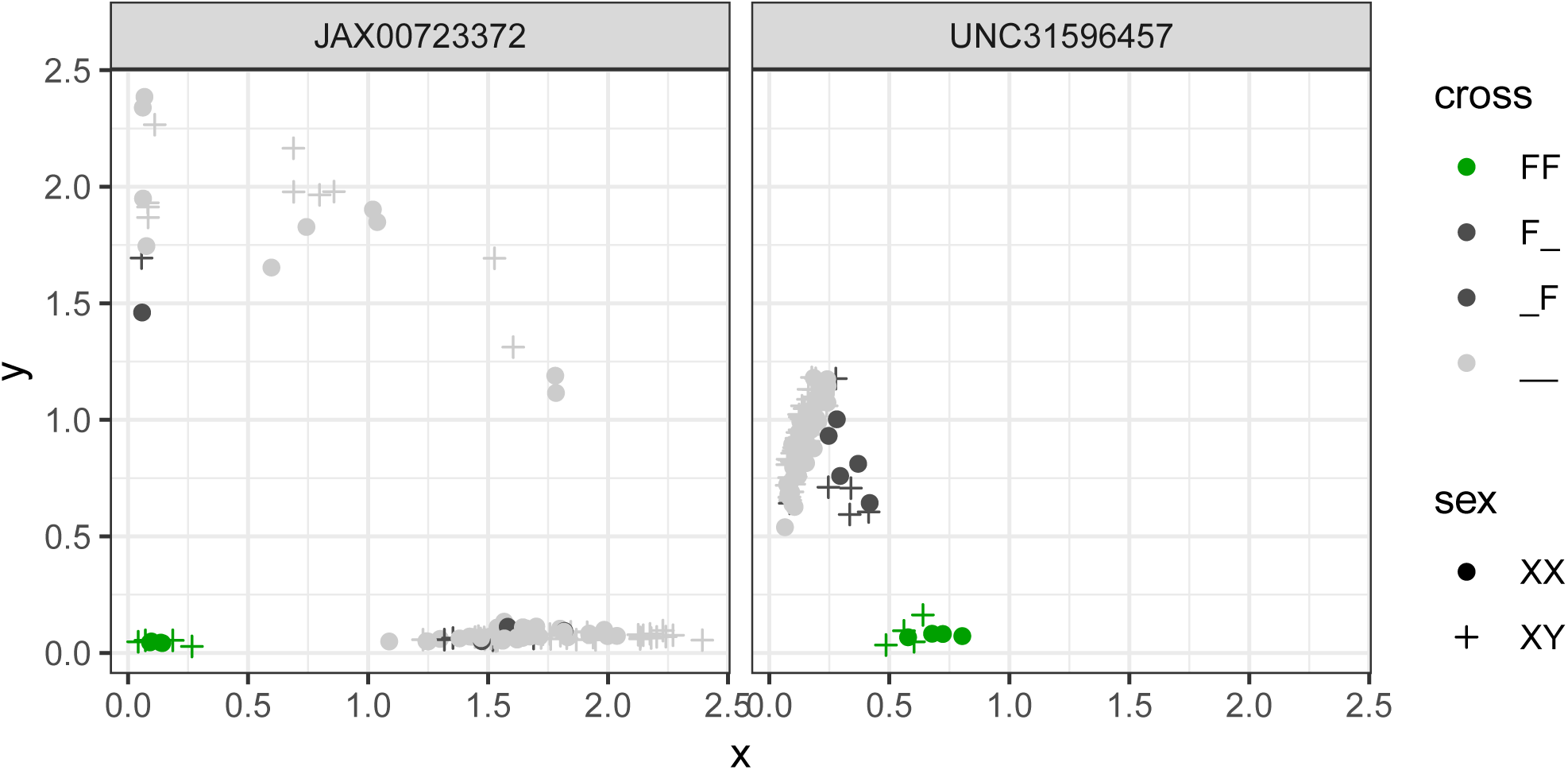
Confirming consistency of genotype calls at two markers in the canonical PAR. Hybridization intensities in Cartesian coordinates for 111 inbred and F1 controls are shown, colored by genotype: FF = CASTEi-JxCAST/EiJ; F_ = CAST/EiJ x other; _F = other x CAST/EiJ; = other x other. Shapes distinguish males and females.

**Figure S8:**
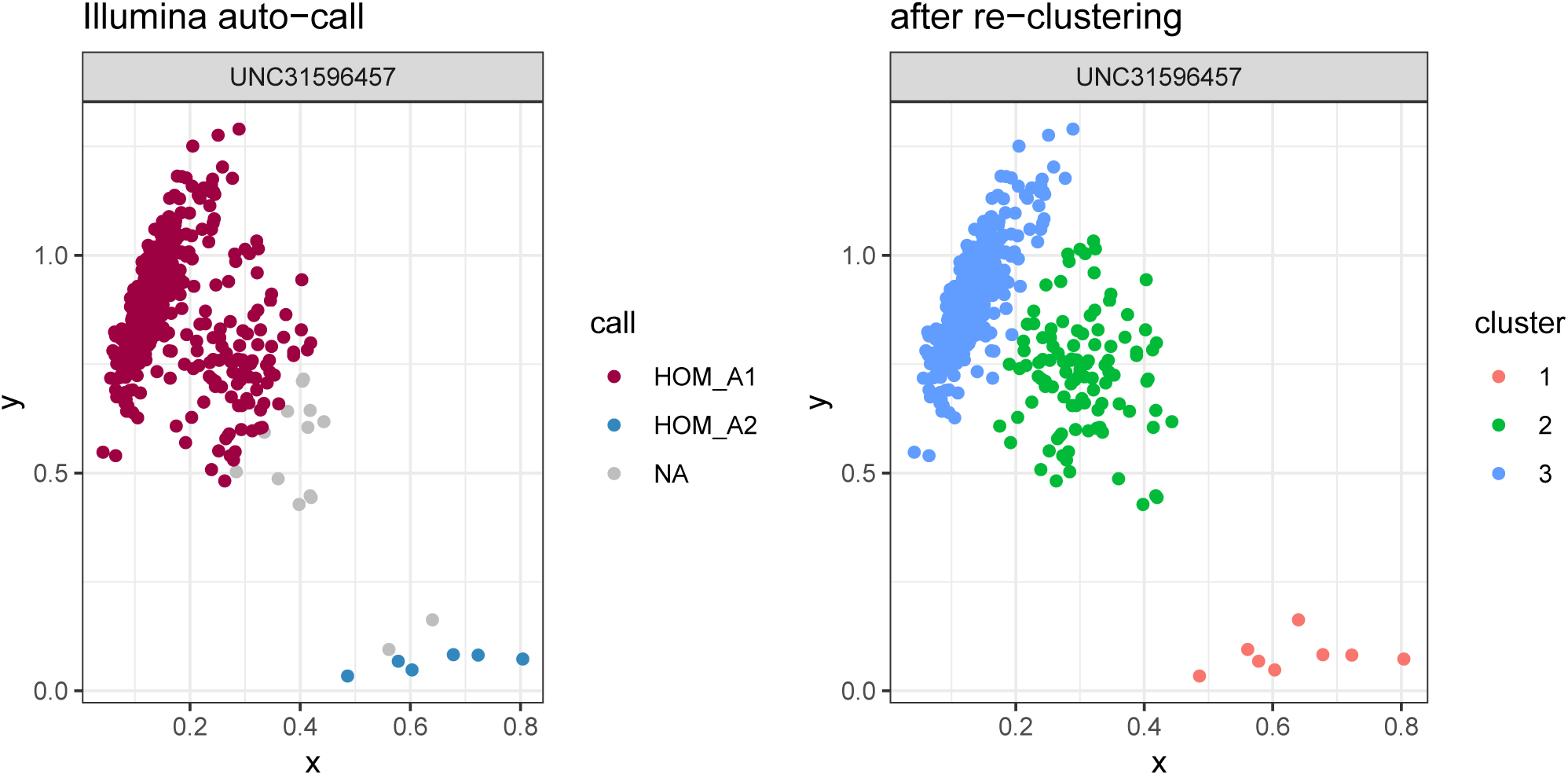
Revision of genotype calls at array marker UNC31596457. Hybridization intensities in Cartesian coordinates for 111 inbred and F1 controls and 186 N2 progeny are shown, colored by genotype call from Illumina BeadStudio (left) or our Gaussian mixture model (right). Clusters 1 and 3 correspond to non-CAST/EiJ and CAST/EiJ homozygotes, respectively, with heterozygotes in cluster 2.

**Figure S9:**
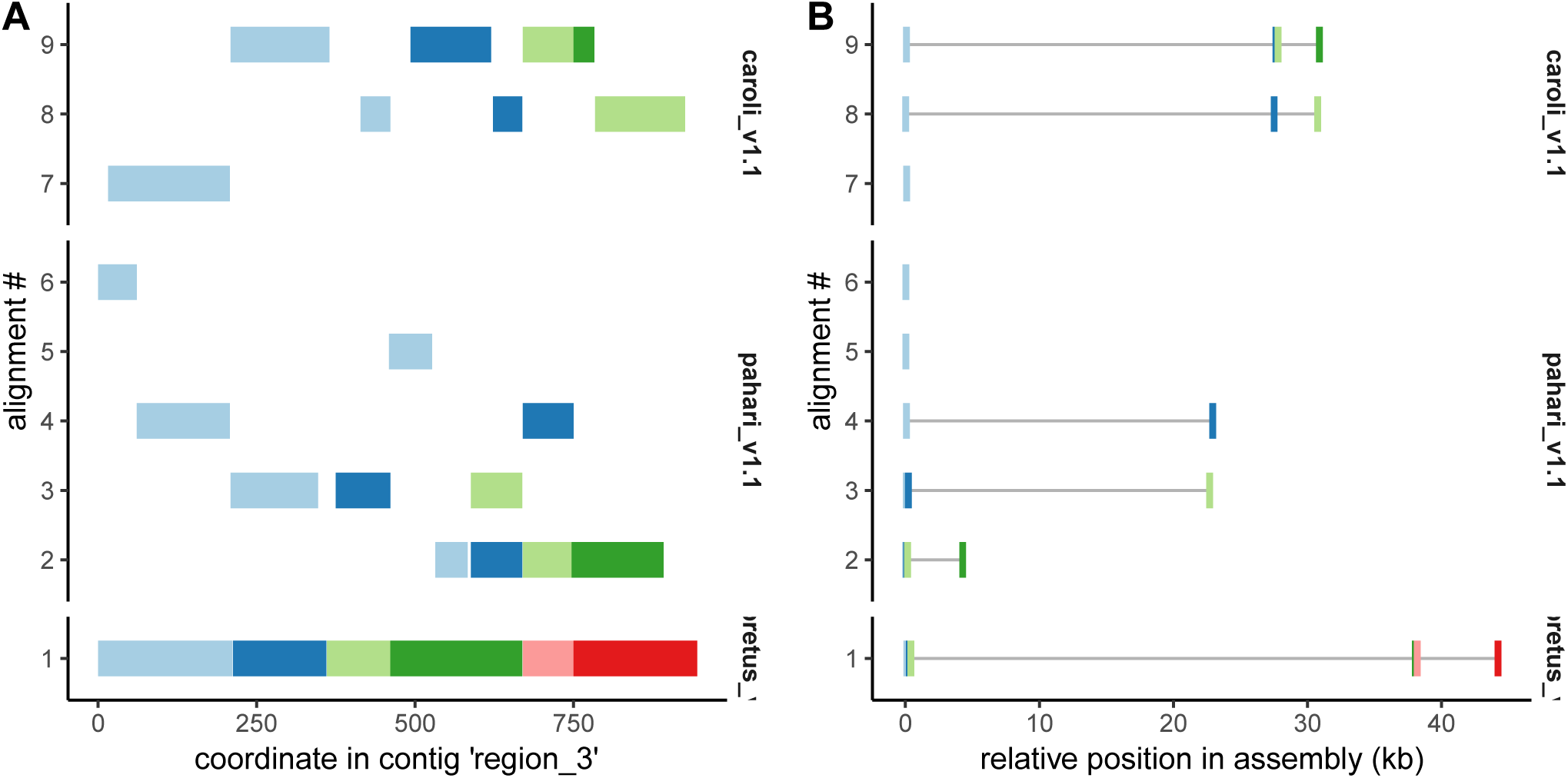
Alignment of breakpoint contig containing novel sequence (“region_3”) to *de novo* assemblies of out-group genomes. (**A**) Aligned segments relative to region_3. Each gapped alignment appears on its own line; segments within each alignment are color-coded. (**B**) Aligned segments relative to outgroup genomes. Coordinates are specified relative to an arbitrary offset to facilitate visual comparison. Segments are colored as in panel A.

**File S1.** Genetic map in the extended pseudoautosomal region.

**File S2.** Informative markers near the pseudoautosomal boundary on the 77K and 11K platforms.

**File S3.** Sequences of *de novo* assembled contigs spanning breakpoints of the PWK/PhJ Y-linked duplication (FASTA format).

**File S4.** List of N2 progeny used in this study.

**File S5.** Array genotypes and hybridization intensities from 11K and 77K platforms for N2 progeny and control inbred strains (ZIP archive).

**File S6.** List of samples analyzed by whole-genome sequencing (including previously published) in this study.

**File S7.** Alignments of a breakpoint-spanning contig (“region_3”) to *M. spretus, M. caroli* and *M. pahari* assemblies (UCSC PSL format).

**File S8.** Genotype calls from whole-genome sequencing in the region of the distal X chromosome corresponding to the PWK/PhJ Y-linked duplication (VCF format).

**File S9.** Breakpoints of copy-number alleles near the PAR boundary in wild mice.

