## Supplementary material for "Instability of the pseudoautosomal boundary in house mice": Figure S5

wild dom from DE
HG06 [XX]

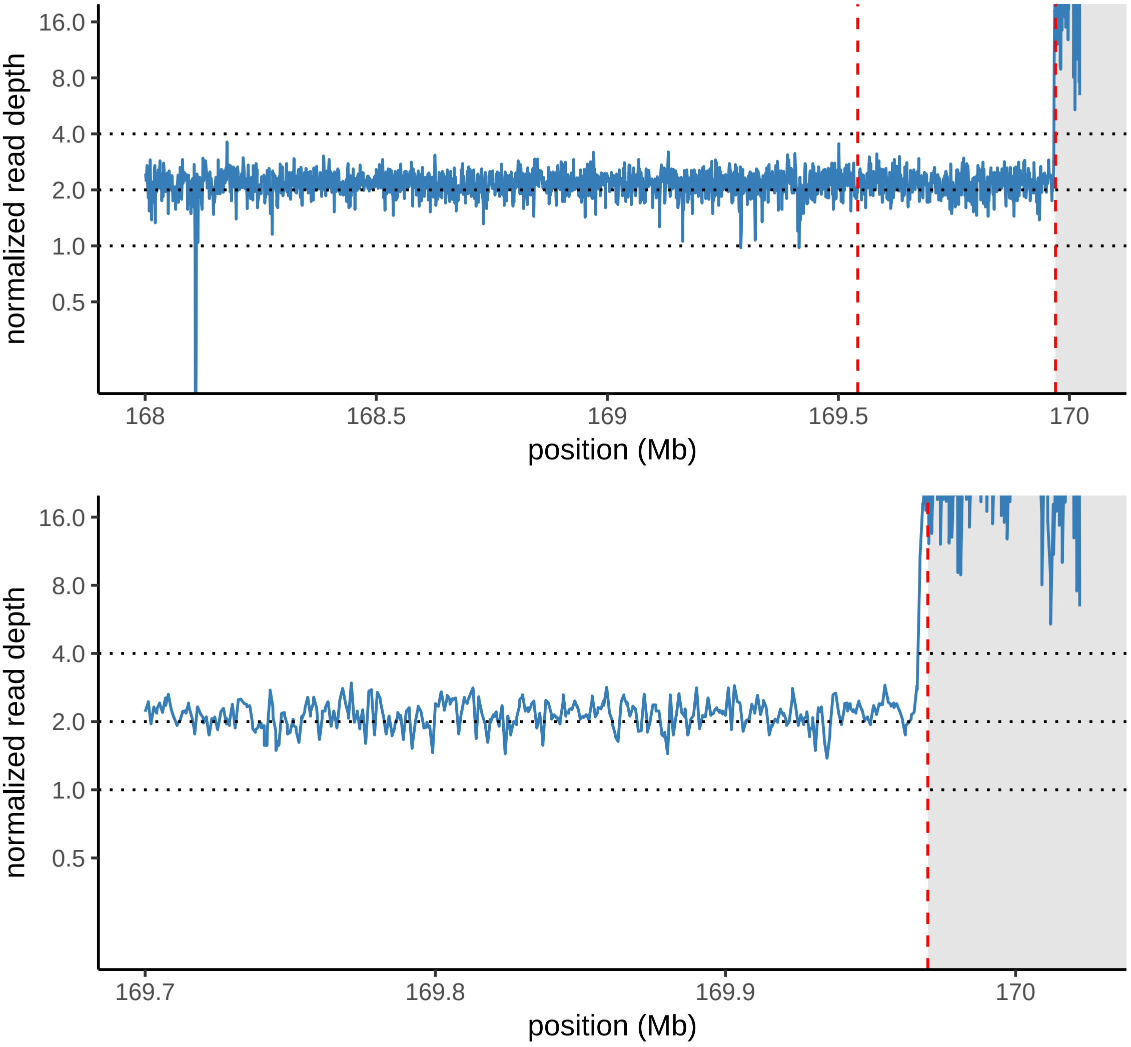

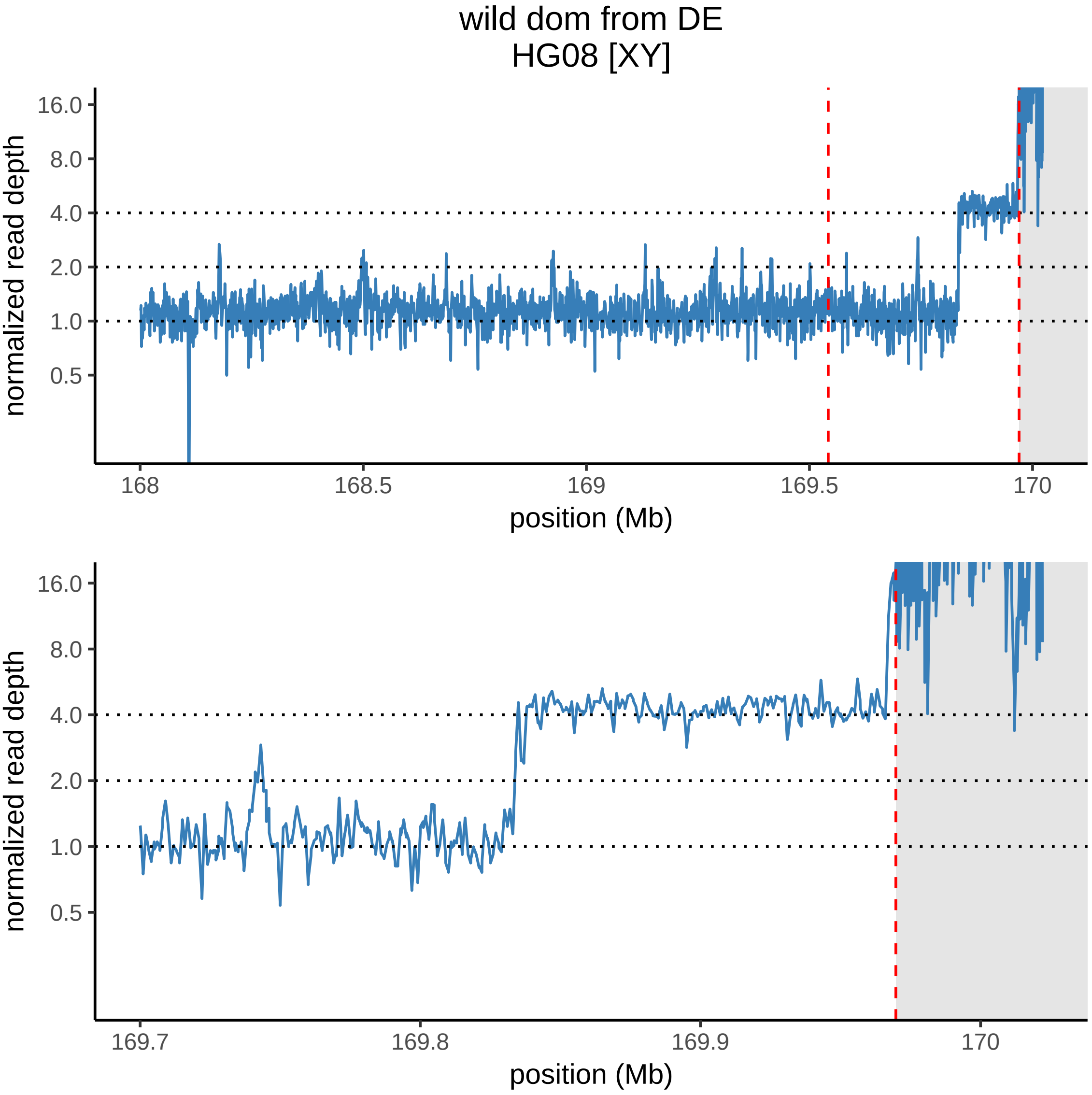

wild dom from DE
HG08 [XY]
normalized read depth
16.0
8.0
4.0
2.0
1.0
0.5
168
168.5
169
169.5
170
position (Mb)
normalized read depth
16.0
8.0
4.0
2.0
1.0
0.5
169.7
169.8
169.9
170
position (Mb)

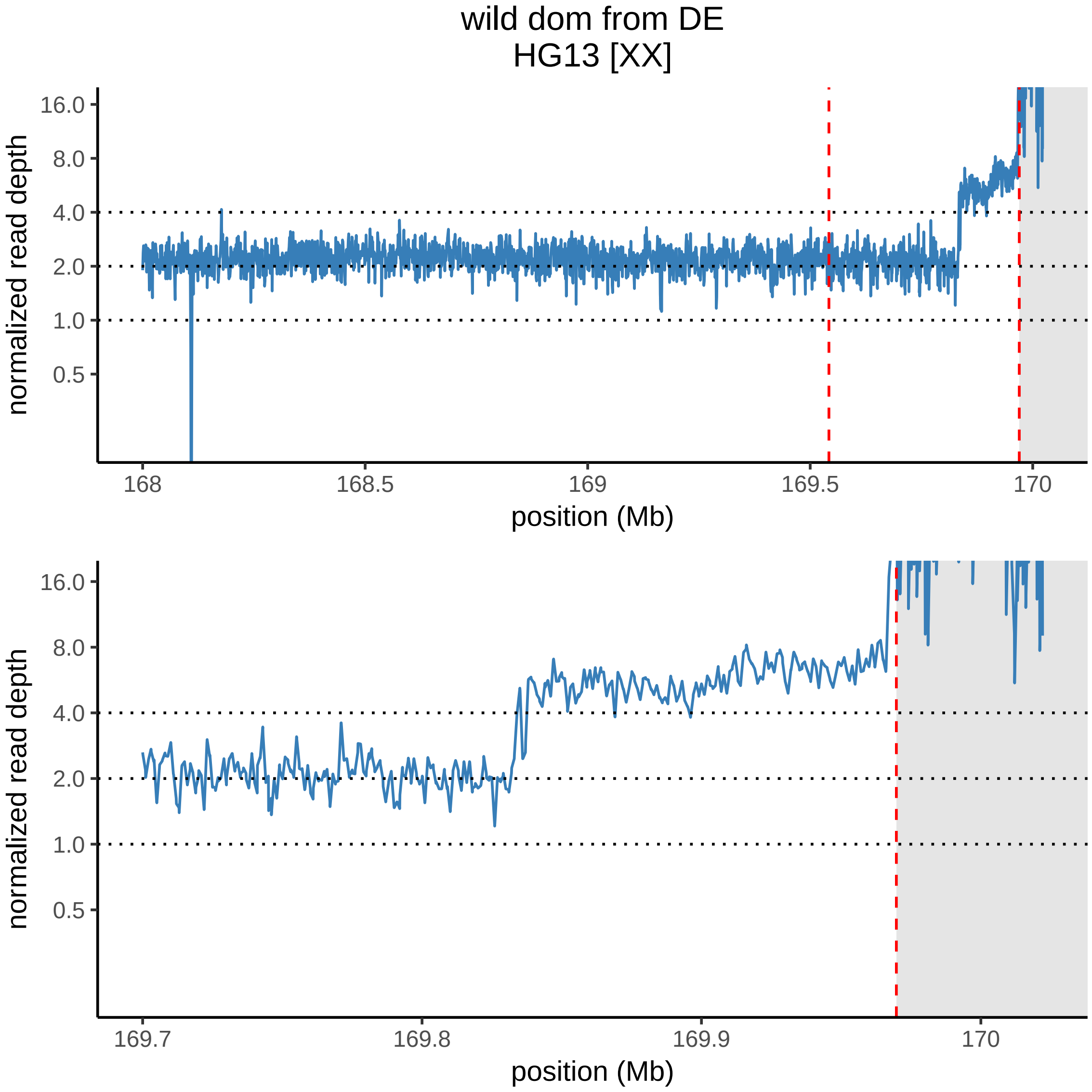

wild dom from DE
HG13 [XX]
normalized read depth
16.0
8.0
4.0
2.0
1.0
0.5
168
168.5
169
169.5
170
position (Mb)
normalized read depth
16.0
8.0
4.0
2.0
1.0
0.5
169.7
169.8
169.9
170
position (Mb)

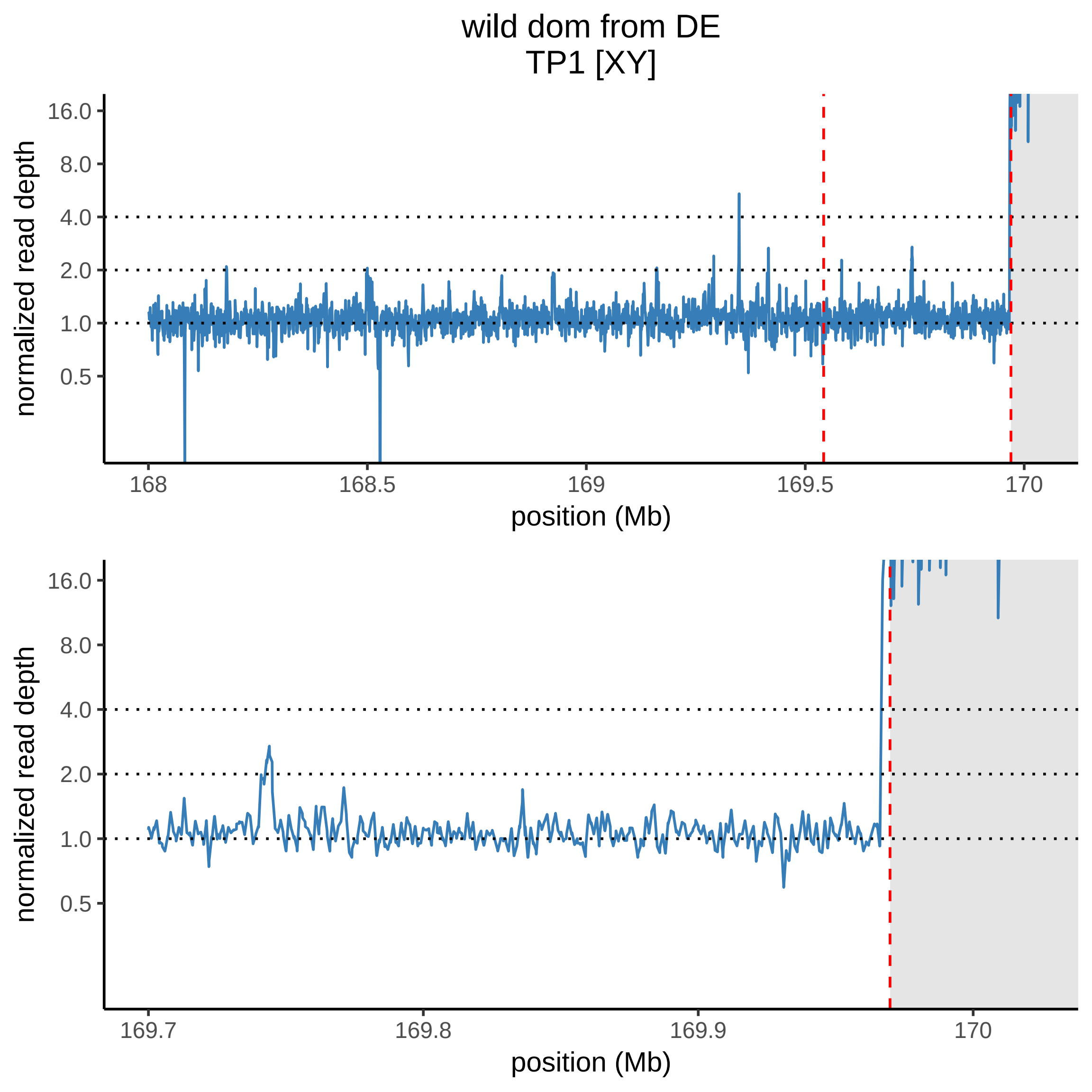

wild dom from DE
TP1 [XY]
normalized read depth
16.0
8.0
4.0
2.0
1.0
0.5
168
168.5
169
169.5
170
position (Mb)
normalized read depth
16.0
8.0
4.0
2.0
1.0
0.5
169.7
169.8
169.9
170
position (Mb)

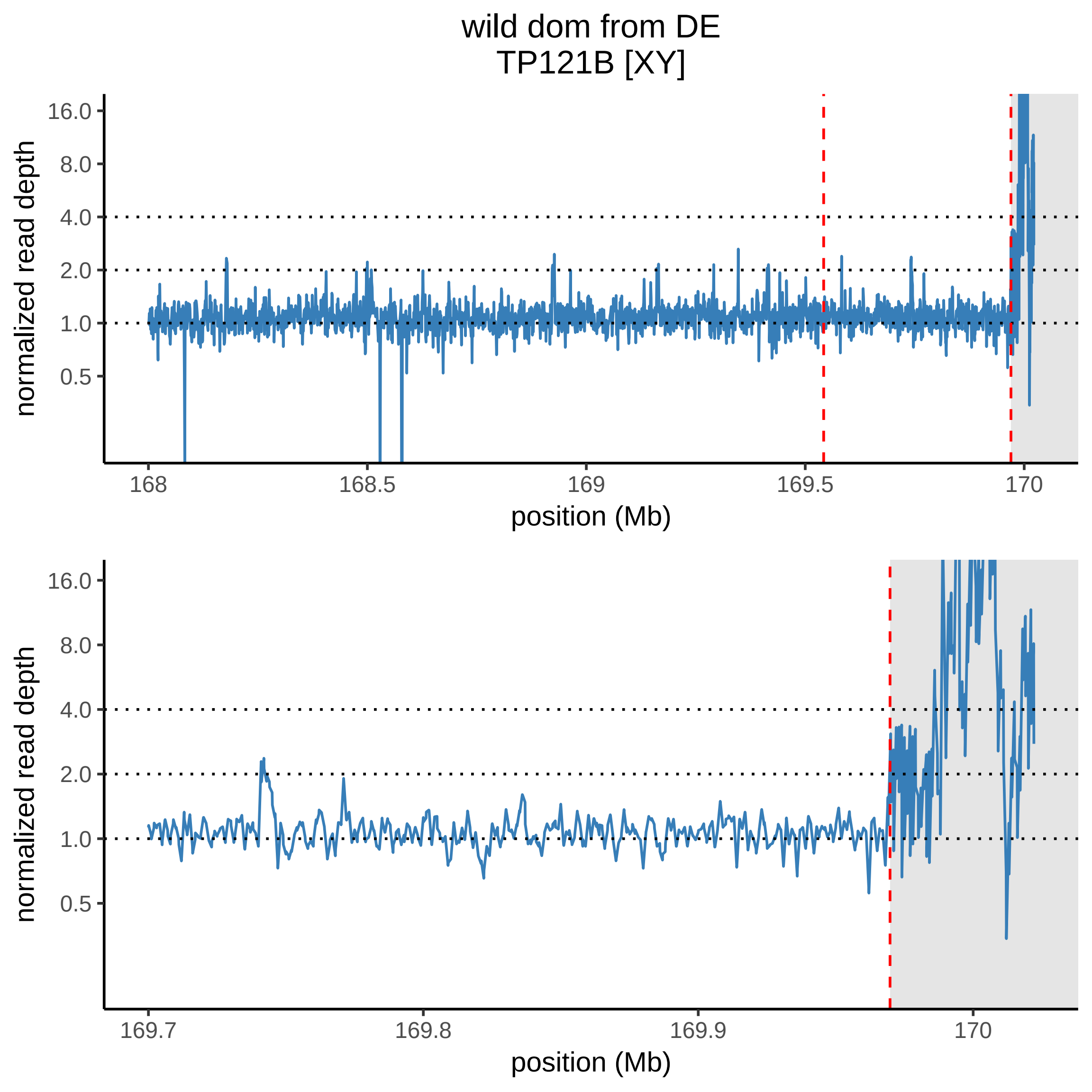

wild dom from DE
TP121B [XY]
normalized read depth
16.0
8.0
4.0
2.0
1.0
0.5
168
168.5
169
169.5
170
position (Mb)
normalized read depth
16.0
8.0
4.0
2.0
1.0
0.5
169.7
169.8
169.9
170
position (Mb)

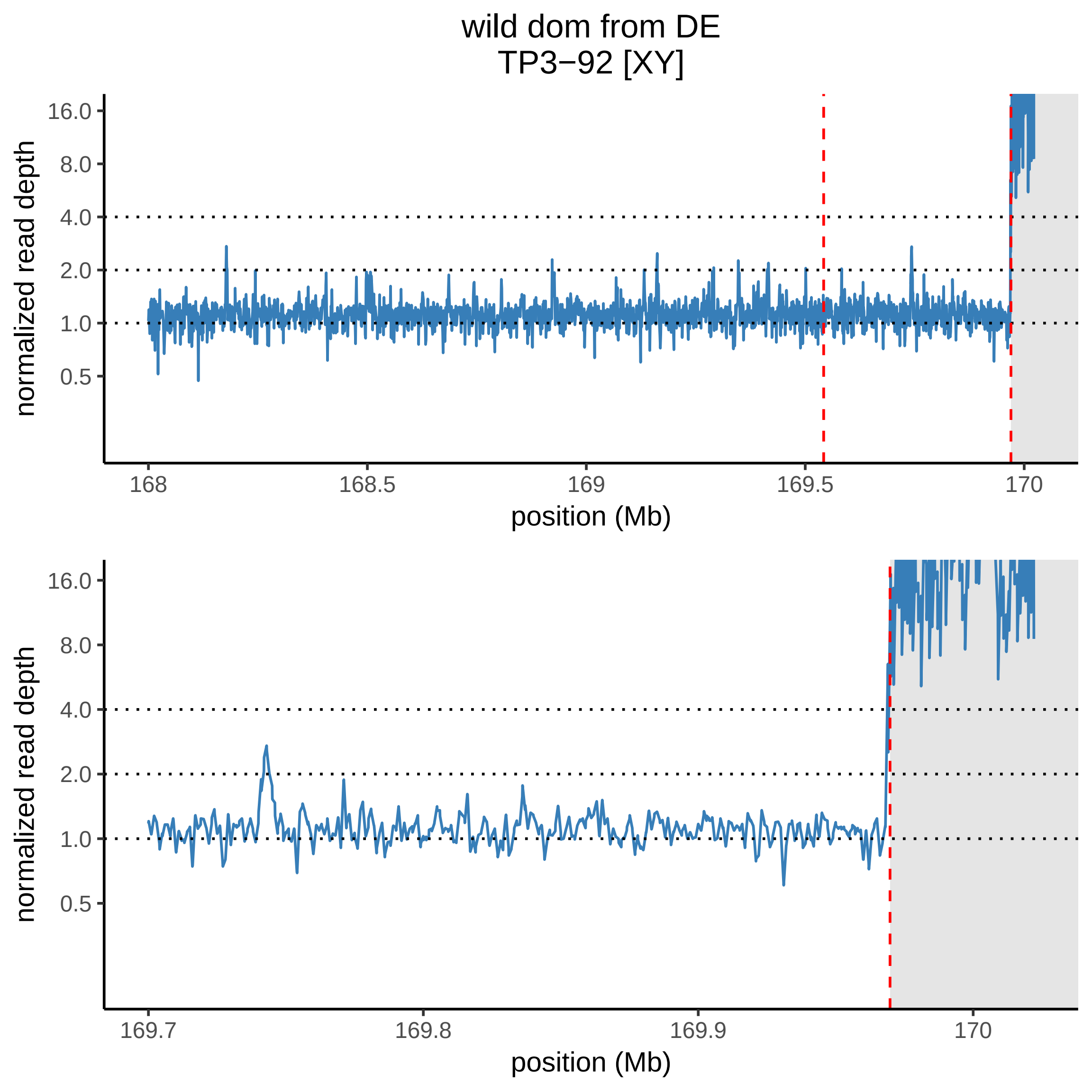

wild dom from DE
TP3−92 [XY]
16.0
8.0
4.0
2.0
1.0
0.5
normalized read depth
168
168.5
169
169.5
170
position (Mb)
16.0
8.0
4.0
2.0
1.0
0.5
normalized read depth
169.7
169.8
169.9
170
position (Mb)

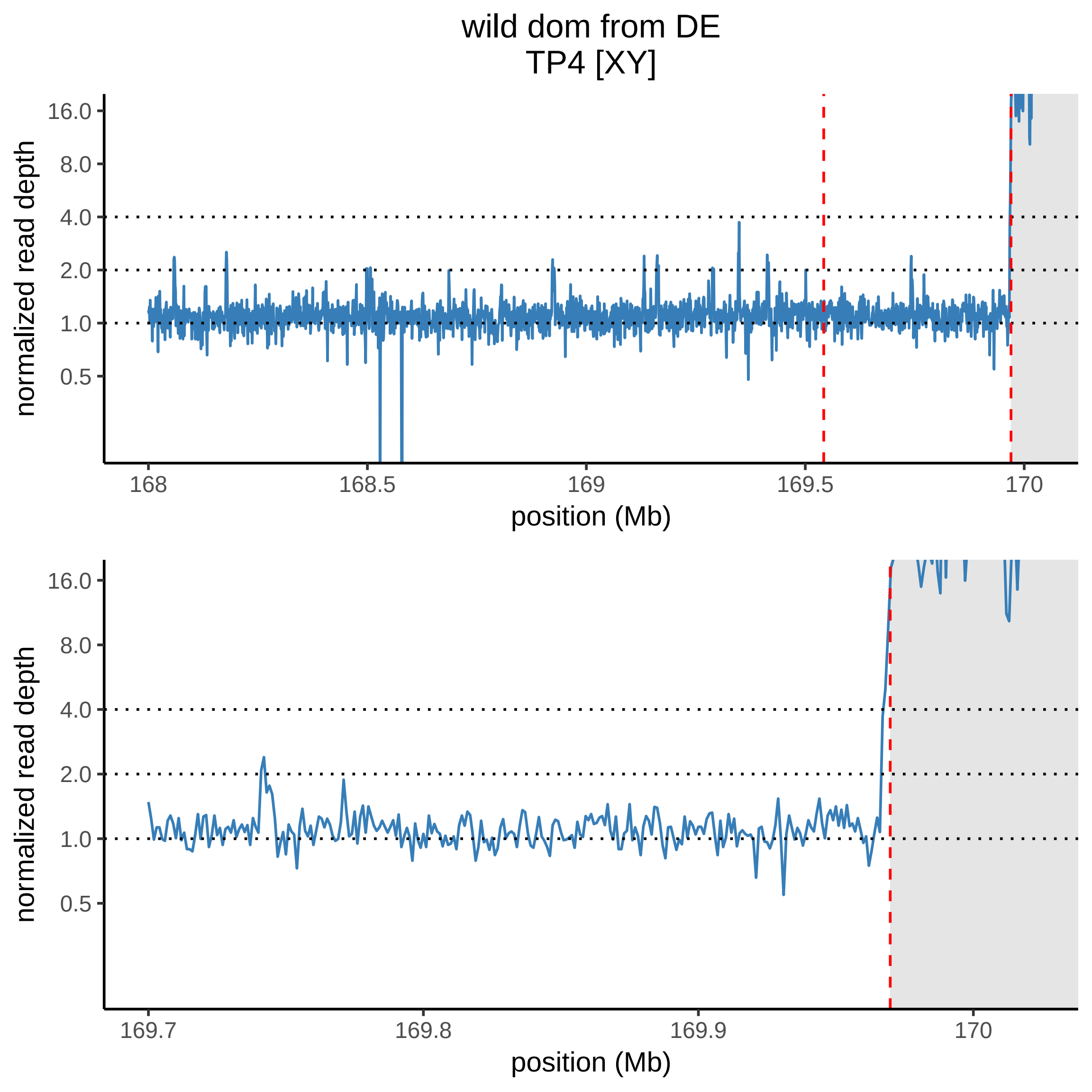

wild dom from DE
TP4 [XY]
normalized read depth
16.0
8.0
4.0
2.0
1.0
0.5
168
168.5
169
169.5
170
position (Mb)
normalized read depth
16.0
8.0
4.0
2.0
1.0
0.5
169.7
169.8
169.9
170
position (Mb)

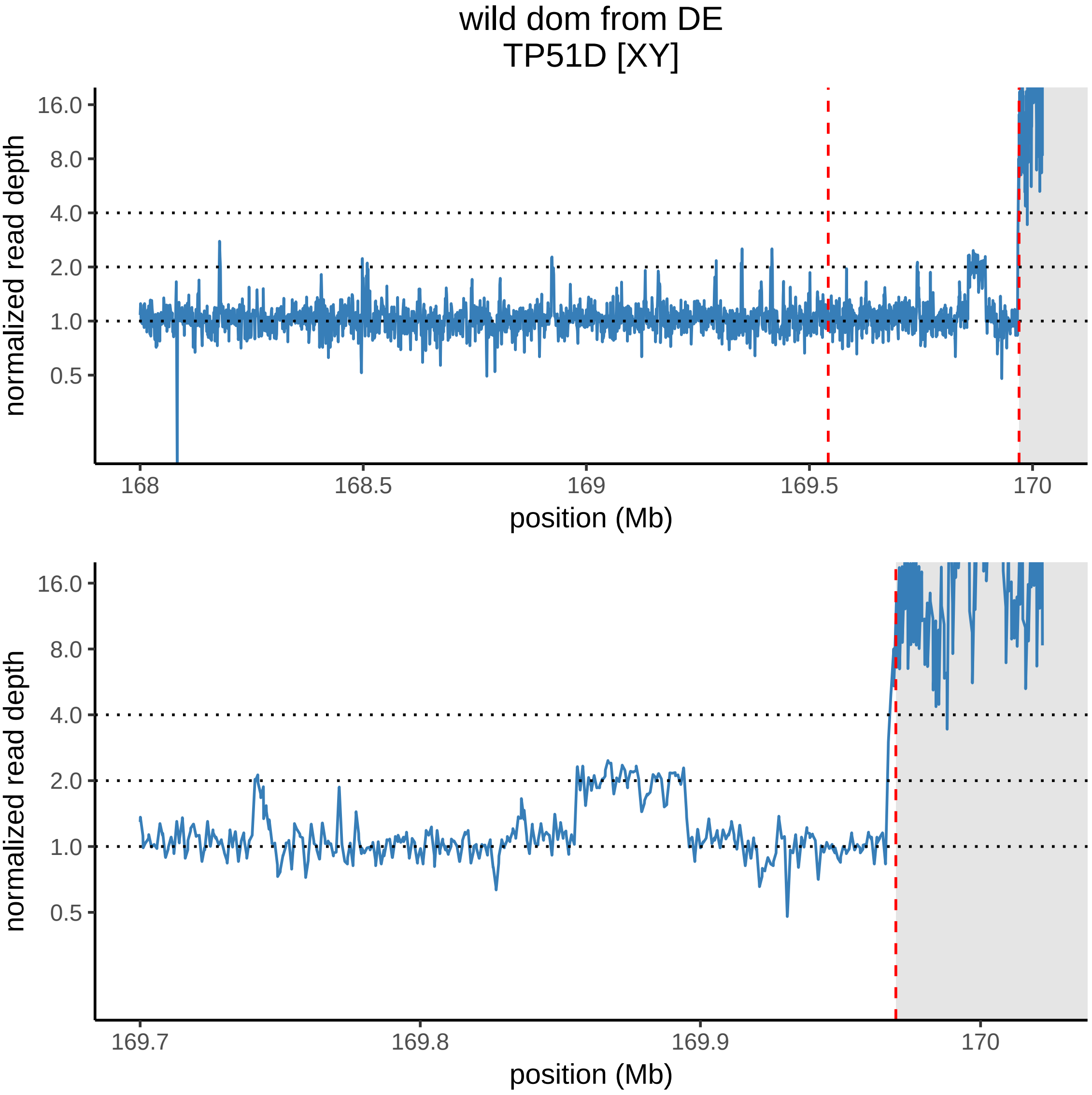

wild dom from DE
TP51D [XY]
normalized read depth
16.0
8.0
4.0
2.0
1.0
0.5
168
168.5
169
169.5
170
position (Mb)
normalized read depth
16.0
8.0
4.0
2.0
1.0
0.5
169.7
169.8
169.9
170
position (Mb)

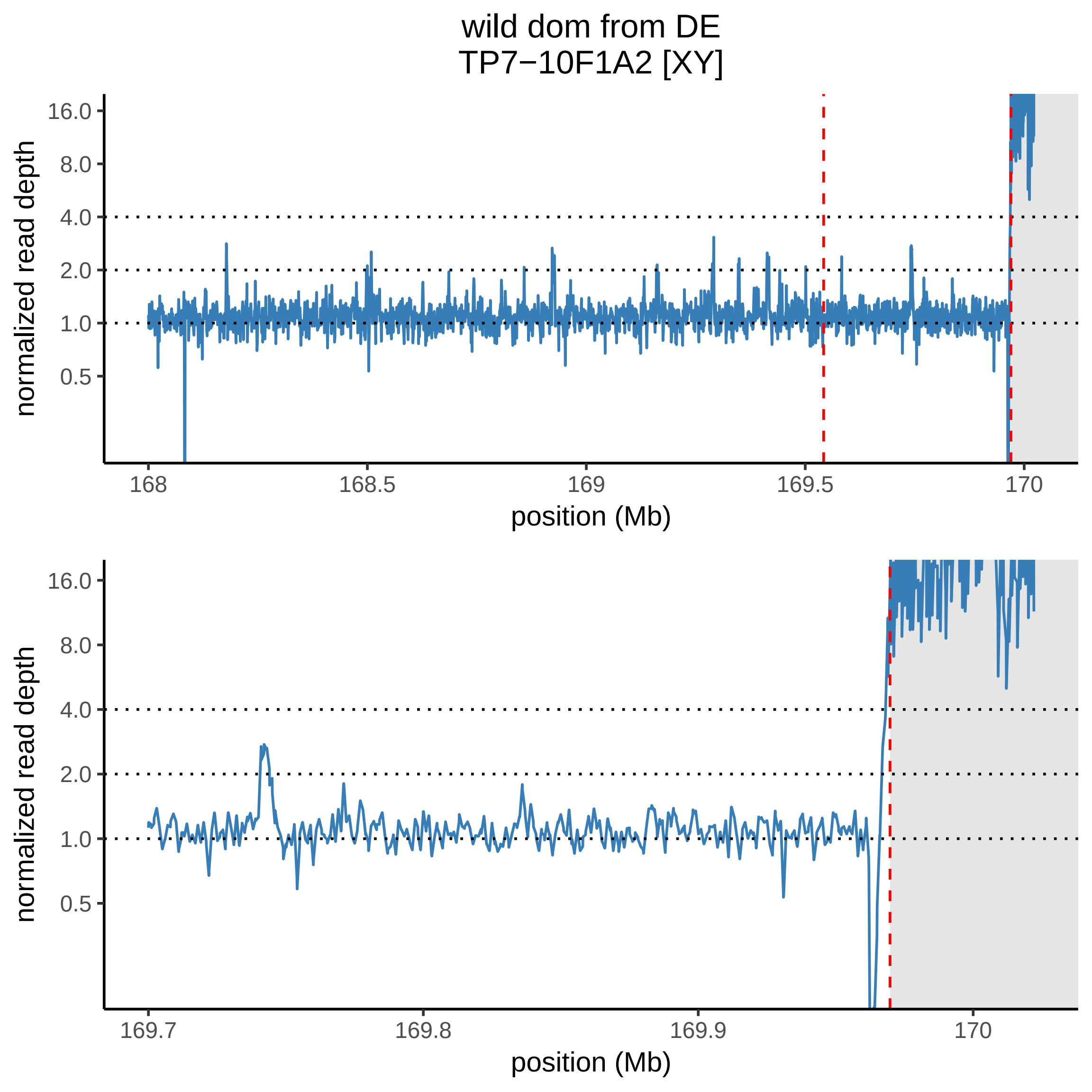
wild dom from DE
TP7−10F1A2 [XY]
normalized read depth
16.0
8.0
4.0
2.0
1.0
0.5
168
168.5
169
169.5
170
position (Mb)
normalized read depth
16.0
8.0
4.0
2.0
1.0
0.5
169.7
169.8
169.9
170
position (Mb)

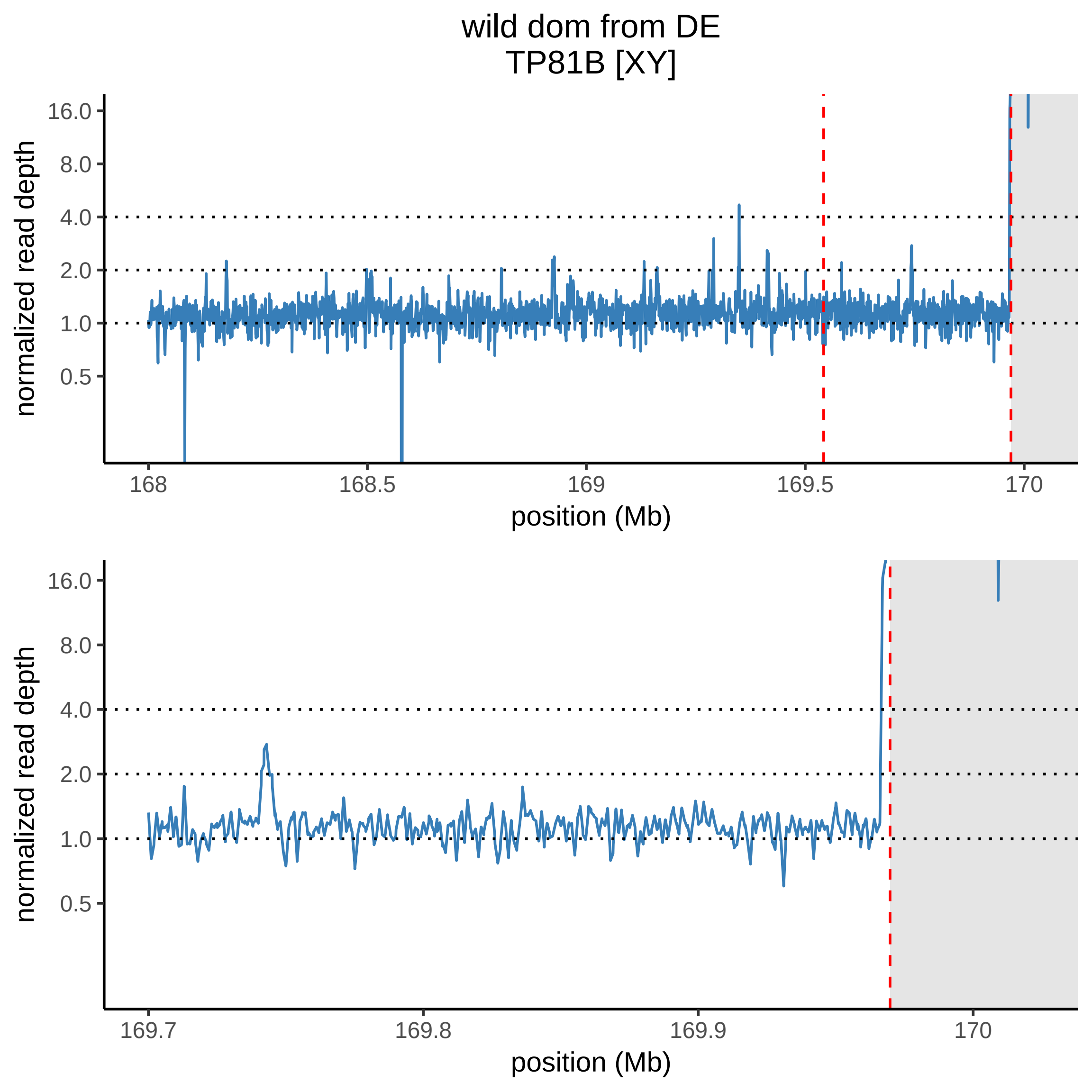
wild dom from DE
TP81B [XY]
normalized read depth
16.0
8.0
4.0
2.0
1.0
0.5
168
168.5
169
169.5
170
position (Mb)
normalized read depth
16.0
8.0
4.0
2.0
1.0
0.5
169.7
169.8
169.9
170
position (Mb)

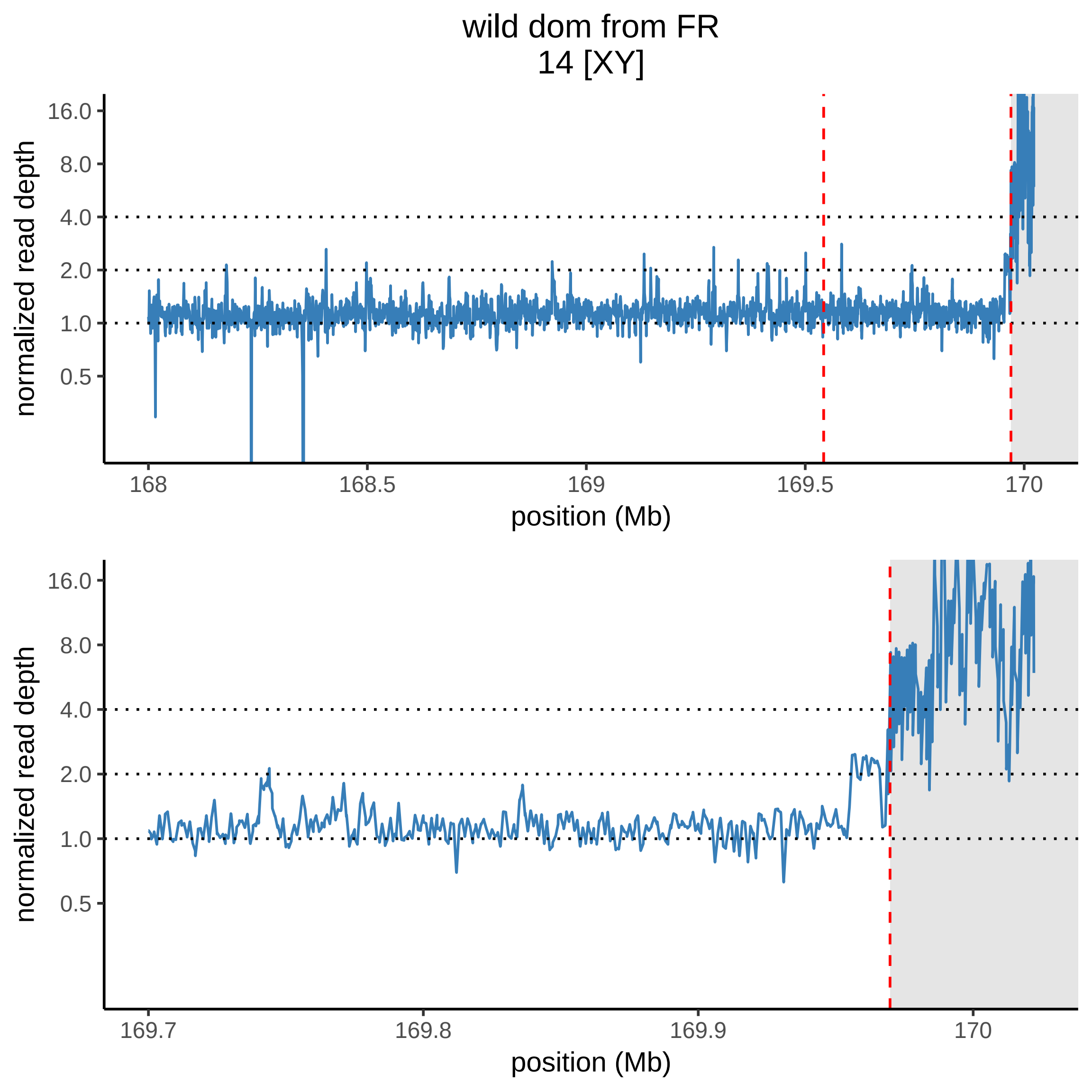

wild dom from FR
14 [XY]
normalized read depth
16.0
8.0
4.0
2.0
1.0
0.5
168
168.5
169
169.5
170
position (Mb)
normalized read depth
16.0
8.0
4.0
2.0
1.0
0.5
169.7
169.8
169.9
170
position (Mb)

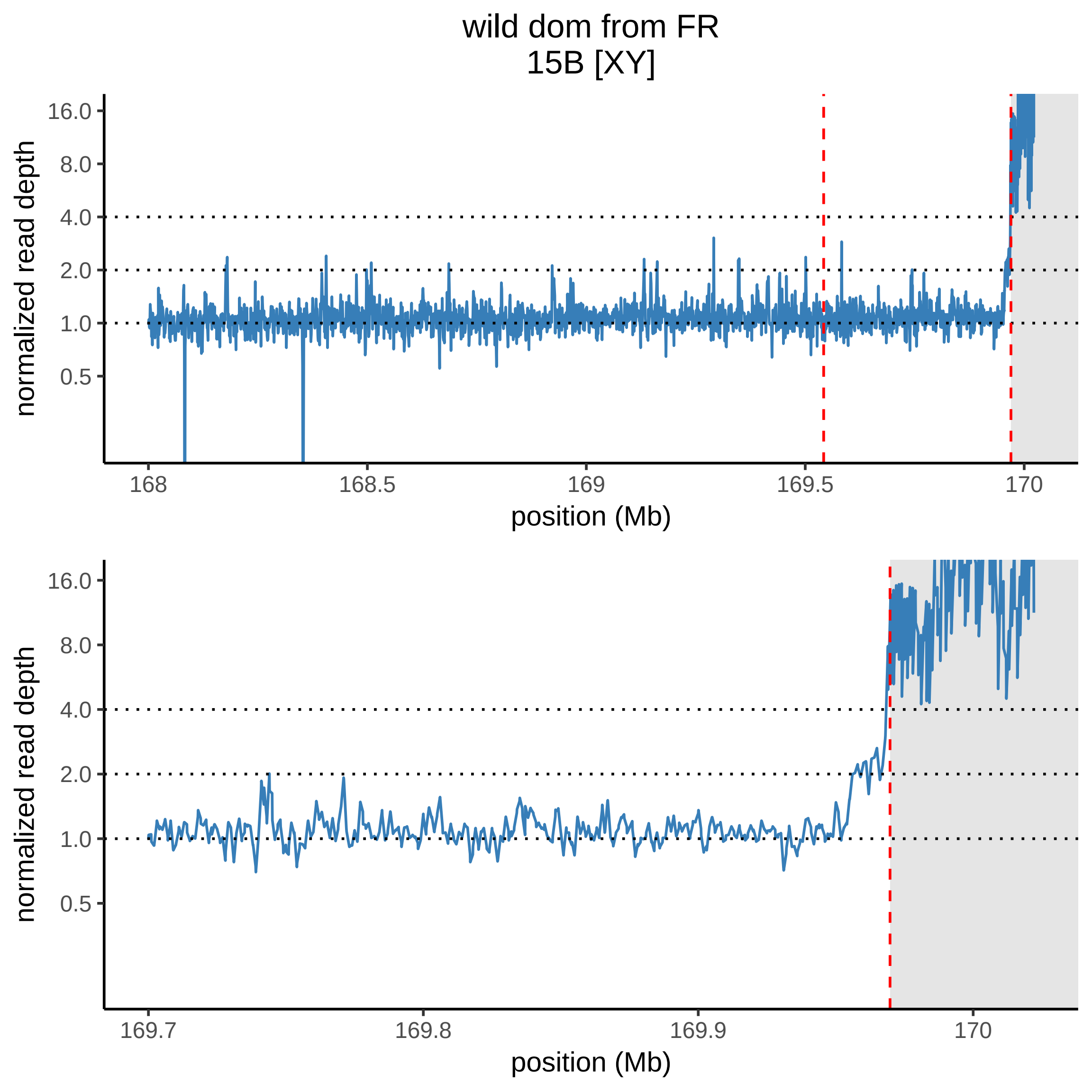
wild dom from FR
15B [XY]
normalized read depth
16.0
8.0
4.0
2.0
1.0
0.5
168
168.5
169
169.5
170
position (Mb)
normalized read depth
16.0
8.0
4.0
2.0
1.0
0.5
169.7
169.8
169.9
170
position (Mb)

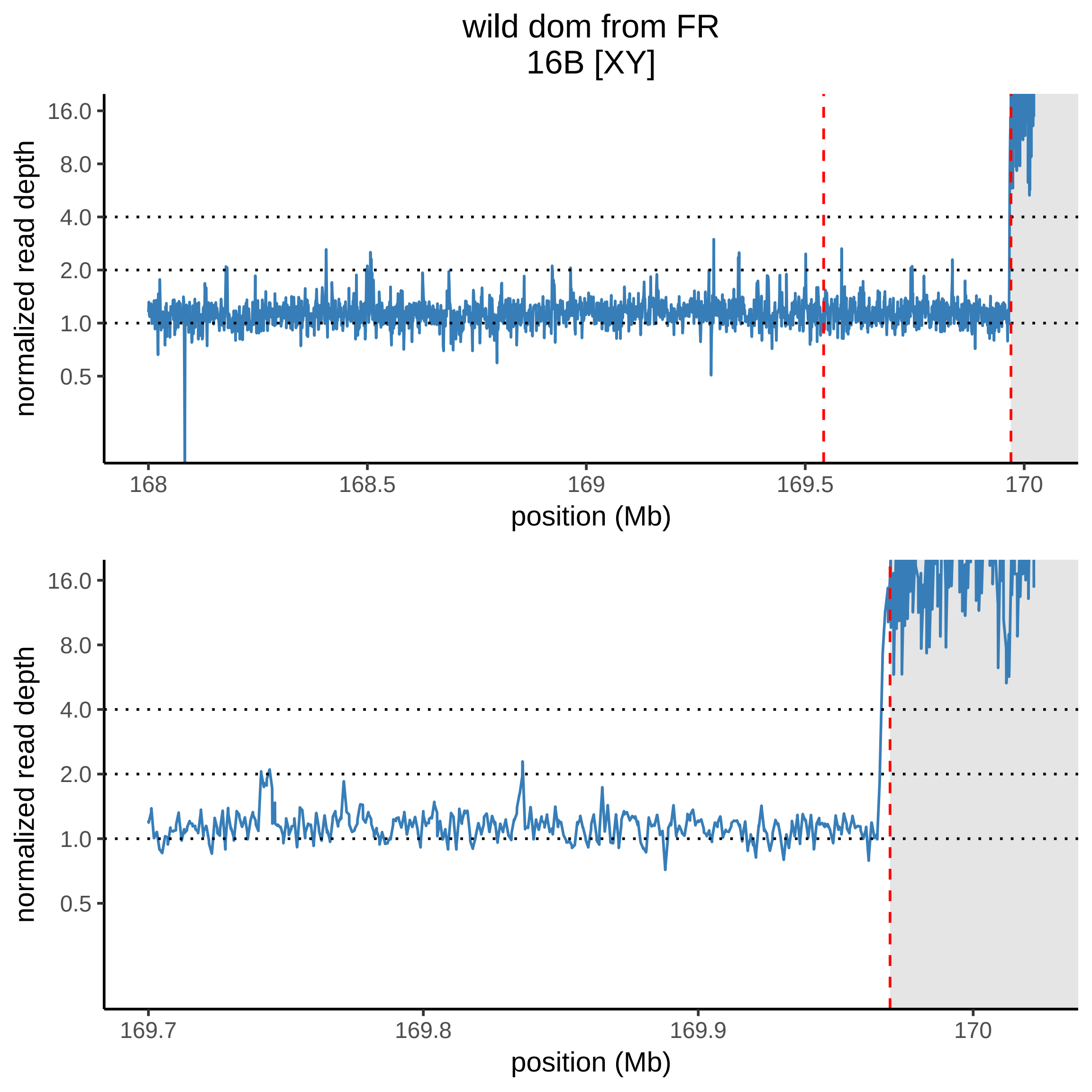

wild dom from FR
16B [XY]
normalized read depth
16.0
8.0
4.0
2.0
1.0
0.5
168
168.5
169
169.5
170
position (Mb)
normalized read depth
16.0
8.0
4.0
2.0
1.0
0.5
169.7
169.8
169.9
170
position (Mb)

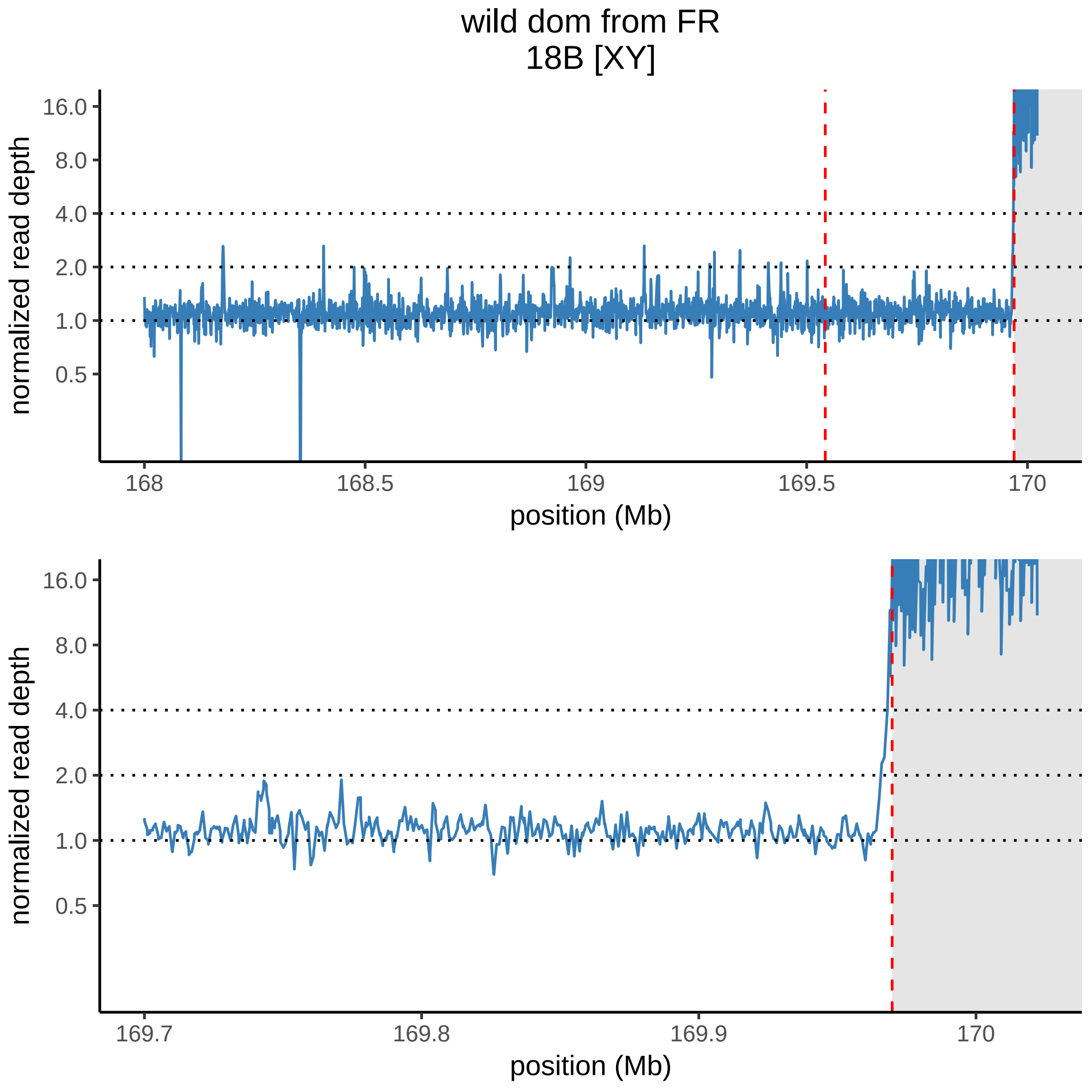
wild dom from FR
18B [XY]
normalized read depth
16.0
8.0
4.0
2.0
1.0
0.5
168
168.5
169
169.5
170
position (Mb)
normalized read depth
16.0
8.0
4.0
2.0
1.0
0.5
169.7
169.8
169.9
170
position (Mb)

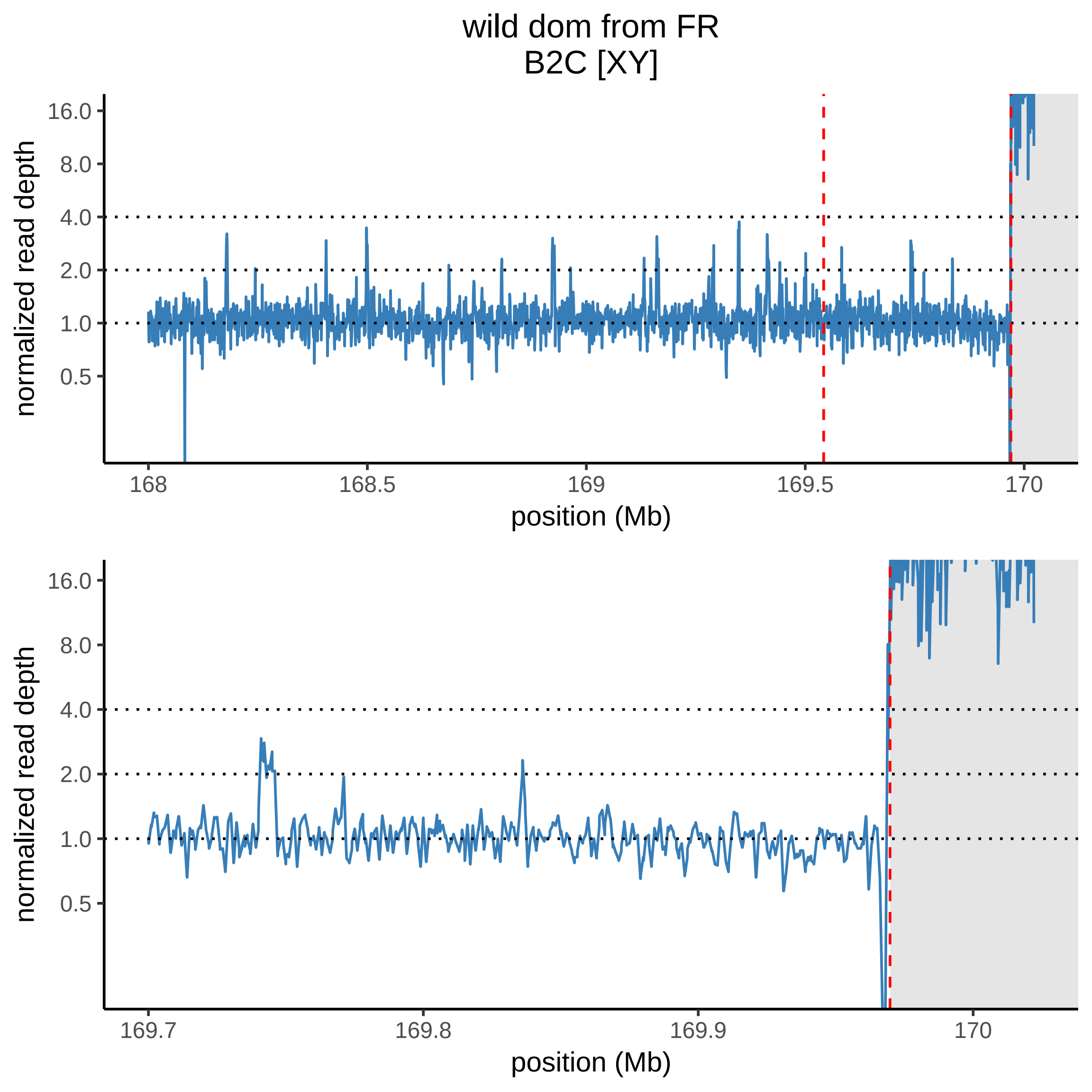

wild dom from FR
B2C [XY]
normalized read depth
16.0
8.0
4.0
2.0
1.0
0.5
168
168.5
169
169.5
170
position (Mb)
normalized read depth
16.0
8.0
4.0
2.0
1.0
0.5
169.7
169.8
169.9
170
position (Mb)

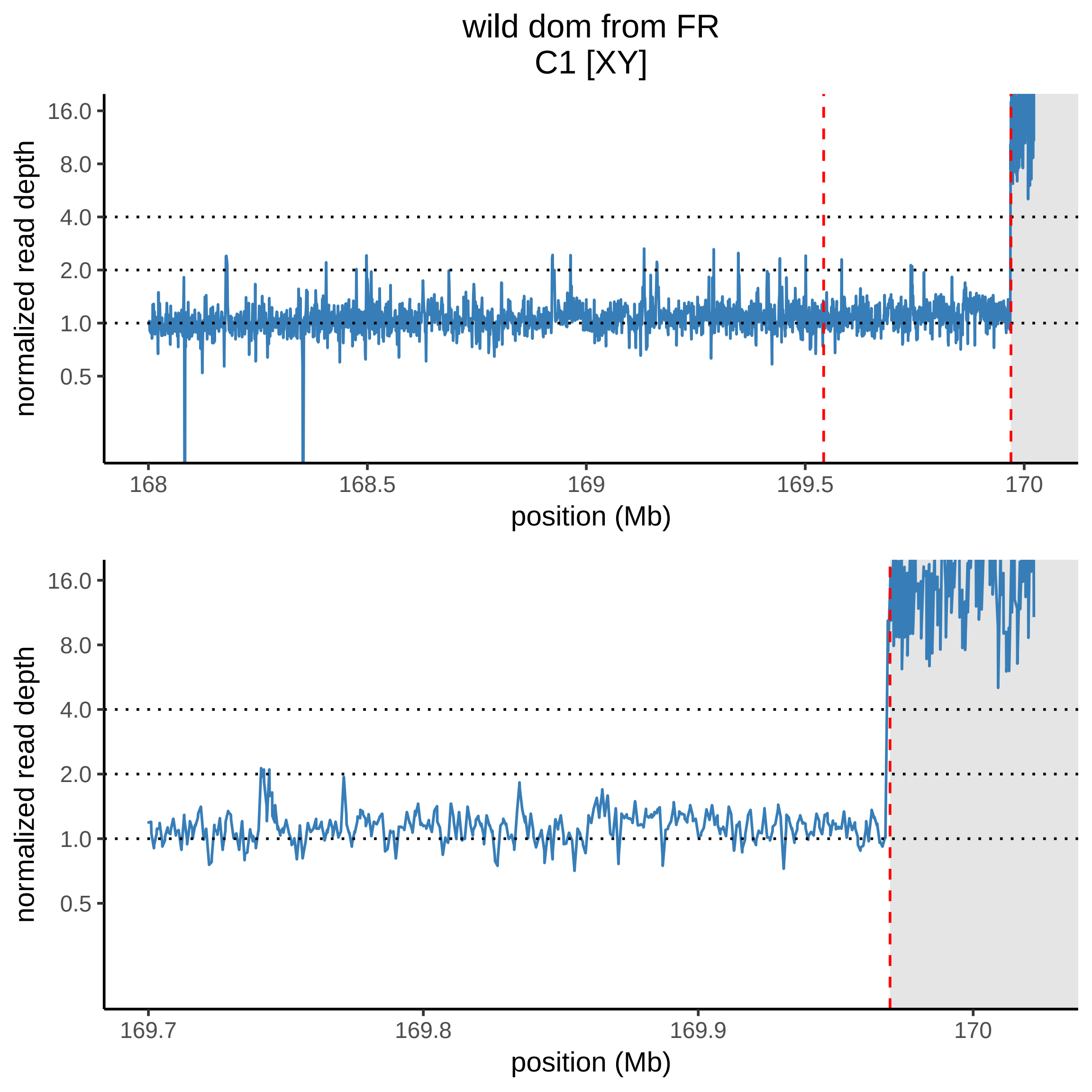
wild dom from FR
C1 [XY]
normalized read depth
16.0
8.0
4.0
2.0
1.0
0.5
168
168.5
169
169.5
170
position (Mb)
normalized read depth
16.0
8.0
4.0
2.0
1.0
0.5
169.7
169.8
169.9
170
position (Mb)

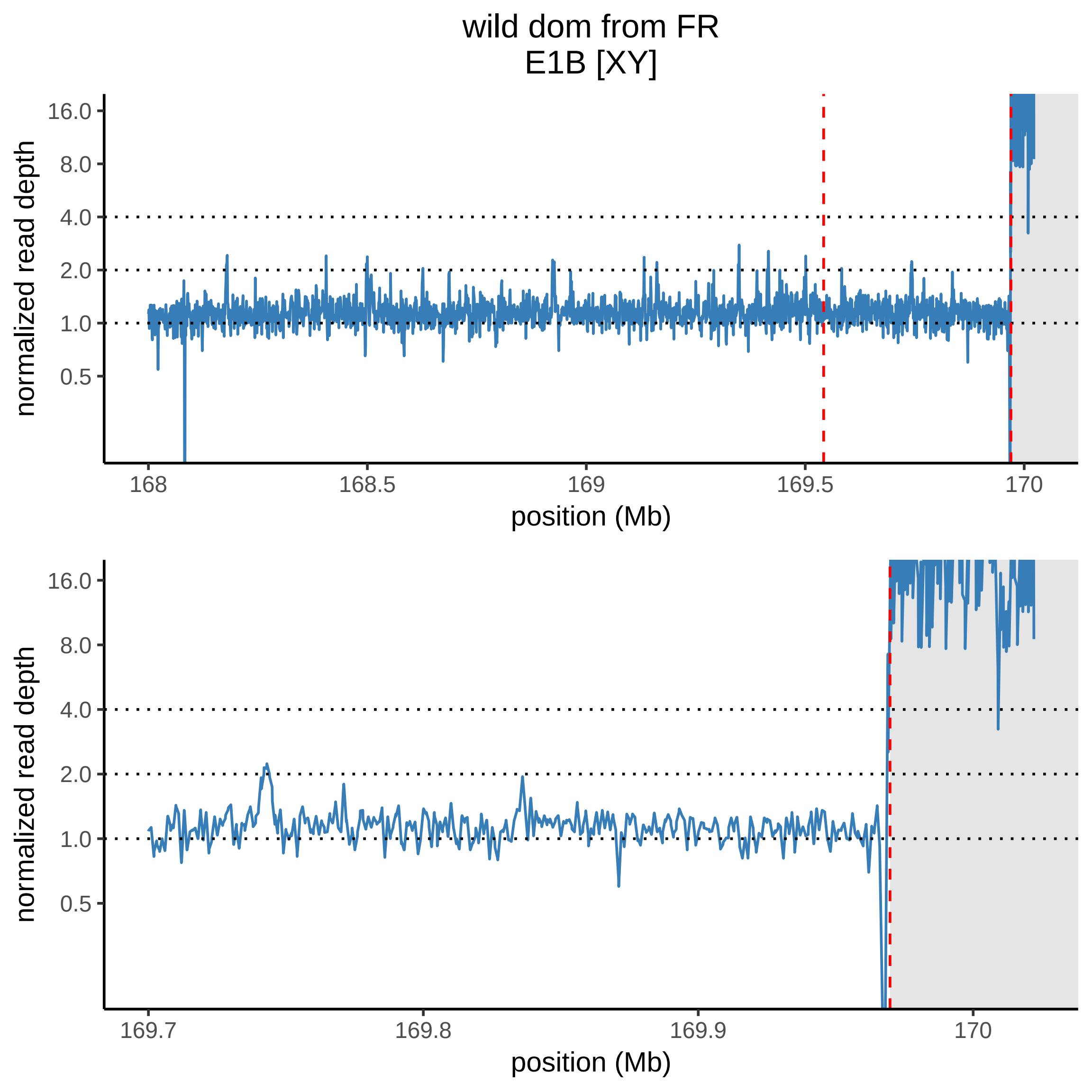

wild dom from FR
E1B [XY]
normalized read depth
16.0
8.0
4.0
2.0
1.0
0.5
168
168.5
169
169.5
170
position (Mb)
normalized read depth
16.0
8.0
4.0
2.0
1.0
0.5
169.7
169.8
169.9
170
position (Mb)

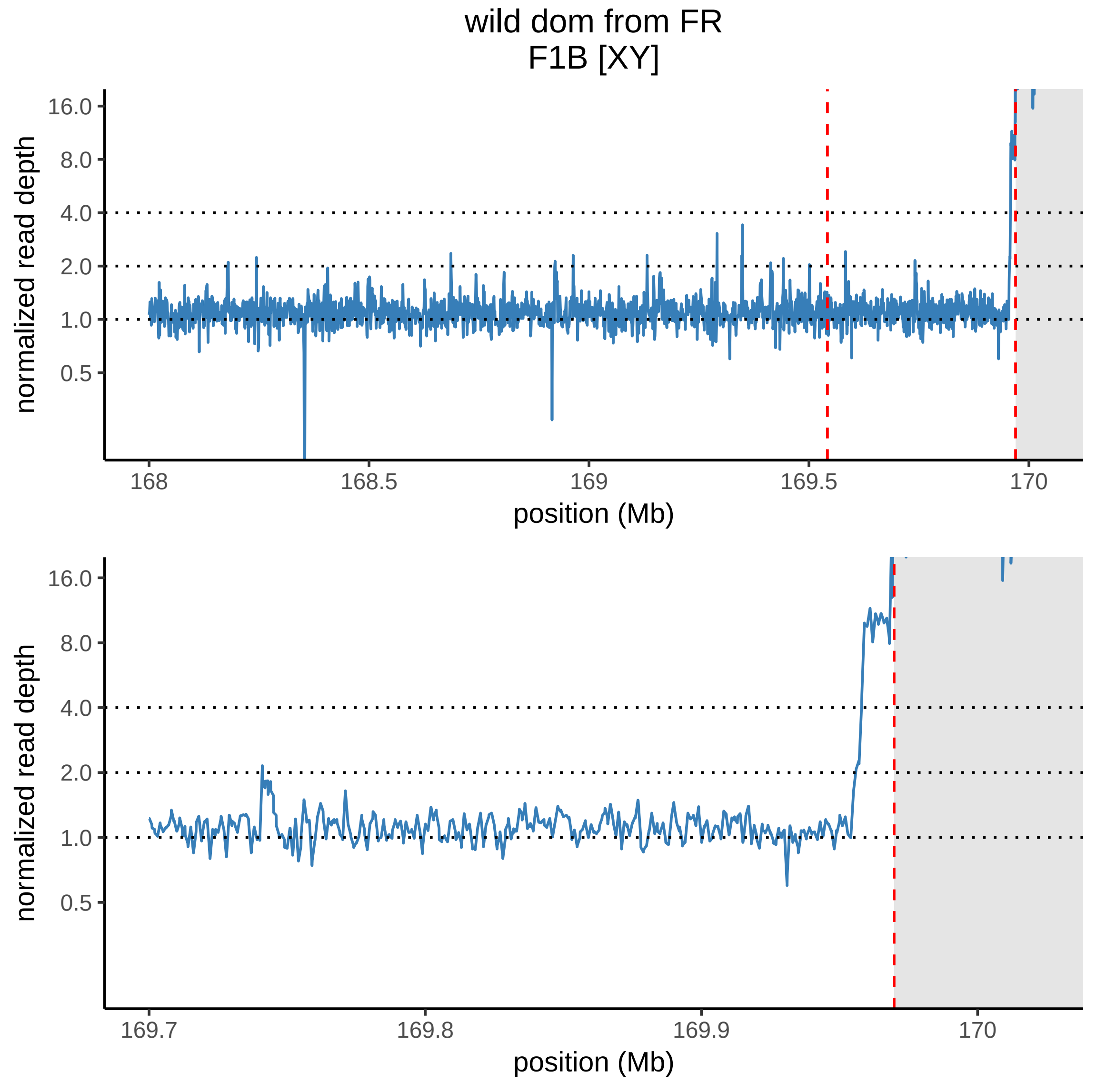
wild dom from FR
F1B [XY]
normalized read depth
16.0
8.0
4.0
2.0
1.0
0.5
168
168.5
169
169.5
170
position (Mb)
normalized read depth
16.0
8.0
4.0
2.0
1.0
0.5
169.7
169.8
169.9
170
position (Mb)

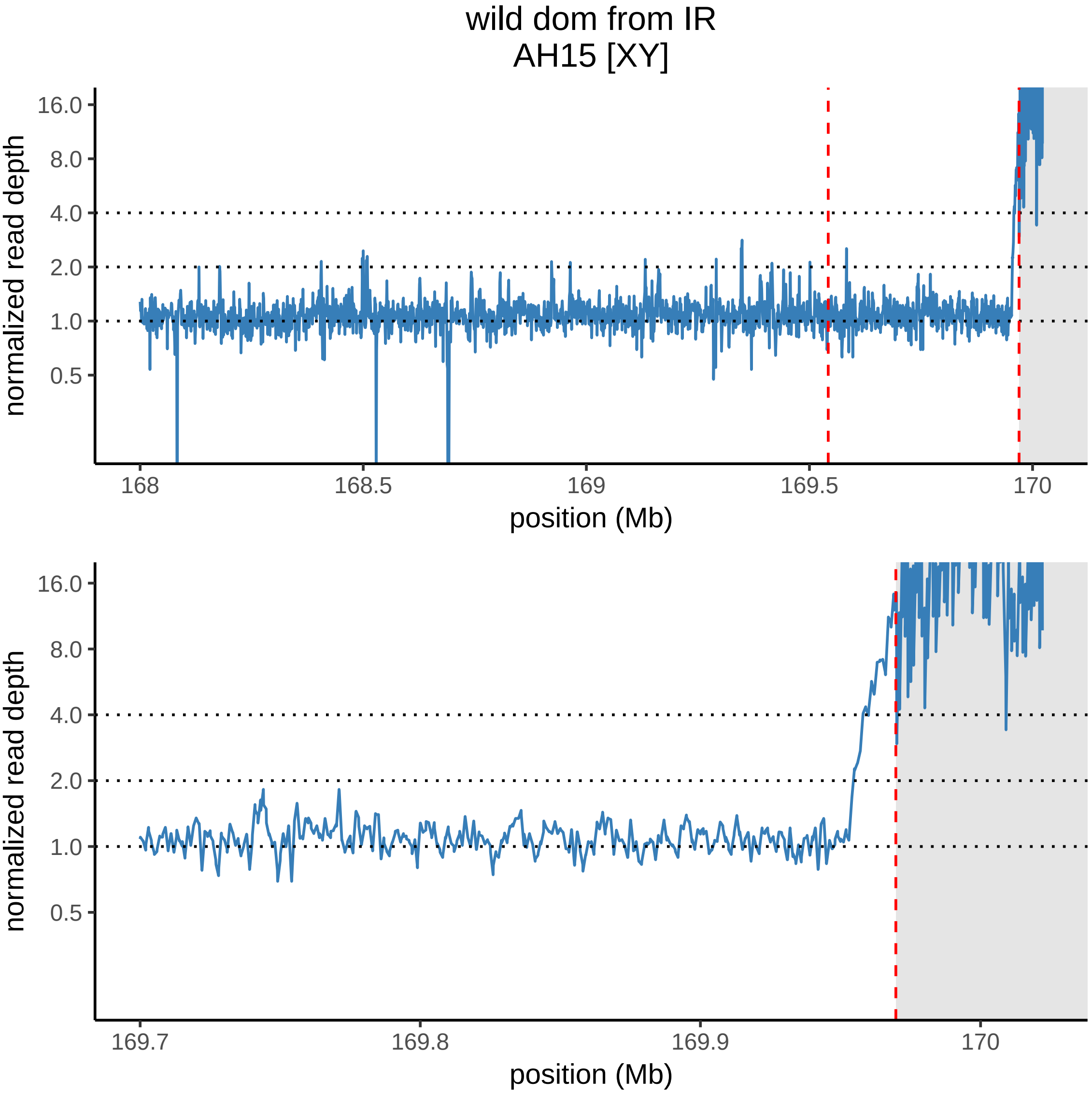
wild dom from IR
AH15 [XY]
normalized read depth
16.0
8.0
4.0
2.0
1.0
0.5
168
168.5
169
169.5
170
position (Mb)
normalized read depth
16.0
8.0
4.0
2.0
1.0
0.5
169.7
169.8
169.9
170
position (Mb)

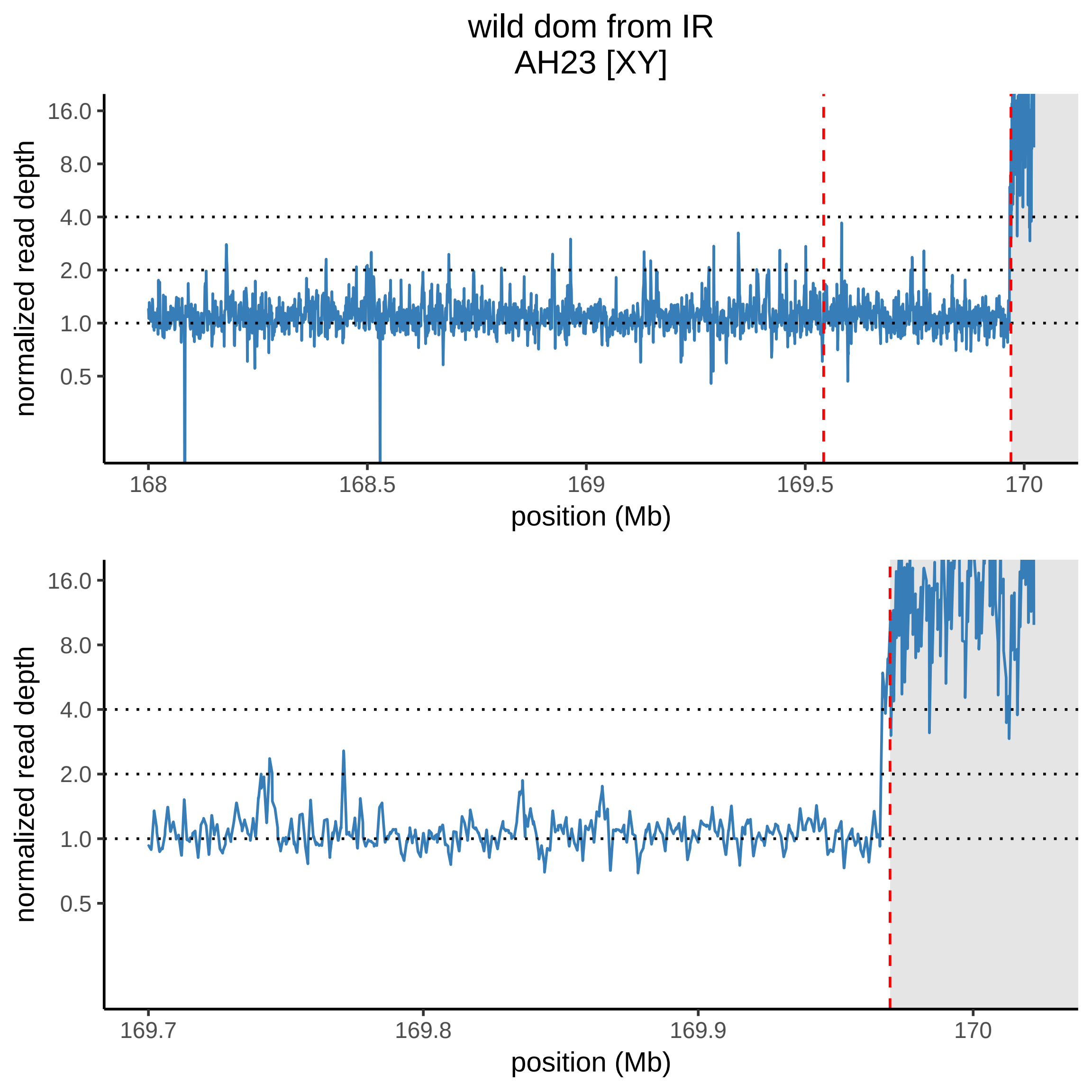

wild dom from IR
AH23 [XY]
normalized read depth
16.0
8.0
4.0
2.0
1.0
0.5
168
168.5
169
169.5
170
position (Mb)
normalized read depth
16.0
8.0
4.0
2.0
1.0
0.5
169.7
169.8
169.9
170
position (Mb)

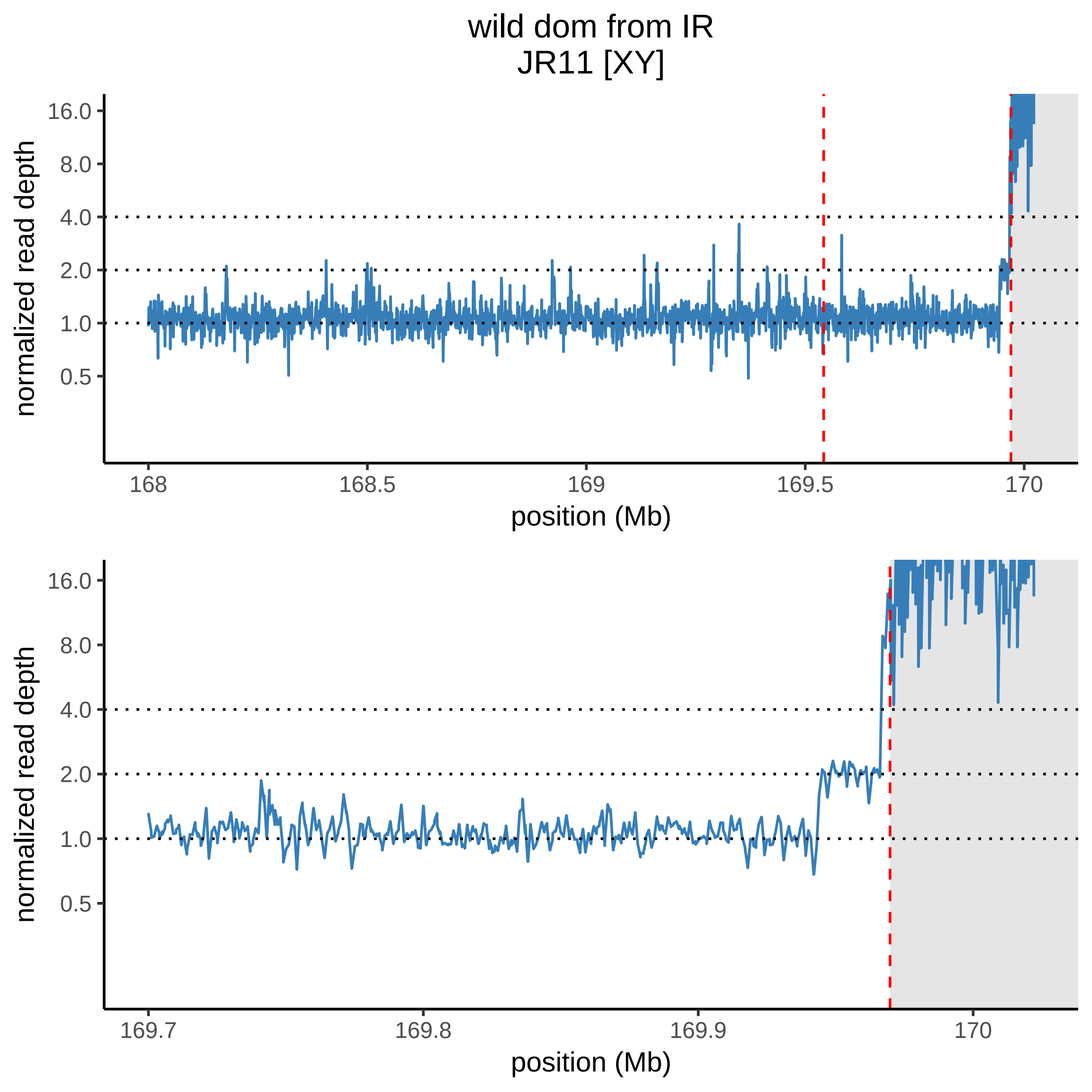

wild dom from IR
JR11 [XY]
normalized read depth
16.0
8.0
4.0
2.0
1.0
0.5
168
168.5
169
169.5
170
position (Mb)
normalized read depth
16.0
8.0
4.0
2.0
1.0
0.5
169.7
169.8
169.9
170
position (Mb)

### wild dom from IR
JR15 [XY]

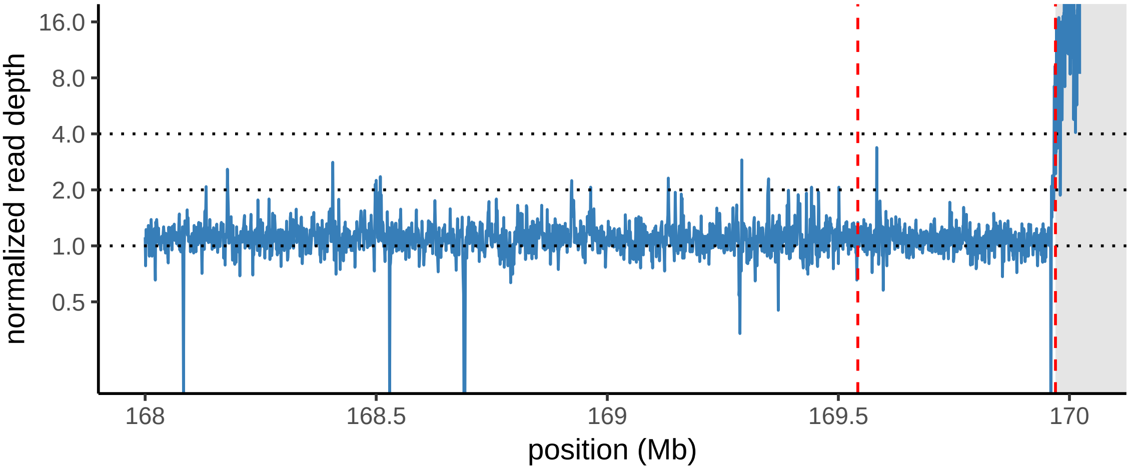


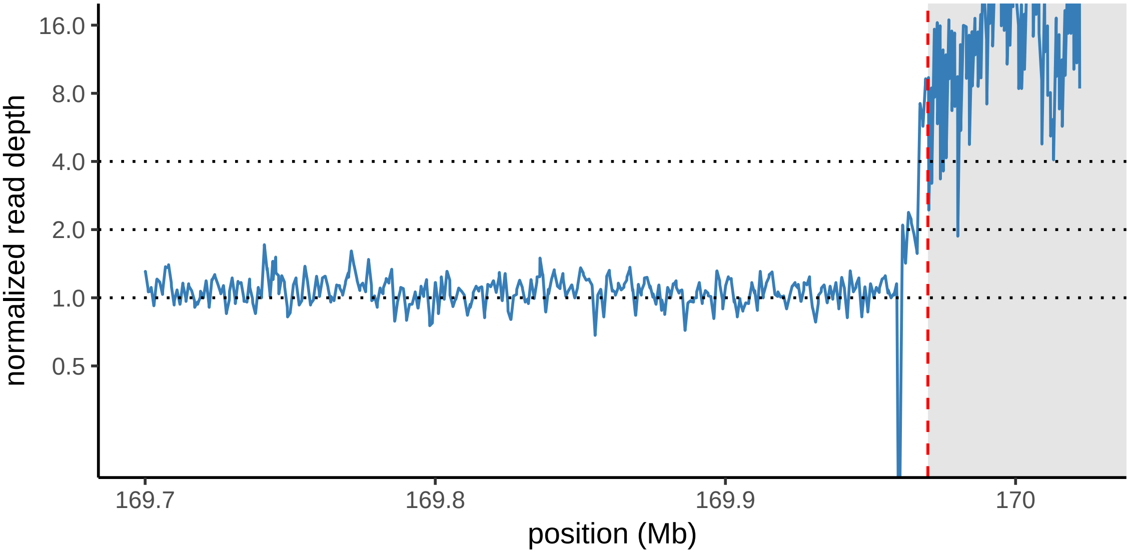

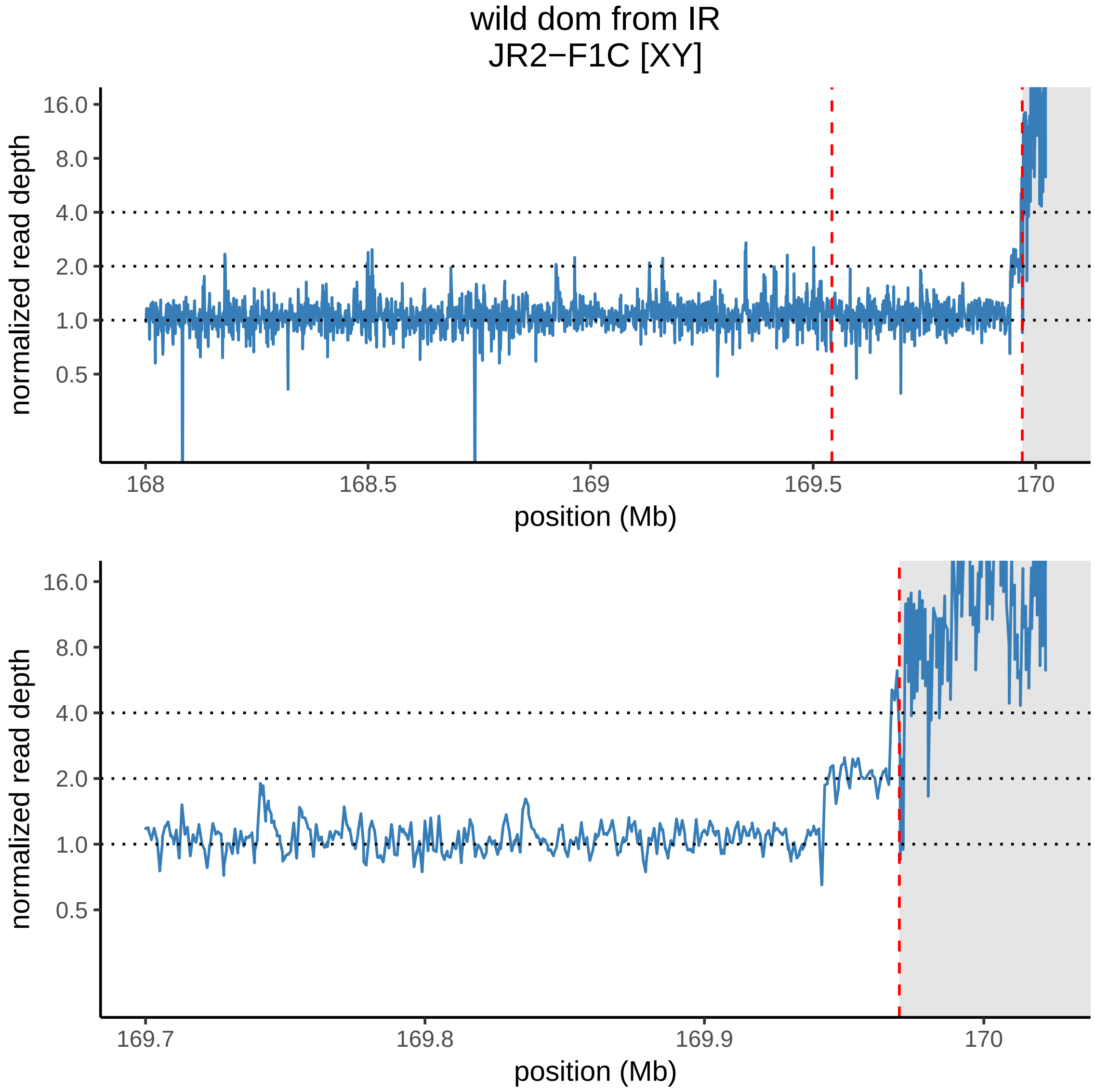
wild dom from IR
JR2−F1C [XY]
normalized read depth
16.0
8.0
4.0
2.0
1.0
0.5
168
168.5
169
169.5
170
position (Mb)
normalized read depth
16.0
8.0
4.0
2.0
1.0
0.5
169.7
169.8
169.9
170
position (Mb)

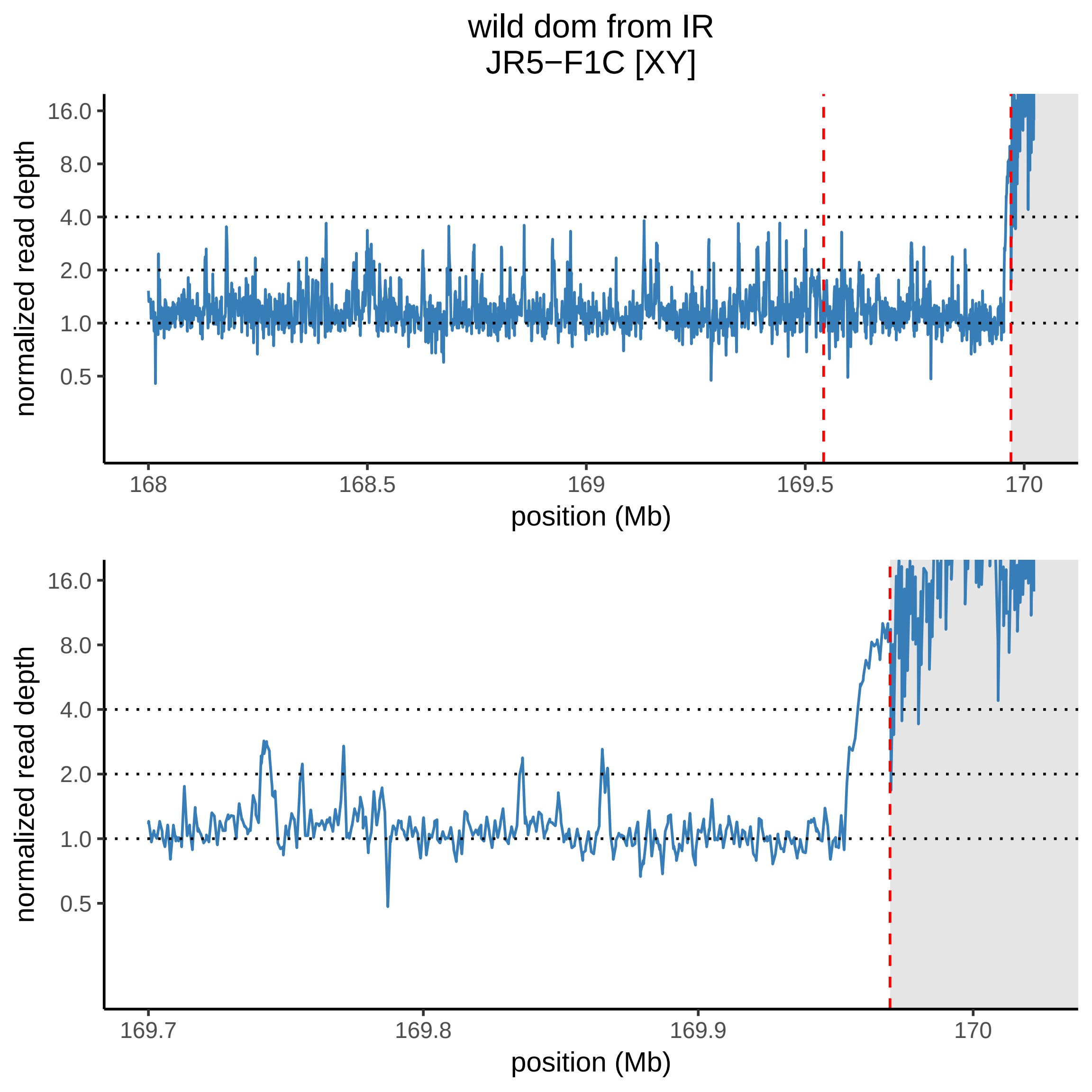
wild dom from IR
JR5−F1C [XY]
16.0
8.0
4.0
2.0
1.0
0.5
normalized read depth
168
168.5
169
169.5
170
position (Mb)
16.0
8.0
4.0
2.0
1.0
0.5
normalized read depth
169.7
169.8
169.9
170
position (Mb)

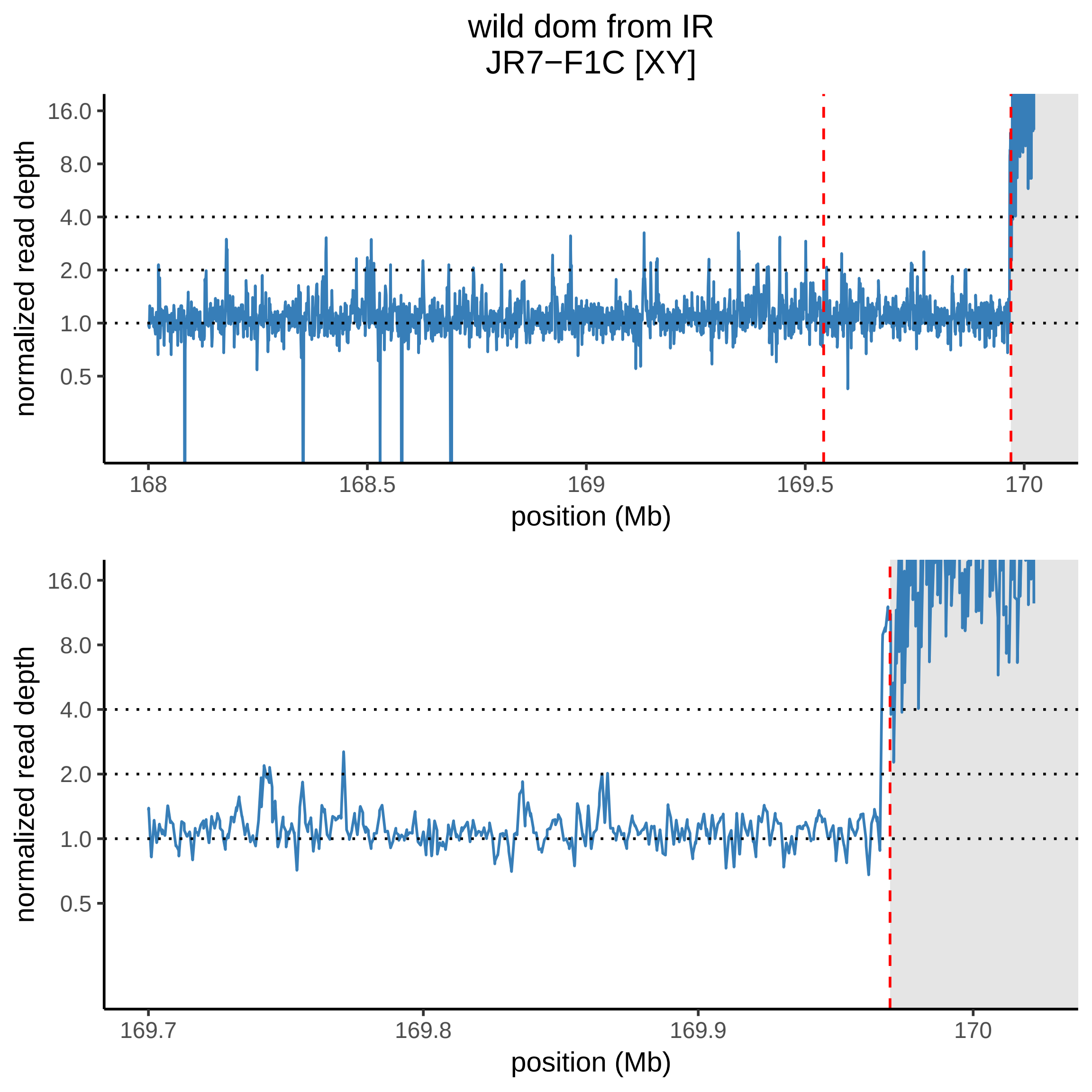

wild dom from IR
JR7−F1C [XY]
normalized read depth
16.0
8.0
4.0
2.0
1.0
0.5
168
168.5
169
169.5
170
position (Mb)
normalized read depth
16.0
8.0
4.0
2.0
1.0
0.5
169.7
169.8
169.9
170
position (Mb)

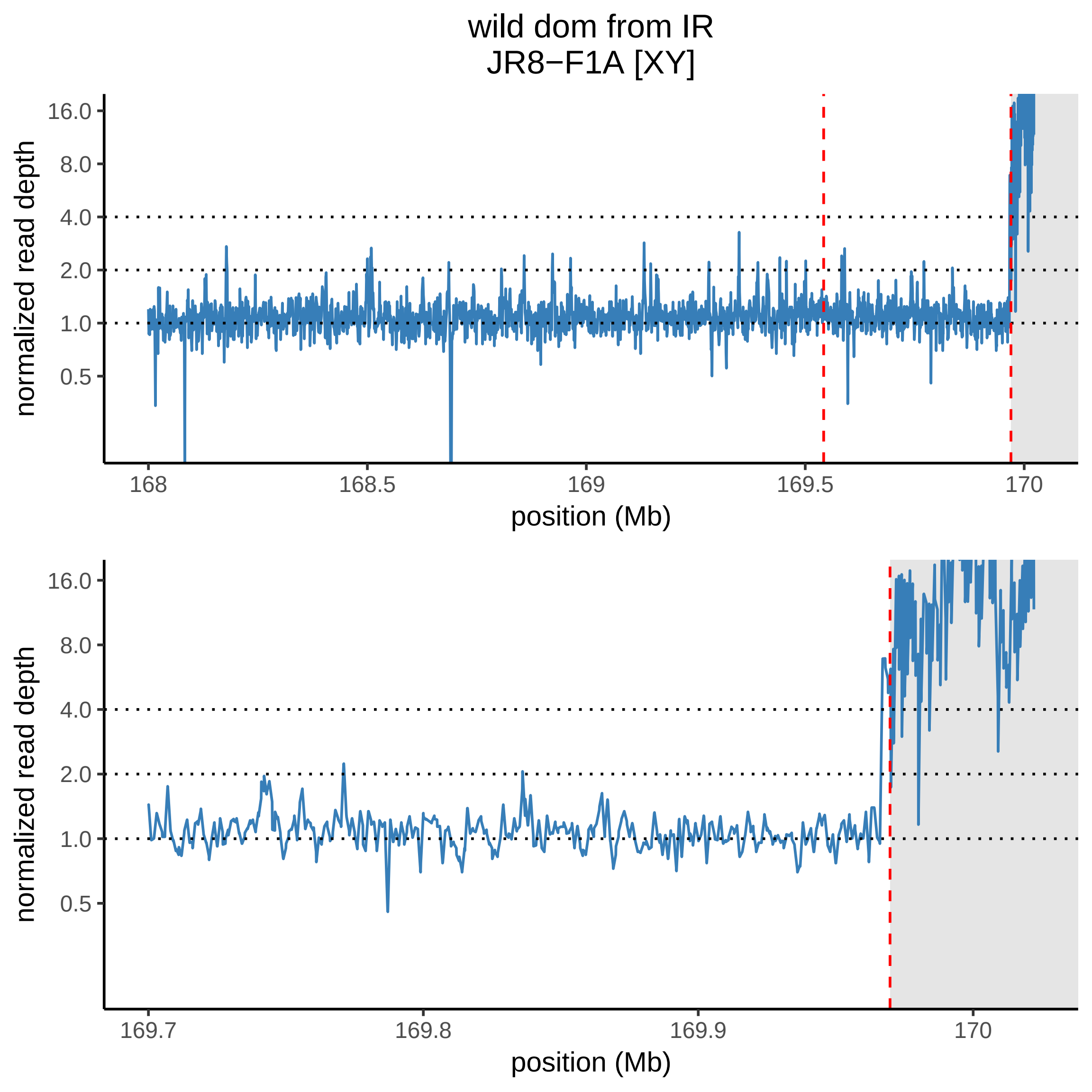

wild dom from IR
JR8−F1A [XY]
16.0
8.0
4.0
2.0
1.0
0.5
normalized read depth
168
168.5
169
169.5
170
position (Mb)
16.0
8.0
4.0
2.0
1.0
0.5
normalized read depth
169.7
169.8
169.9
170
position (Mb)

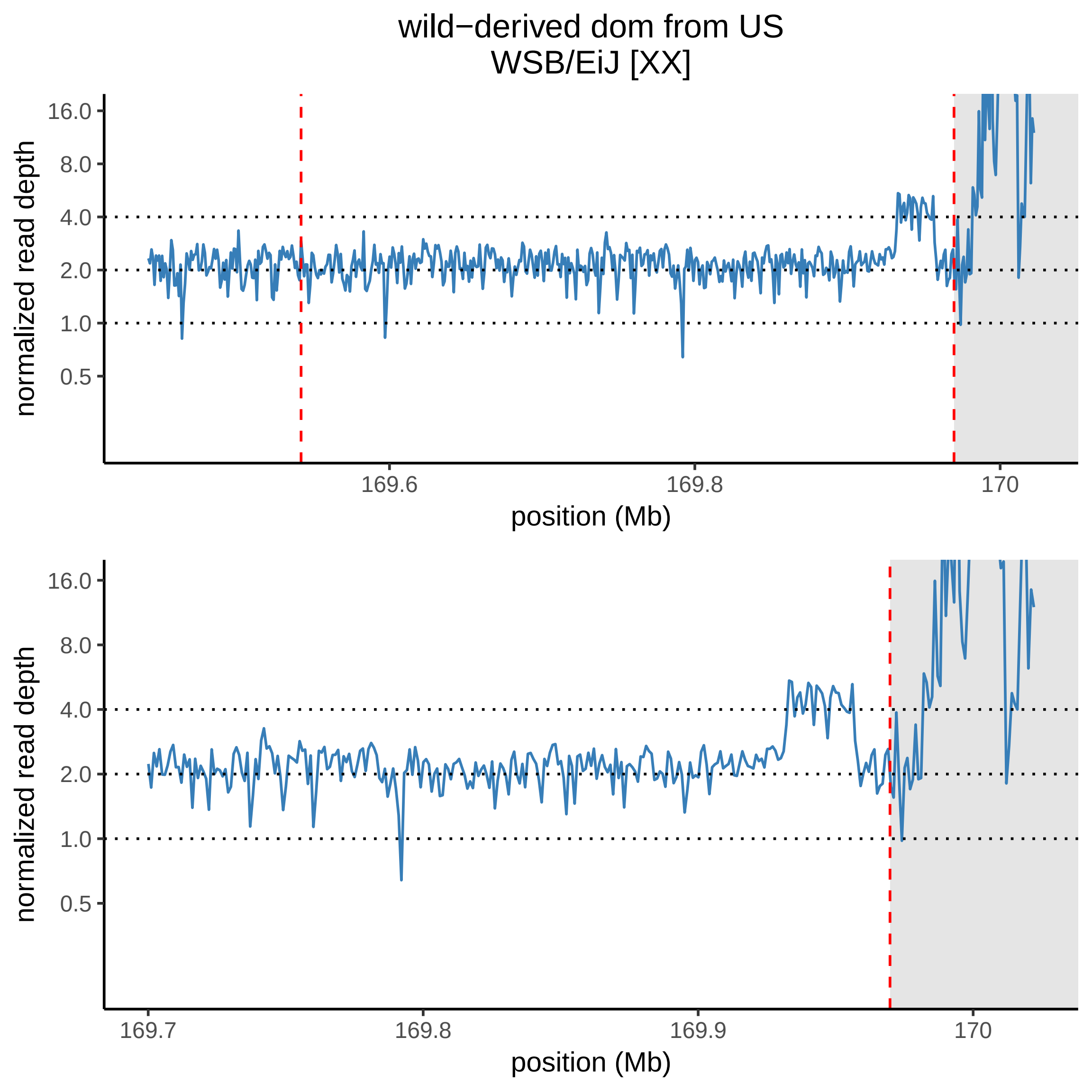
wild−derived dom from US
WSB/EiJ [XX]
normalized read depth
16.0
8.0
4.0
2.0
1.0
0.5
169.6
169.8
170
position (Mb)
normalized read depth
16.0
8.0
4.0
2.0
1.0
0.5
169.7
169.8
169.9
170
position (Mb)

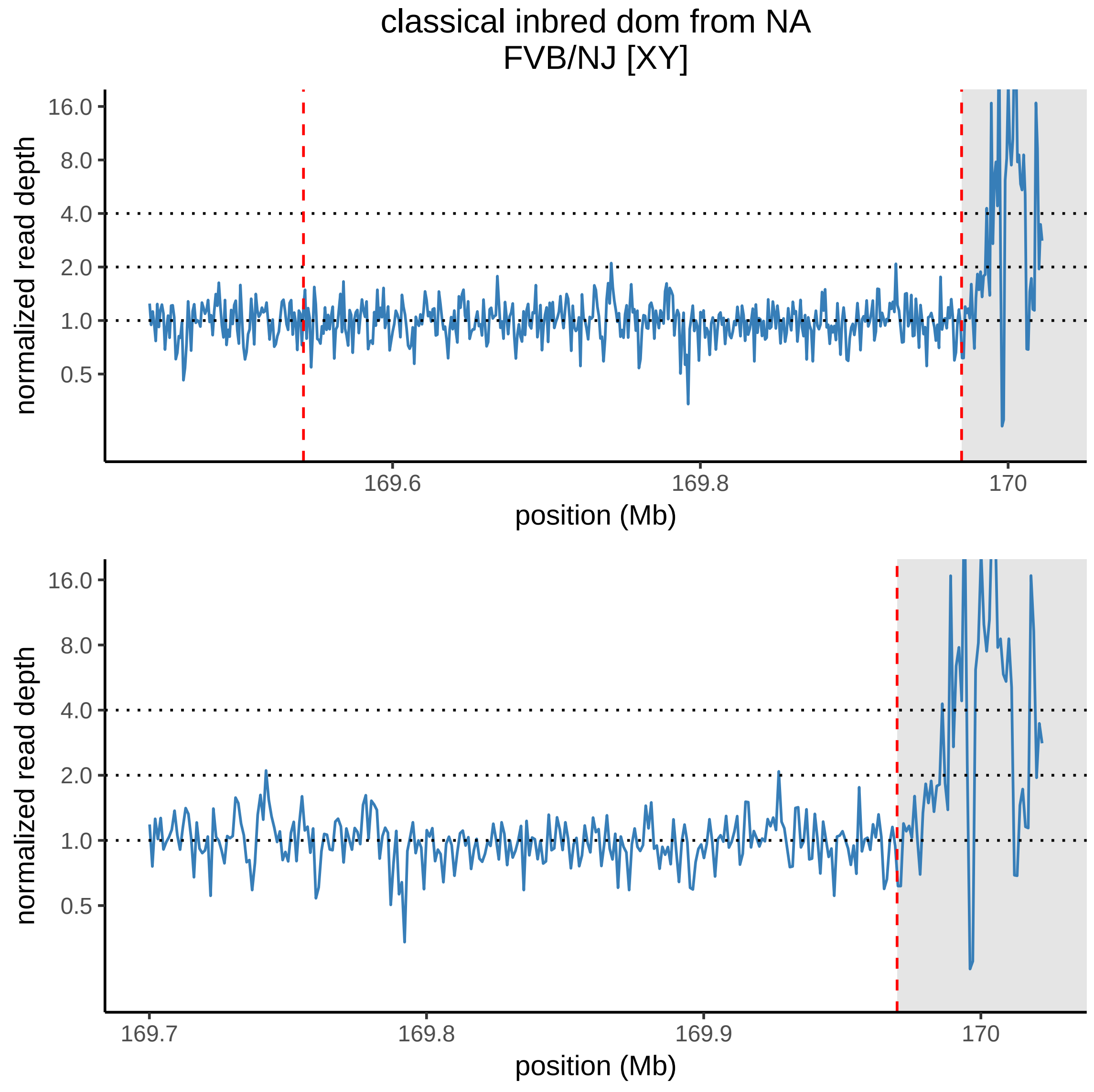
classical inbred dom from NA
FVB/NJ [XY]
normalized read depth
16.0
8.0
4.0
2.0
1.0
0.5
169.6
169.8
170
position (Mb)
normalized read depth
16.0
8.0
4.0
2.0
1.0
0.5
169.7
169.8
169.9
170
position (Mb)

wild mus from AF
AFG1 [XY]

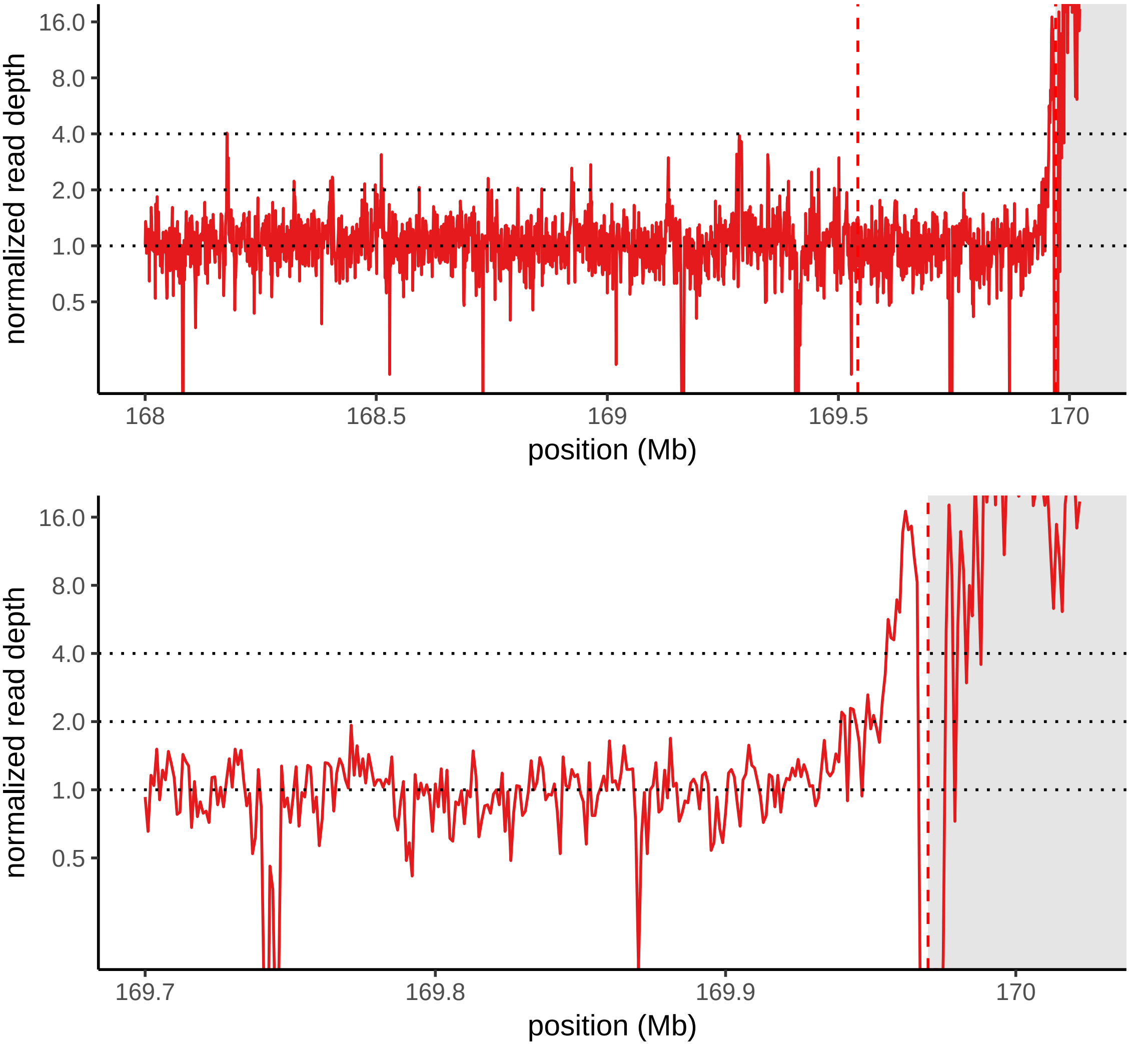

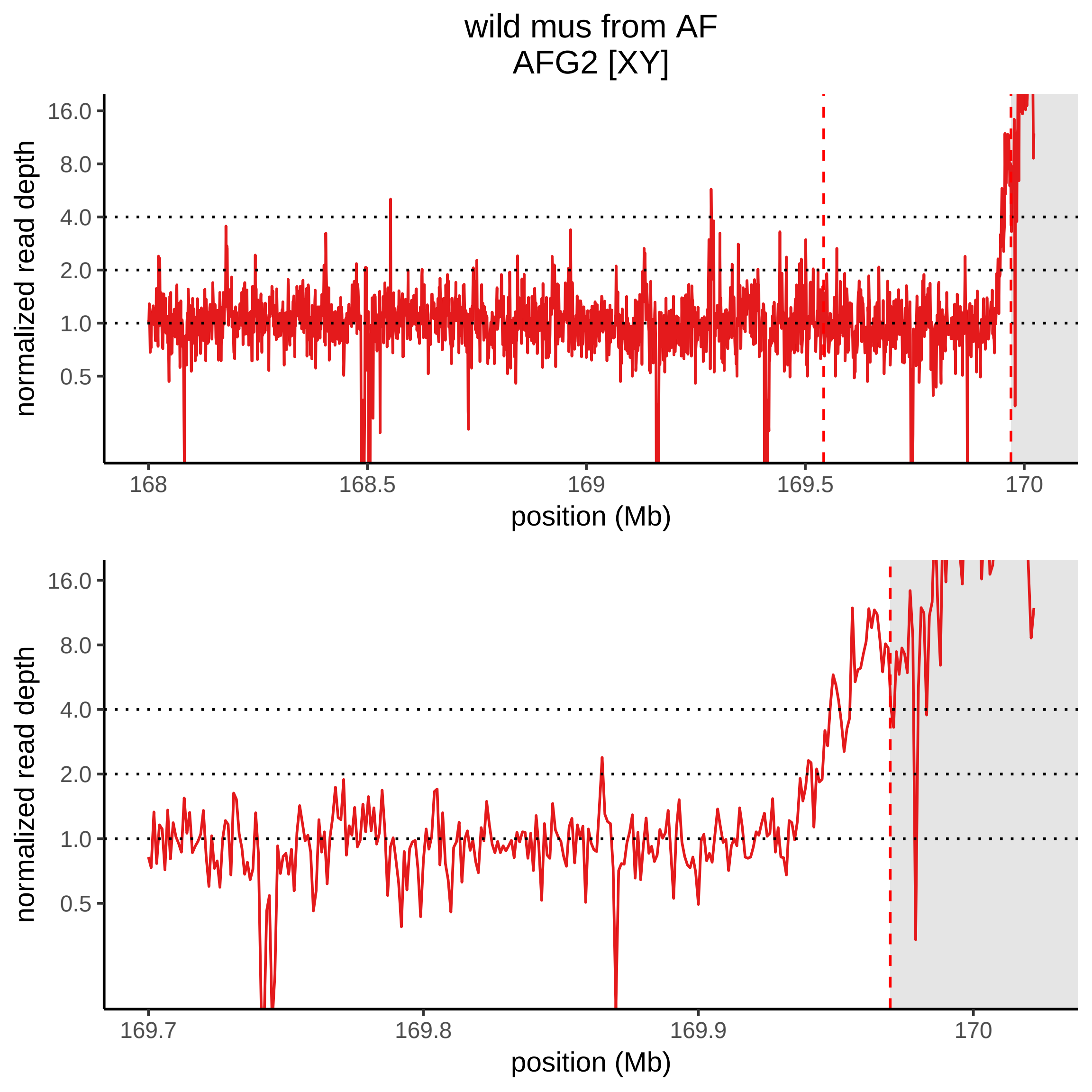
wild mus from AF
AFG2 [XY]
normalized read depth
16.0
8.0
4.0
2.0
1.0
0.5
168
168.5
169
169.5
170
position (Mb)
normalized read depth
16.0
8.0
4.0
2.0
1.0
0.5
169.7
169.8
169.9
170
position (Mb)

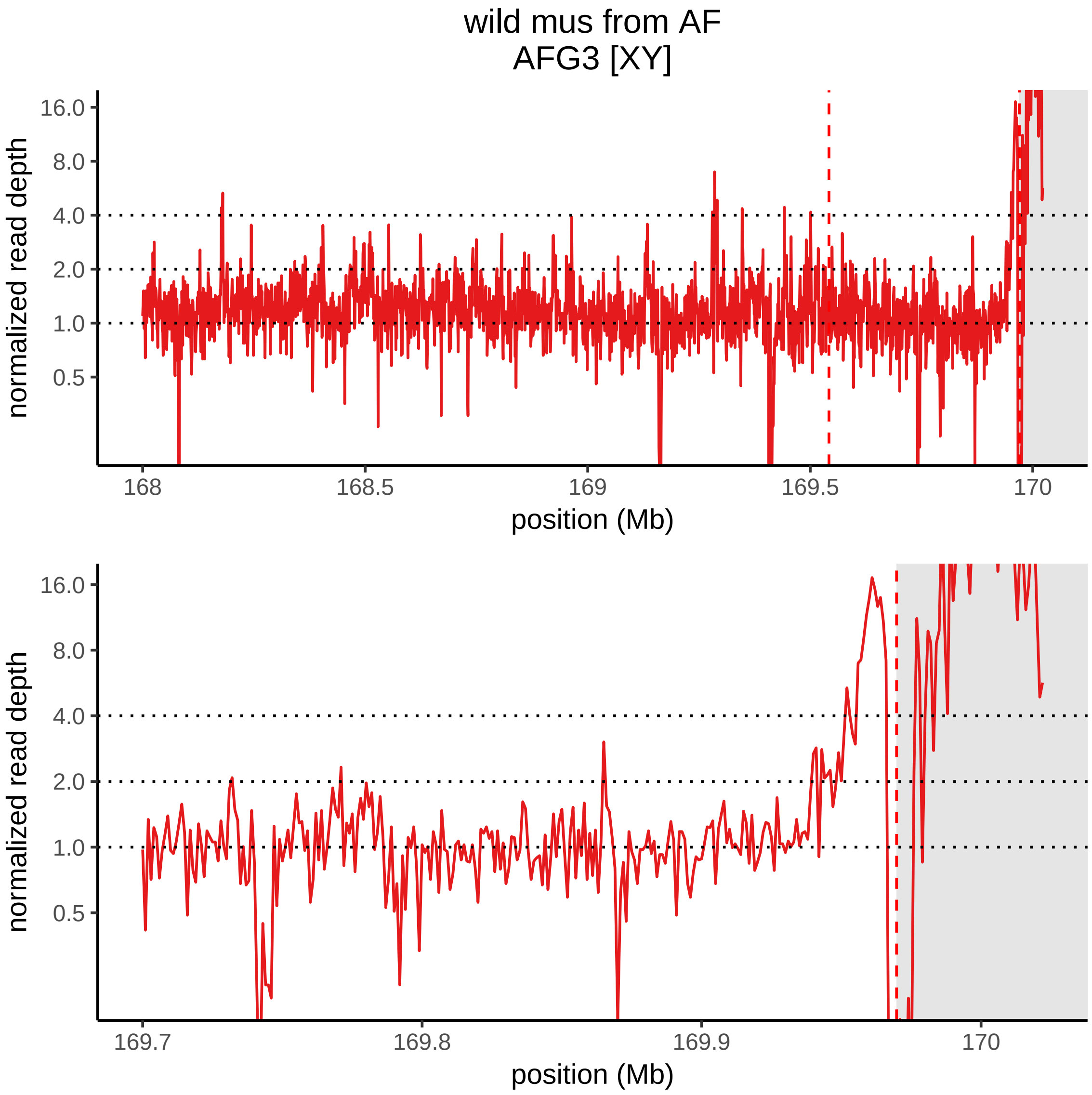

wild mus from AF
AFG3 [XY]
normalized read depth
16.0
8.0
4.0
2.0
1.0
0.5
168
168.5
169
169.5
170
position (Mb)
normalized read depth
16.0
8.0
4.0
2.0
1.0
0.5
169.7
169.8
169.9
170
position (Mb)

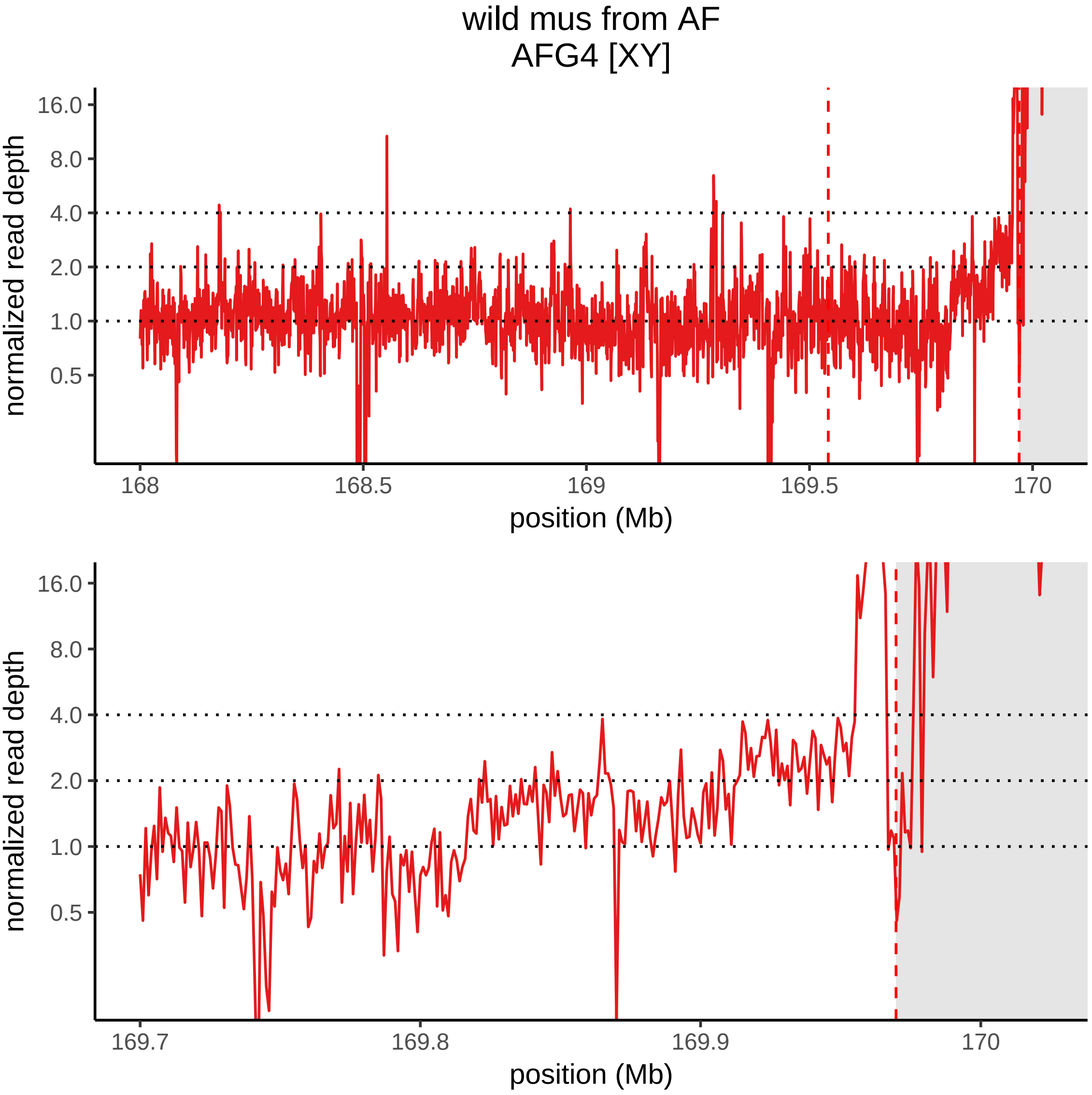
wild mus from AF
AFG4 [XY]
normalized read depth
16.0
8.0
4.0
2.0
1.0
0.5
168
168.5
169
169.5
170
position (Mb)
normalized read depth
16.0
8.0
4.0
2.0
1.0
0.5
169.7
169.8
169.9
170
position (Mb)

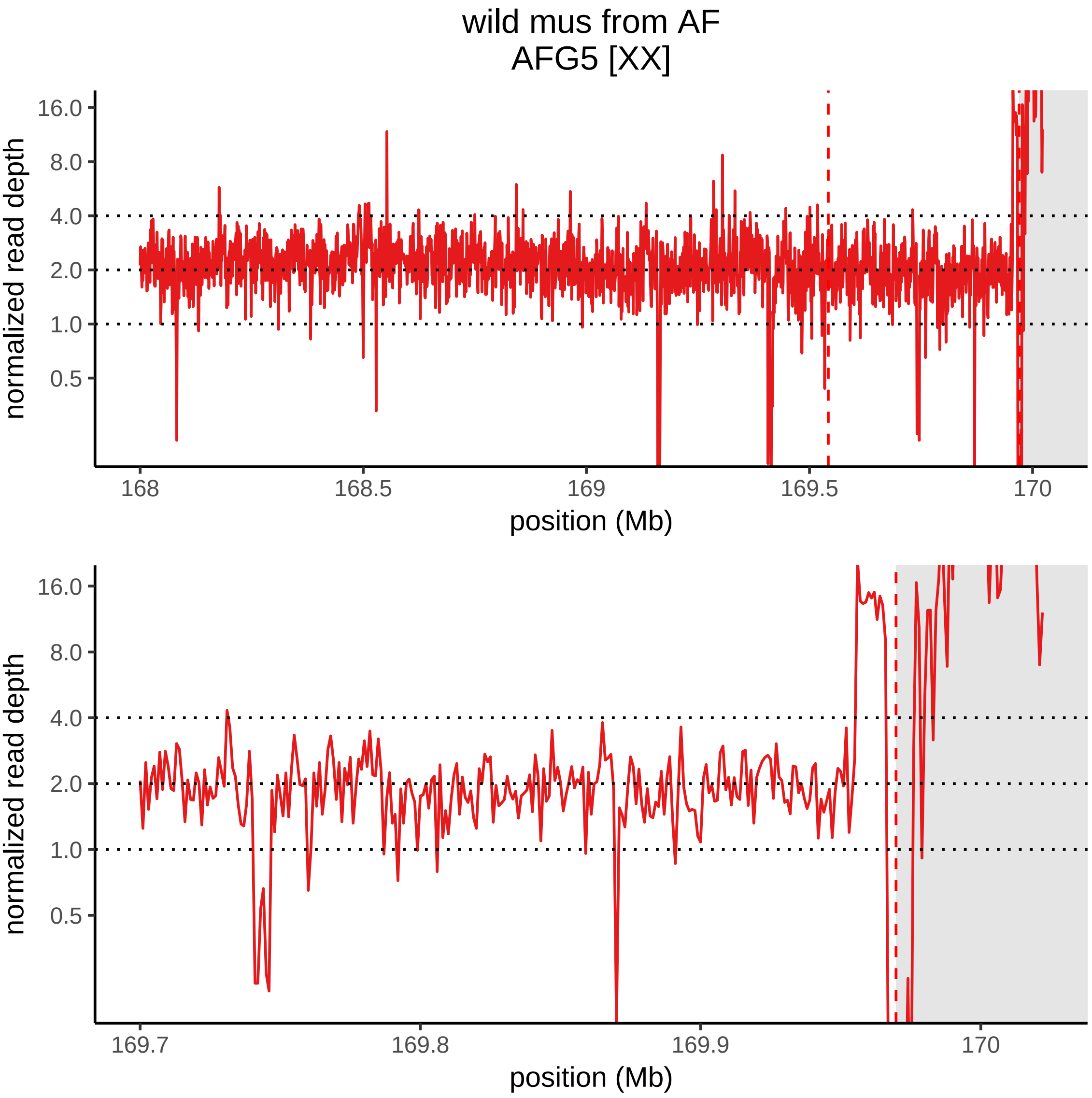

wild mus from AF
AFG5 [XX]
normalized read depth
16.0
8.0
4.0
2.0
1.0
0.5
168
168.5
169
169.5
170
position (Mb)
normalized read depth
16.0
8.0
4.0
2.0
1.0
0.5
169.7
169.8
169.9
170
position (Mb)

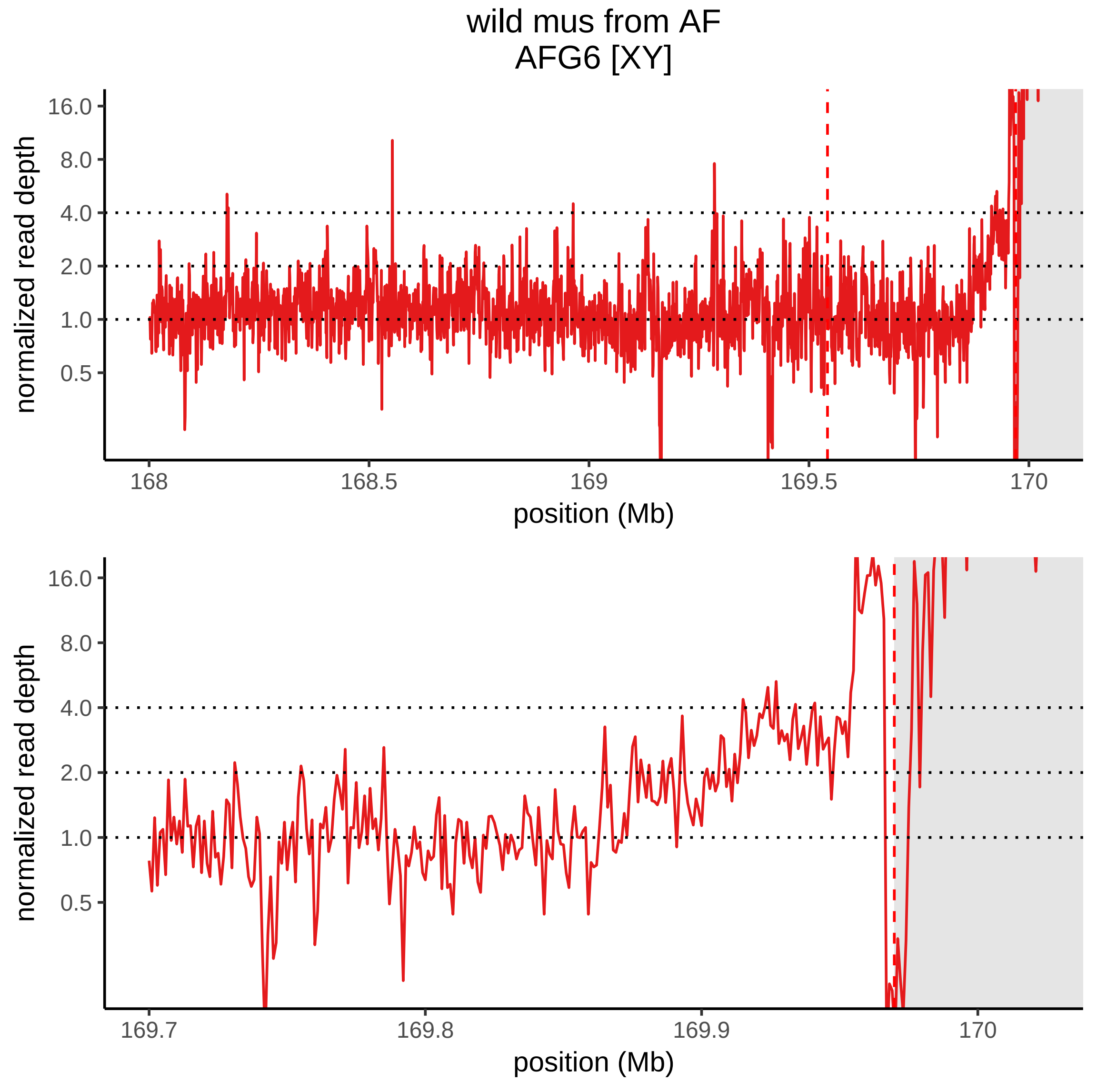
wild mus from AF
AFG6 [XY]
normalized read depth
16.0
8.0
4.0
2.0
1.0
0.5
168
168.5
169
169.5
170
position (Mb)
normalized read depth
16.0
8.0
4.0
2.0
1.0
0.5
169.7
169.8
169.9
170
position (Mb)

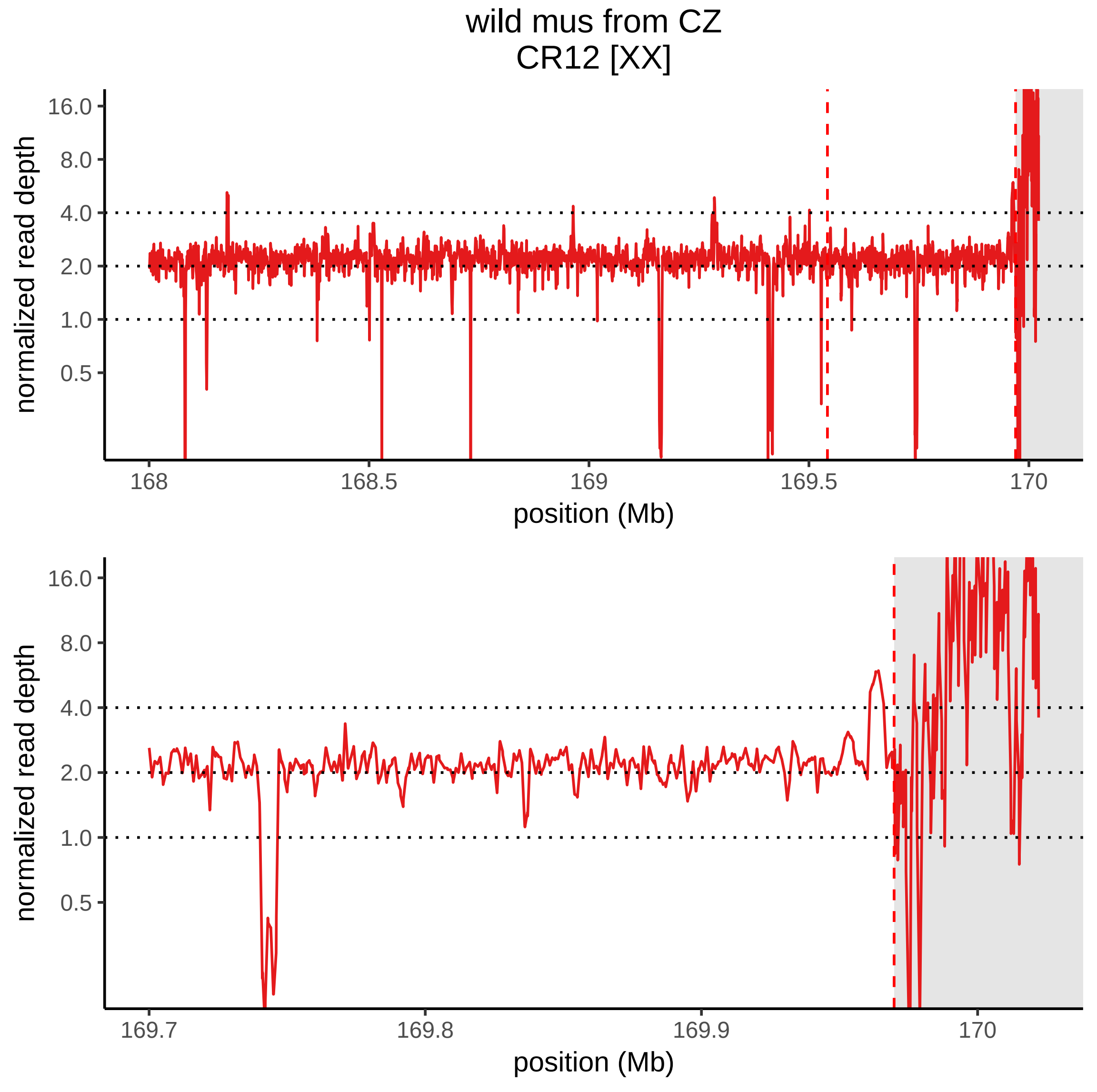
wild mus from CZ
CR12 [XX]
normalized read depth
16.0
8.0
4.0
2.0
1.0
0.5
168
168.5
169
169.5
170
position (Mb)
normalized read depth
16.0
8.0
4.0
2.0
1.0
0.5
169.7
169.8
169.9
170
position (Mb)

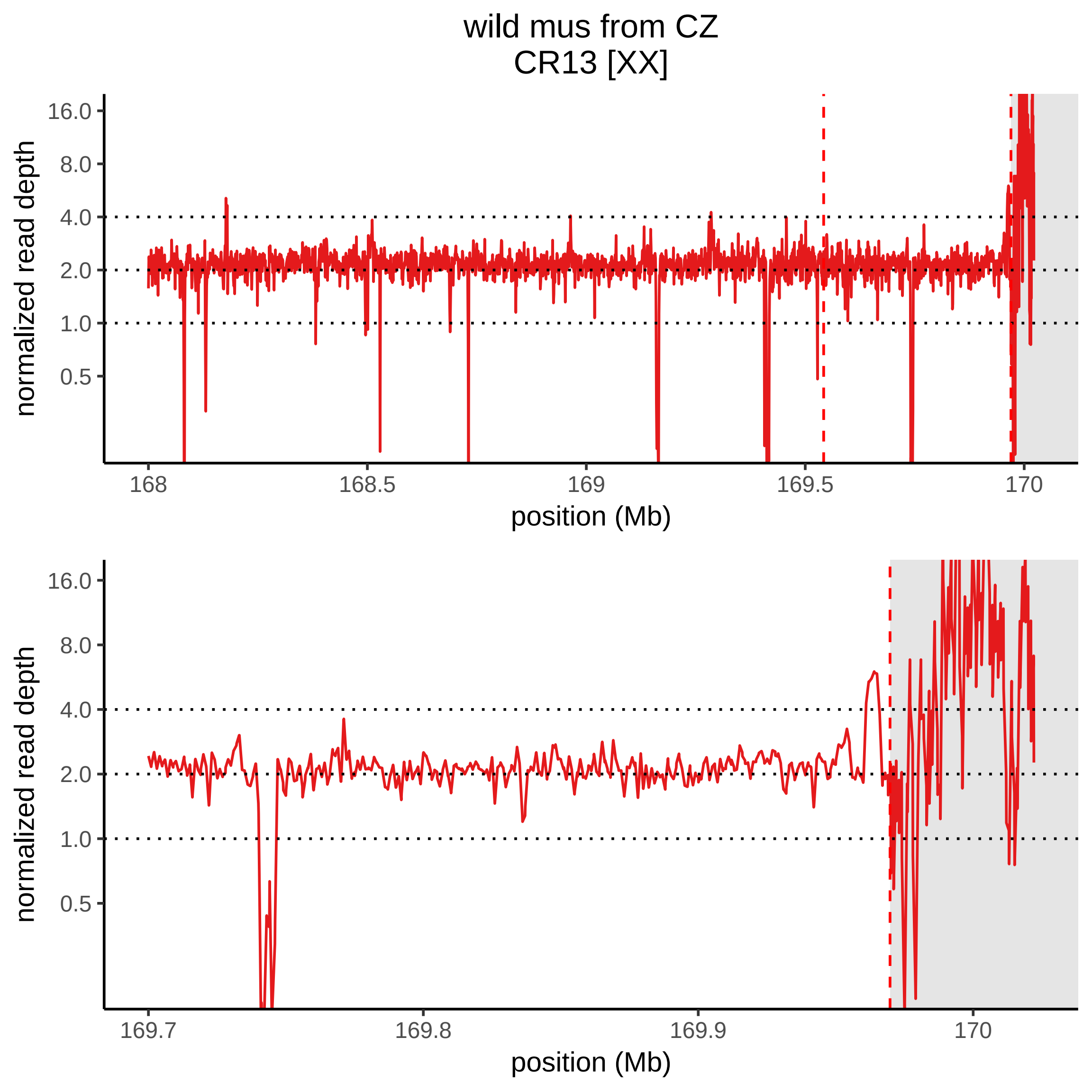

wild mus from CZ
CR13 [XX]
normalized read depth
16.0
8.0
4.0
2.0
1.0
0.5
168
168.5
169
169.5
170
position (Mb)
normalized read depth
16.0
8.0
4.0
2.0
1.0
0.5
169.7
169.8
169.9
170
position (Mb)

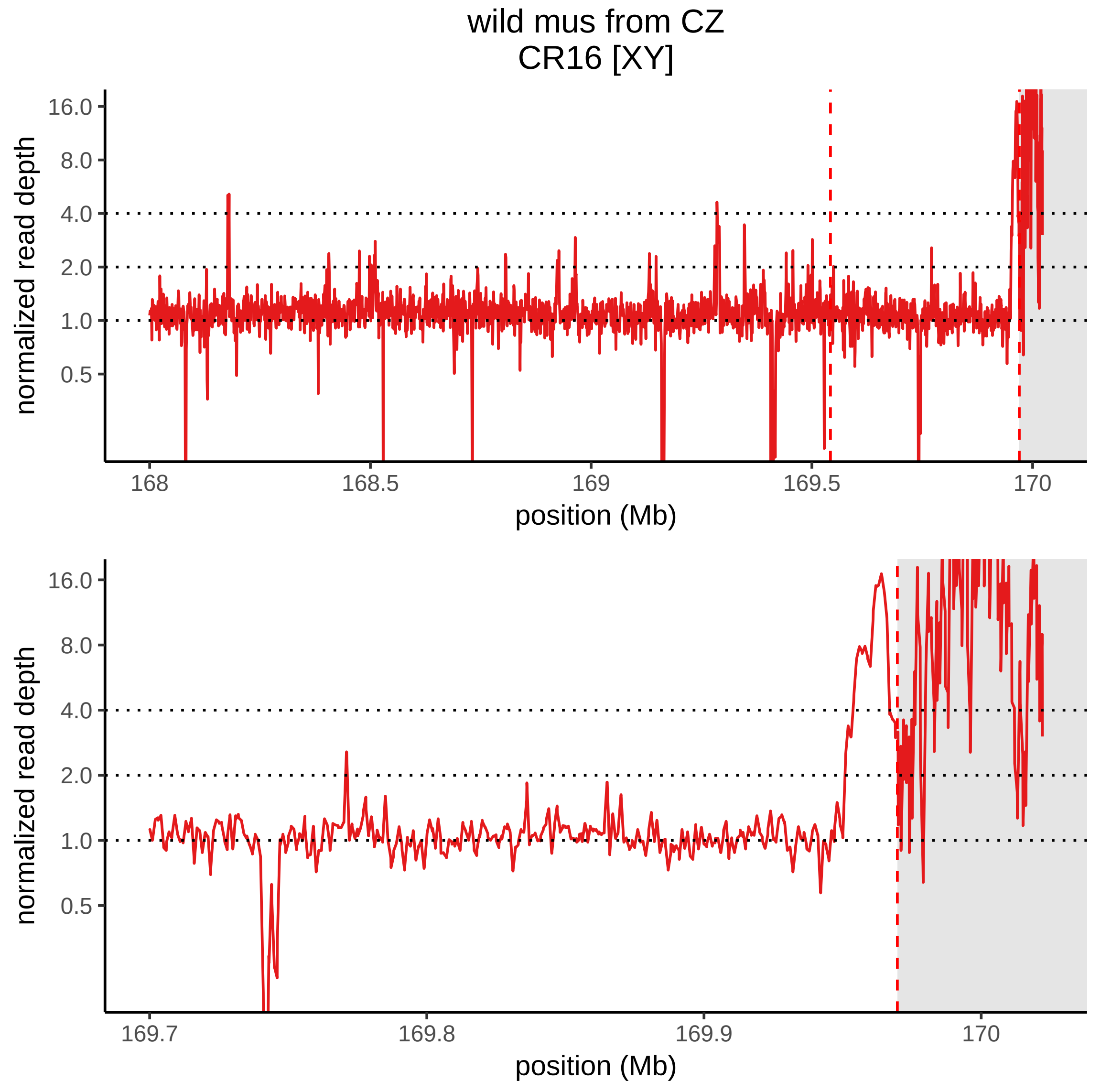

wild mus from CZ
CR16 [XY]
normalized read depth
16.0
8.0
4.0
2.0
1.0
0.5
168
168.5
169
169.5
170
position (Mb)
normalized read depth
16.0
8.0
4.0
2.0
1.0
0.5
169.7
169.8
169.9
170
position (Mb)

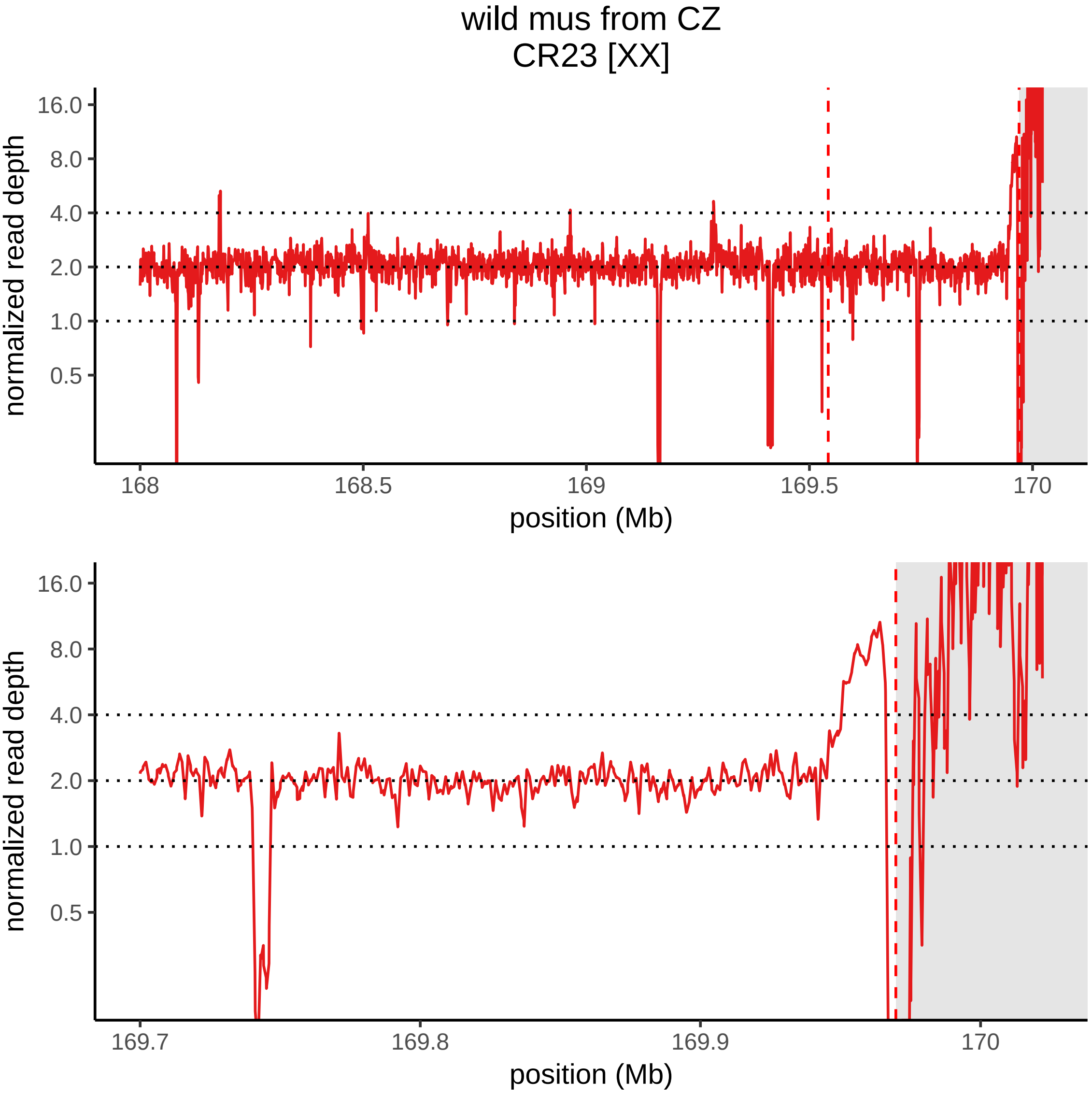

wild mus from CZ
CR23 [XX]
normalized read depth
16.0
8.0
4.0
2.0
1.0
0.5
168
168.5
169
169.5
170
position (Mb)
normalized read depth
16.0
8.0
4.0
2.0
1.0
0.5
169.7
169.8
169.9
170
position (Mb)

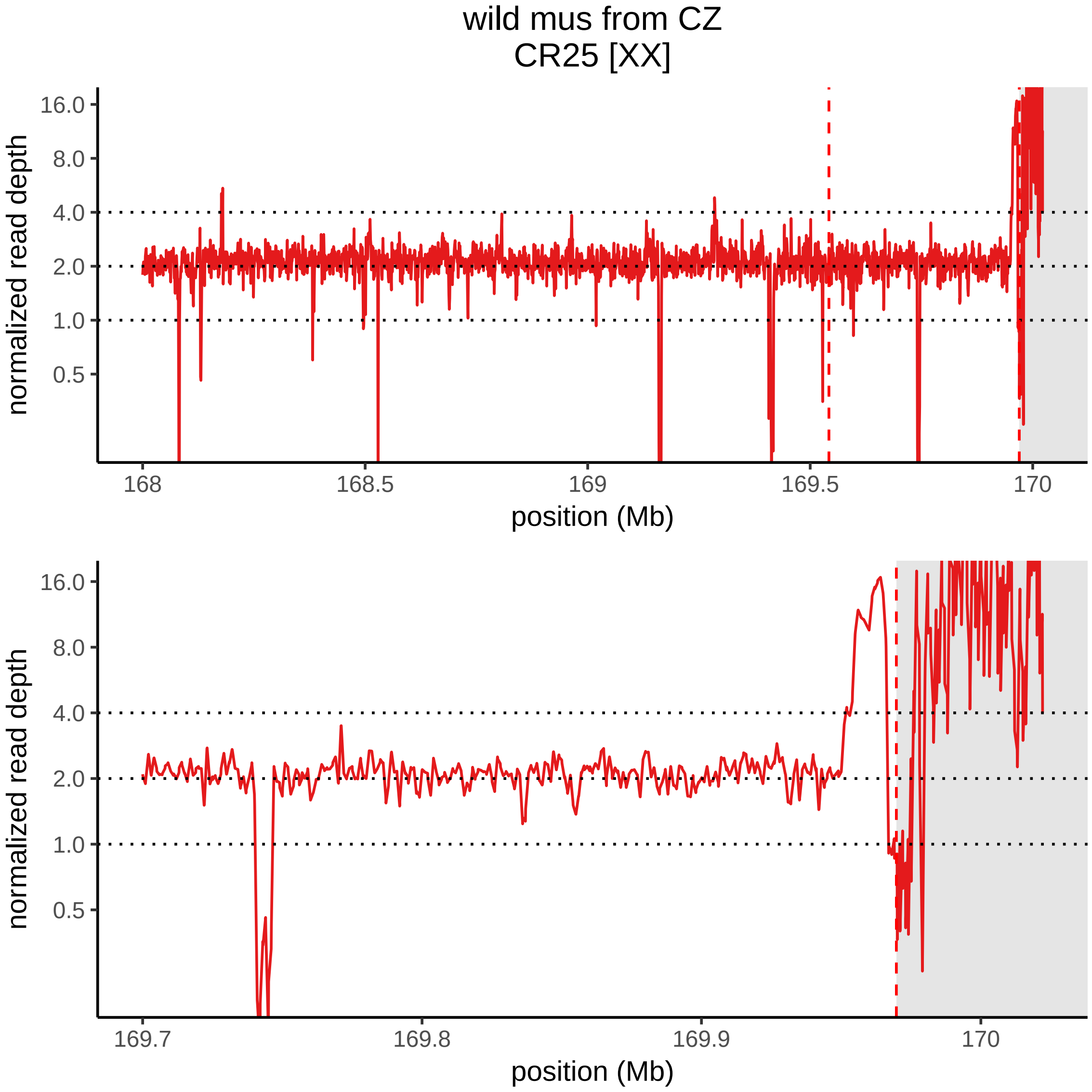

wild mus from CZ
CR25 [XX]
normalized read depth
16.0
8.0
4.0
2.0
1.0
0.5
168
168.5
169
169.5
170
position (Mb)
normalized read depth
16.0
8.0
4.0
2.0
1.0
0.5
169.7
169.8
169.9
170
position (Mb)

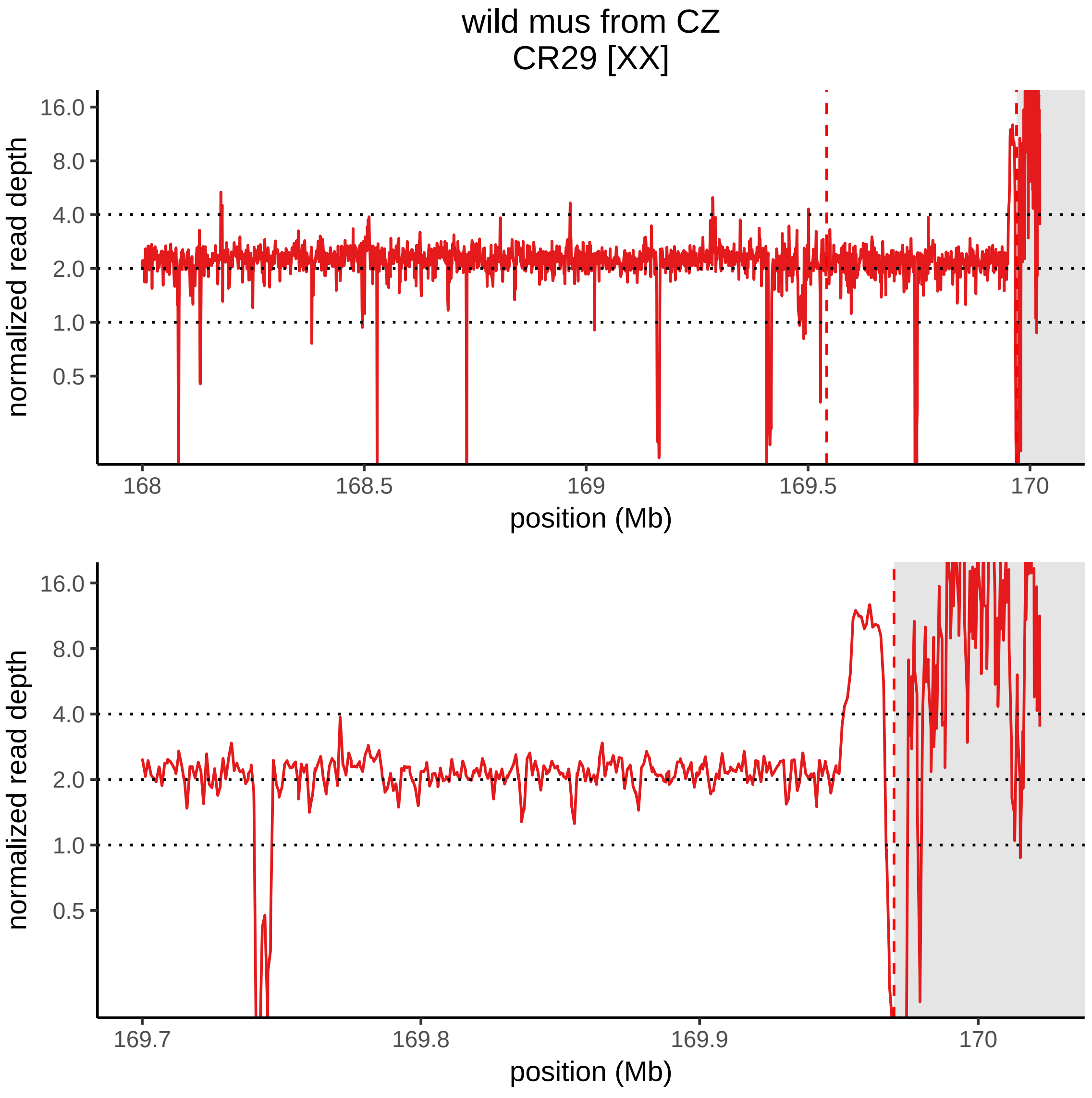
wild mus from CZ
CR29 [XX]
normalized read depth
16.0
8.0
4.0
2.0
1.0
0.5
168
168.5
169
169.5
170
position (Mb)
normalized read depth
16.0
8.0
4.0
2.0
1.0
0.5
169.7
169.8
169.9
170
position (Mb)

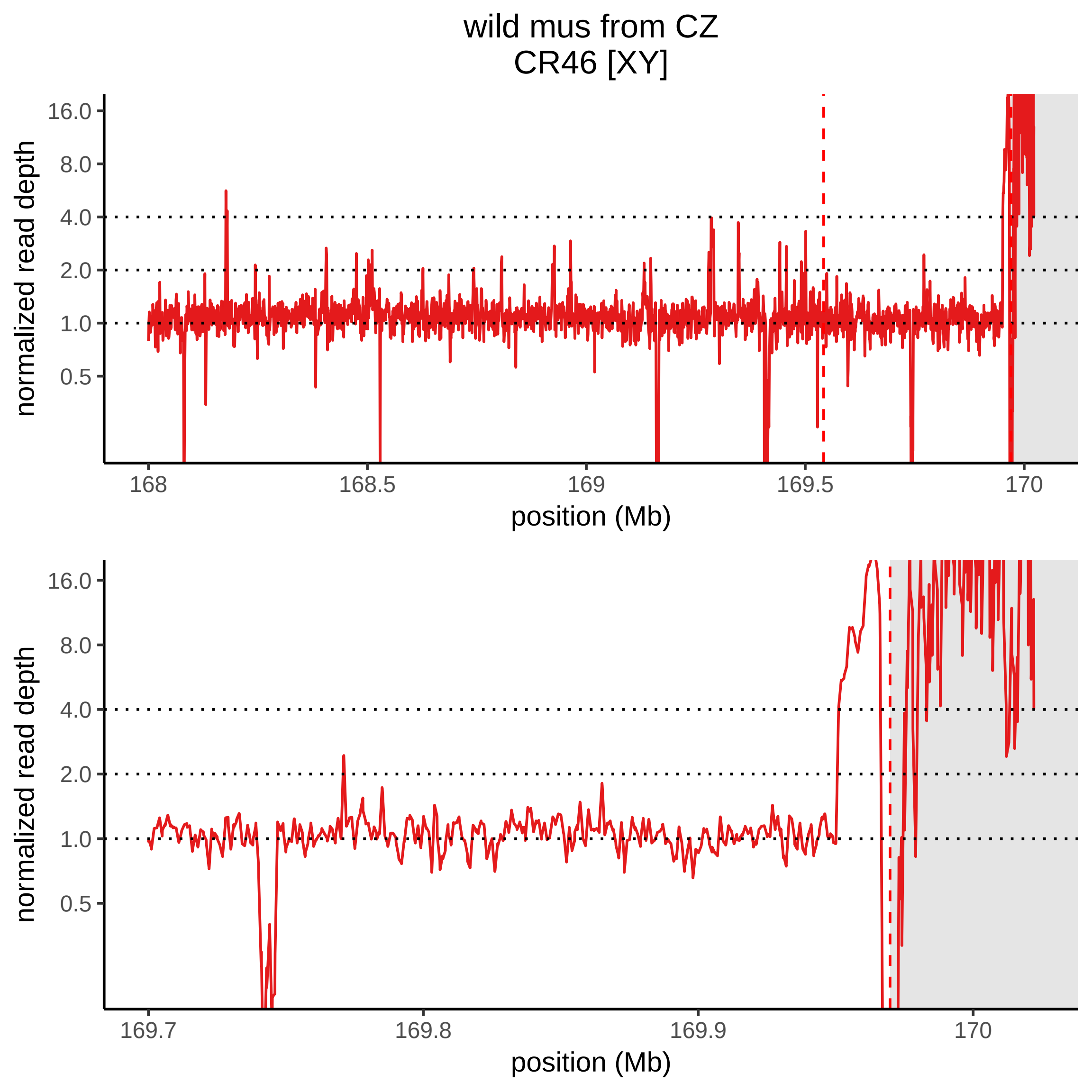

wild mus from CZ
CR46 [XY]
normalized read depth
16.0
8.0
4.0
2.0
1.0
0.5
168
168.5
169
169.5
170
position (Mb)
normalized read depth
16.0
8.0
4.0
2.0
1.0
0.5
169.7
169.8
169.9
170
position (Mb)

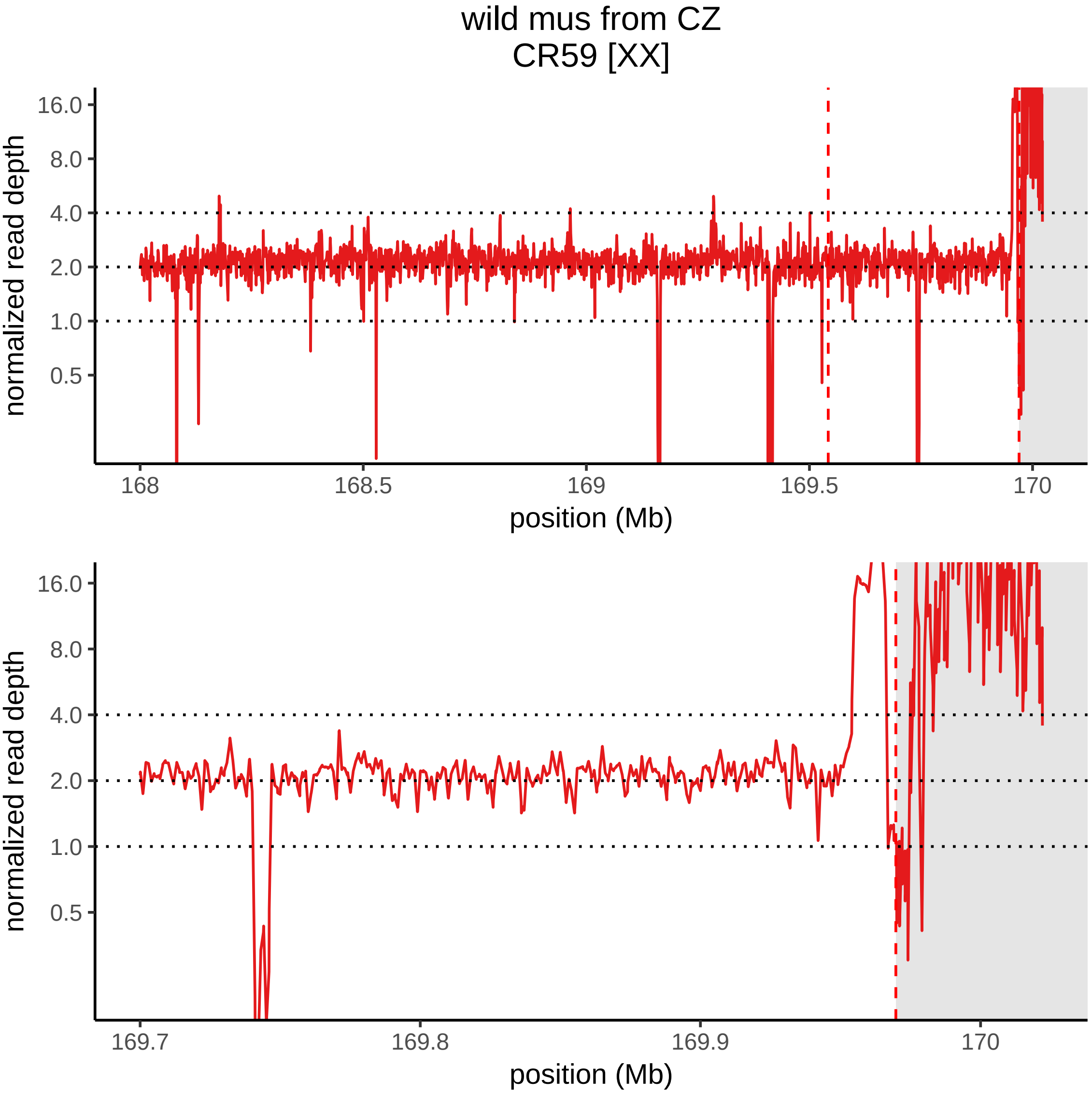
wild mus from CZ
CR59 [XX]
normalized read depth
16.0
8.0
4.0
2.0
1.0
0.5
168
168.5
169
169.5
170
position (Mb)
normalized read depth
16.0
8.0
4.0
2.0
1.0
0.5
169.7
169.8
169.9
170
position (Mb)

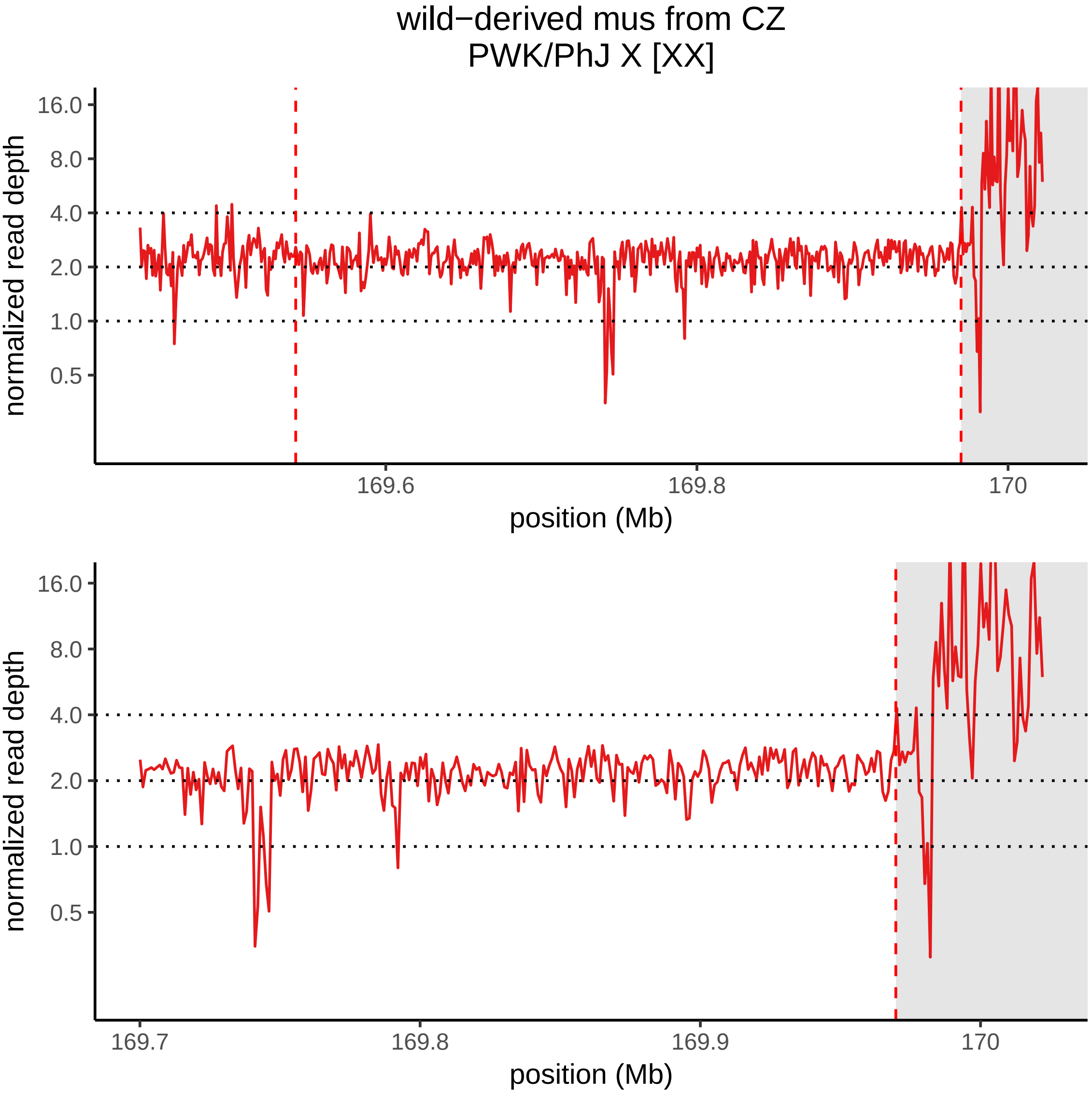
wild−derived mus from CZ
PWK/PhJ X [XX]
normalized read depth
16.0
8.0
4.0
2.0
1.0
0.5
169.6
169.8
170
position (Mb)
normalized read depth
16.0
8.0
4.0
2.0
1.0
0.5
169.7
169.8
169.9
170
position (Mb)

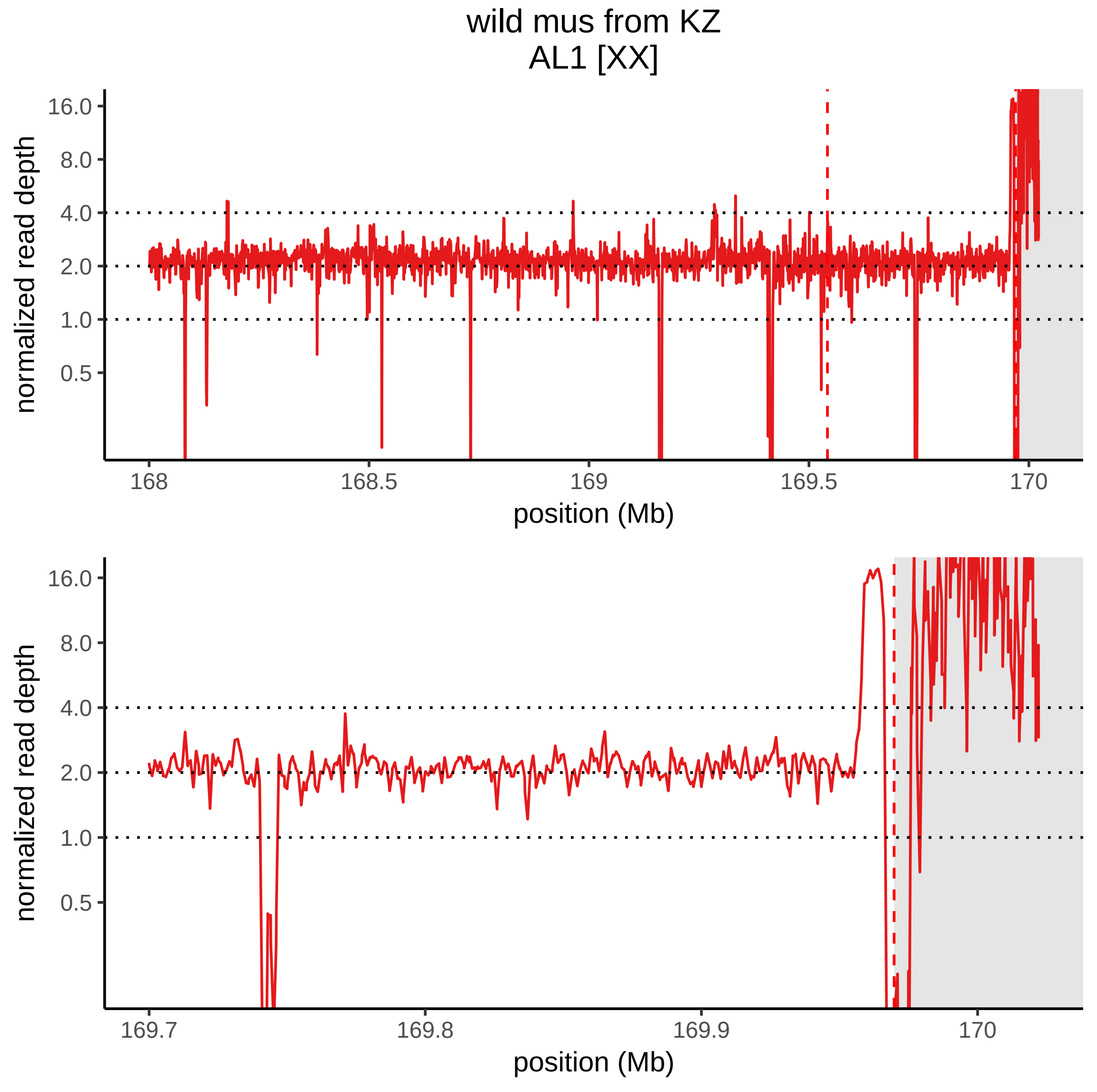

wild mus from KZ
AL1 [XX]
normalized read depth
16.0
8.0
4.0
2.0
1.0
0.5
168
168.5
169
169.5
170
position (Mb)
normalized read depth
16.0
8.0
4.0
2.0
1.0
0.5
169.7
169.8
169.9
170
position (Mb)

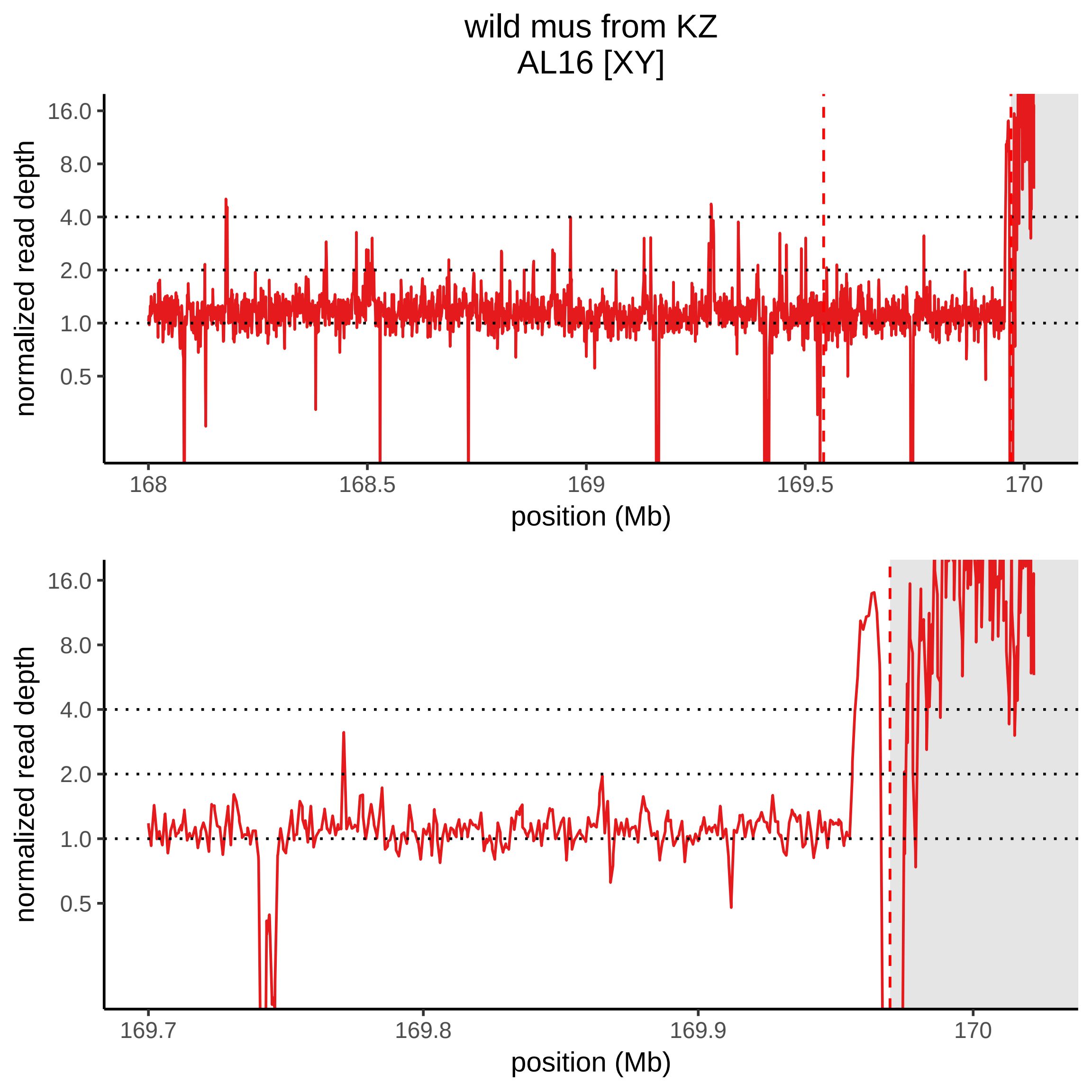
wild mus from KZ
AL16 [XY]
normalized read depth
16.0
8.0
4.0
2.0
1.0
0.5
168
168.5
169
169.5
170
position (Mb)
normalized read depth
16.0
8.0
4.0
2.0
1.0
0.5
169.7
169.8
169.9
170
position (Mb)

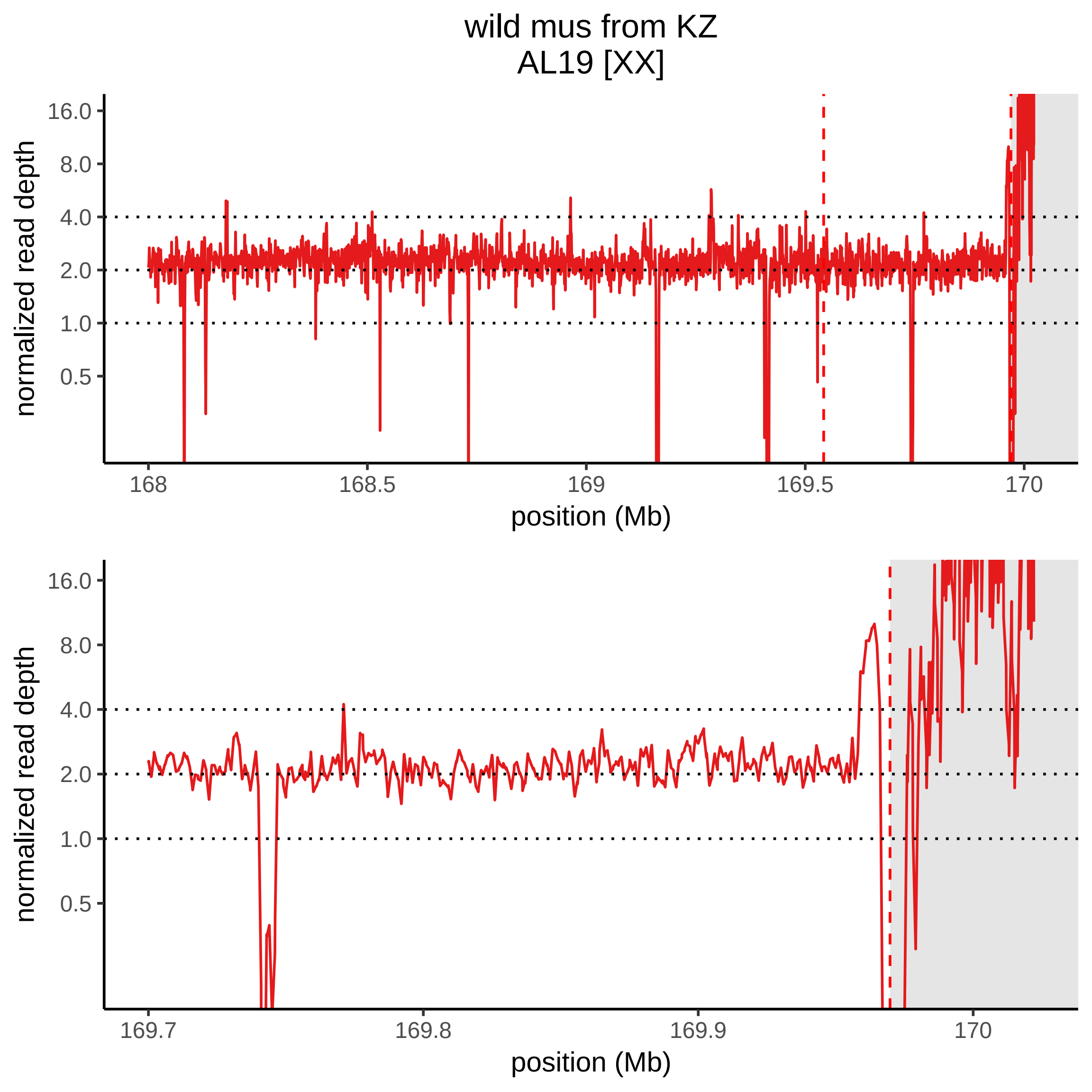
wild mus from KZ
AL19 [XX]
normalized read depth
16.0
8.0
4.0
2.0
1.0
0.5
168
168.5
169
169.5
170
position (Mb)
normalized read depth
16.0
8.0
4.0
2.0
1.0
0.5
169.7
169.8
169.9
170
position (Mb)

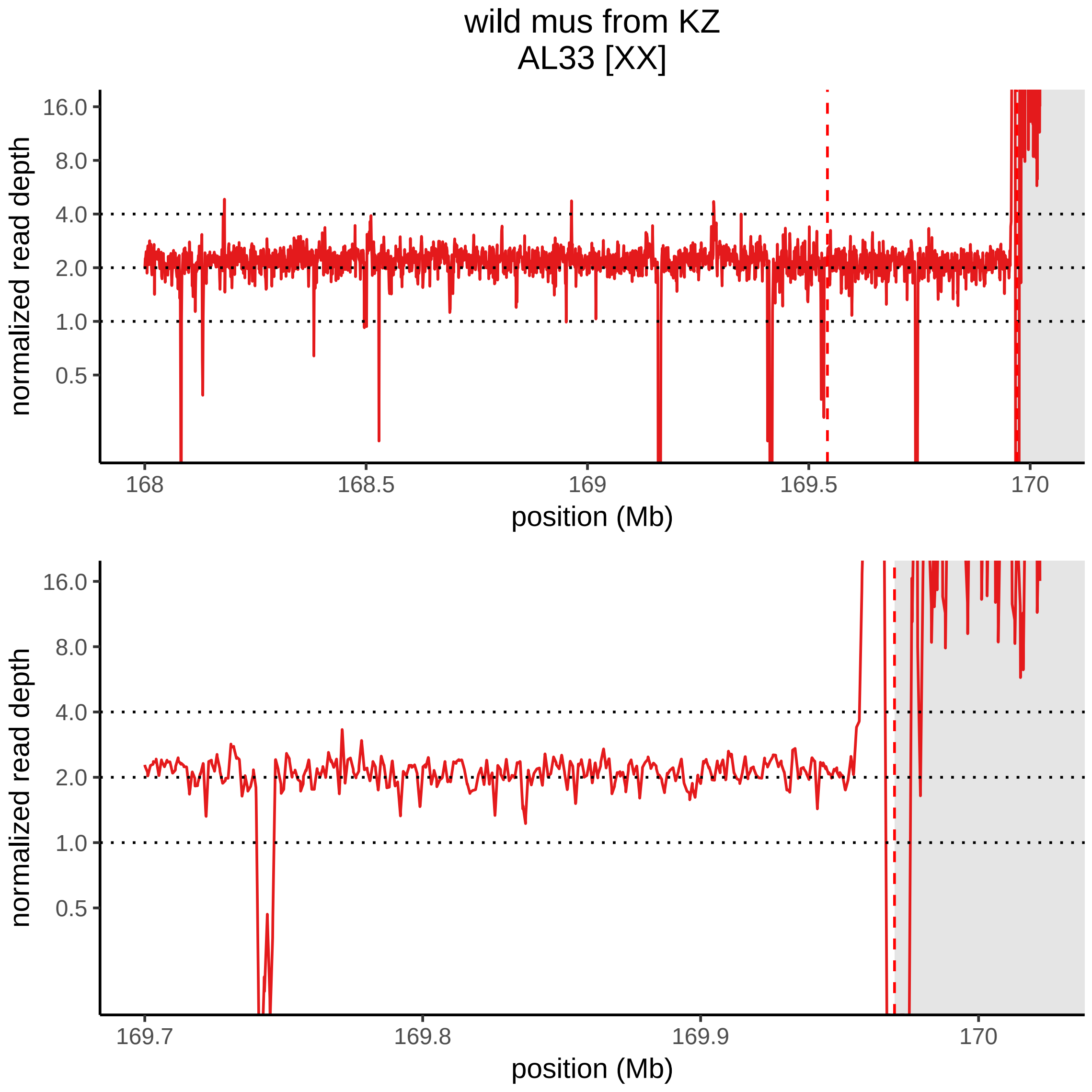
wild mus from KZ
AL33 [XX]
normalized read depth
16.0
8.0
4.0
2.0
1.0
0.5
168
168.5
169
169.5
170
position (Mb)
normalized read depth
16.0
8.0
4.0
2.0
1.0
0.5
169.7
169.8
169.9
170
position (Mb)

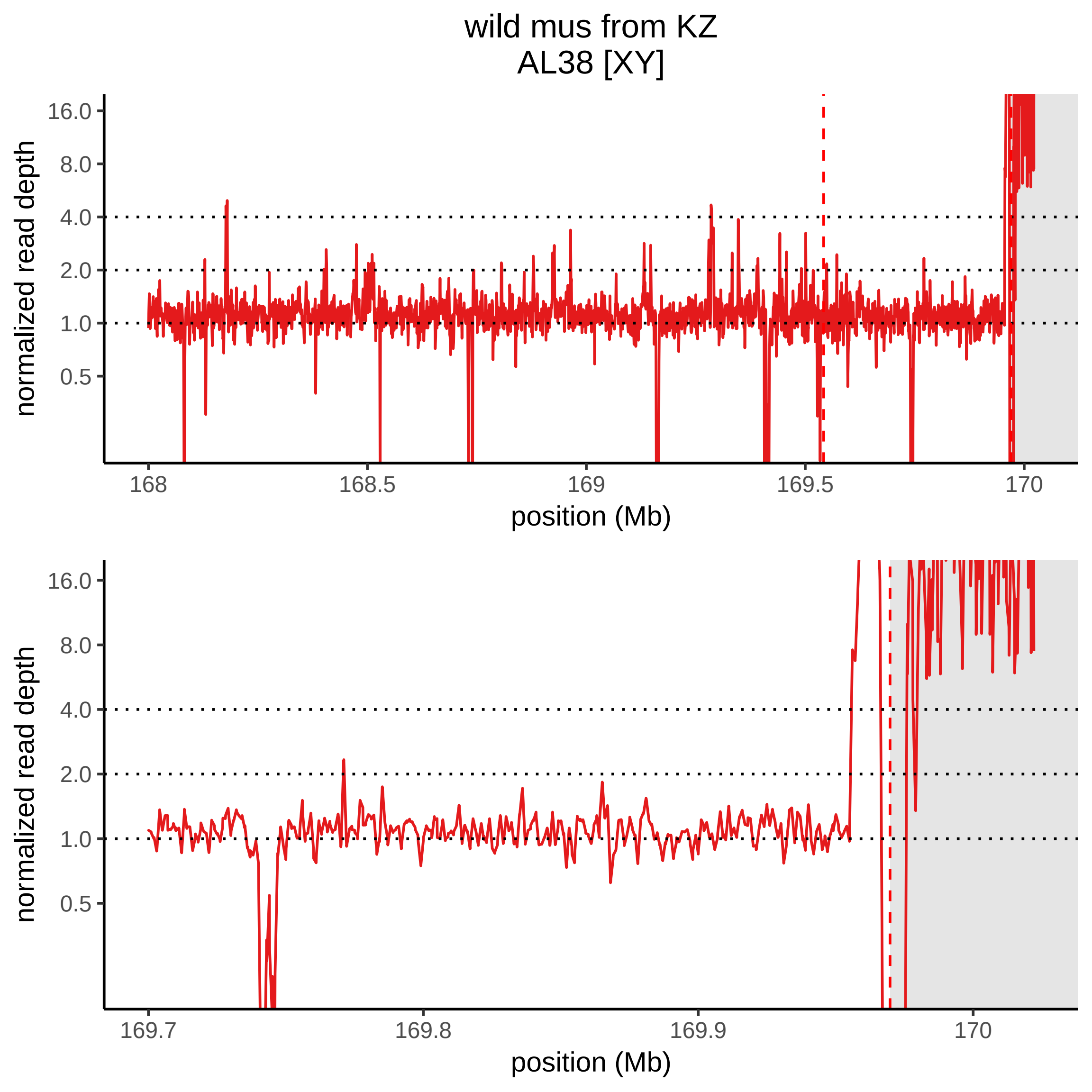
wild mus from KZ
AL38 [XY]
normalized read depth
16.0
8.0
4.0
2.0
1.0
0.5
168
168.5
169
169.5
170
position (Mb)
normalized read depth
16.0
8.0
4.0
2.0
1.0
0.5
169.7
169.8
169.9
170
position (Mb)

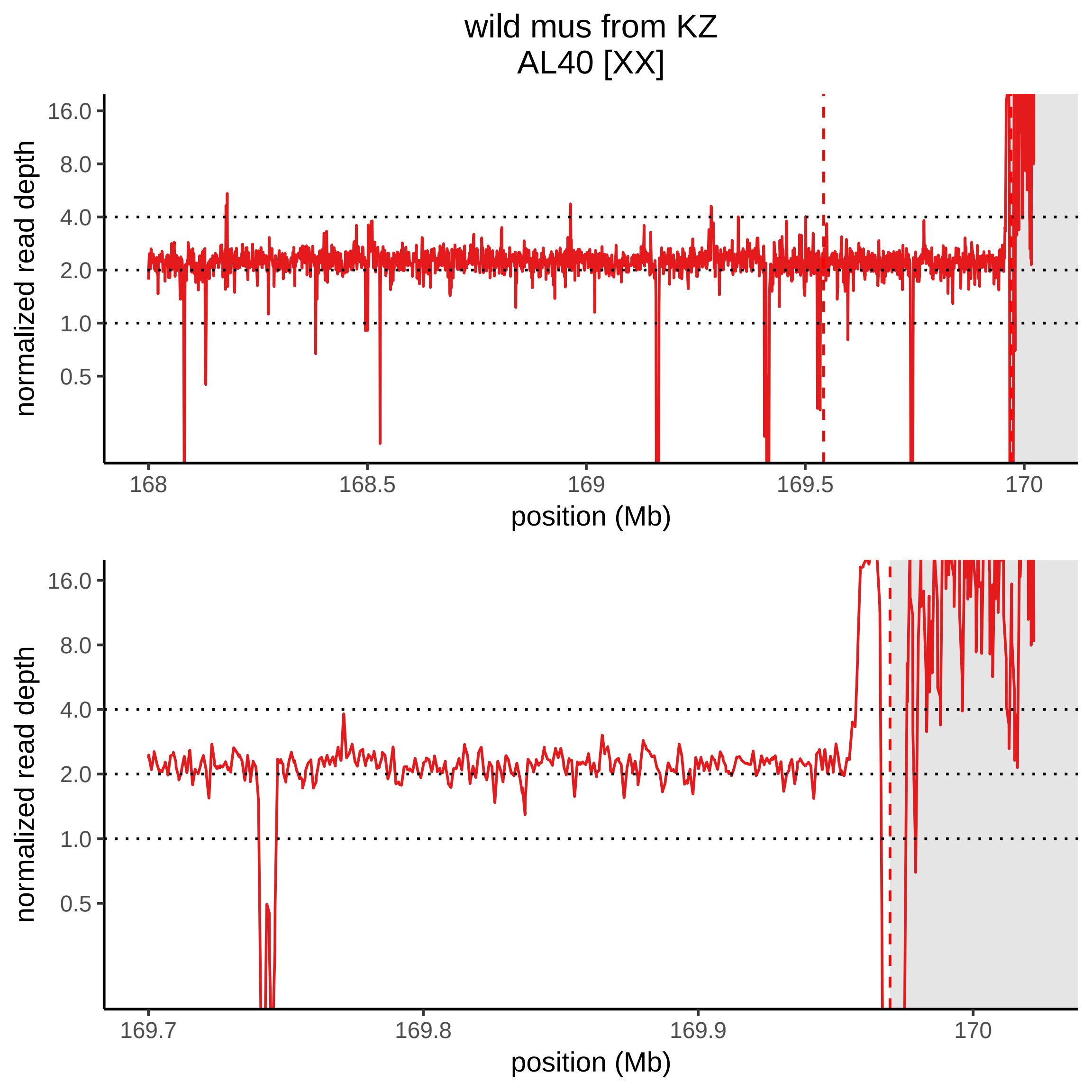

wild mus from KZ
AL40 [XX]
normalized read depth
16.0
8.0
4.0
2.0
1.0
0.5
168
168.5
169
169.5
170
position (Mb)
normalized read depth
16.0
8.0
4.0
2.0
1.0
0.5
169.7
169.8
169.9
170
position (Mb)

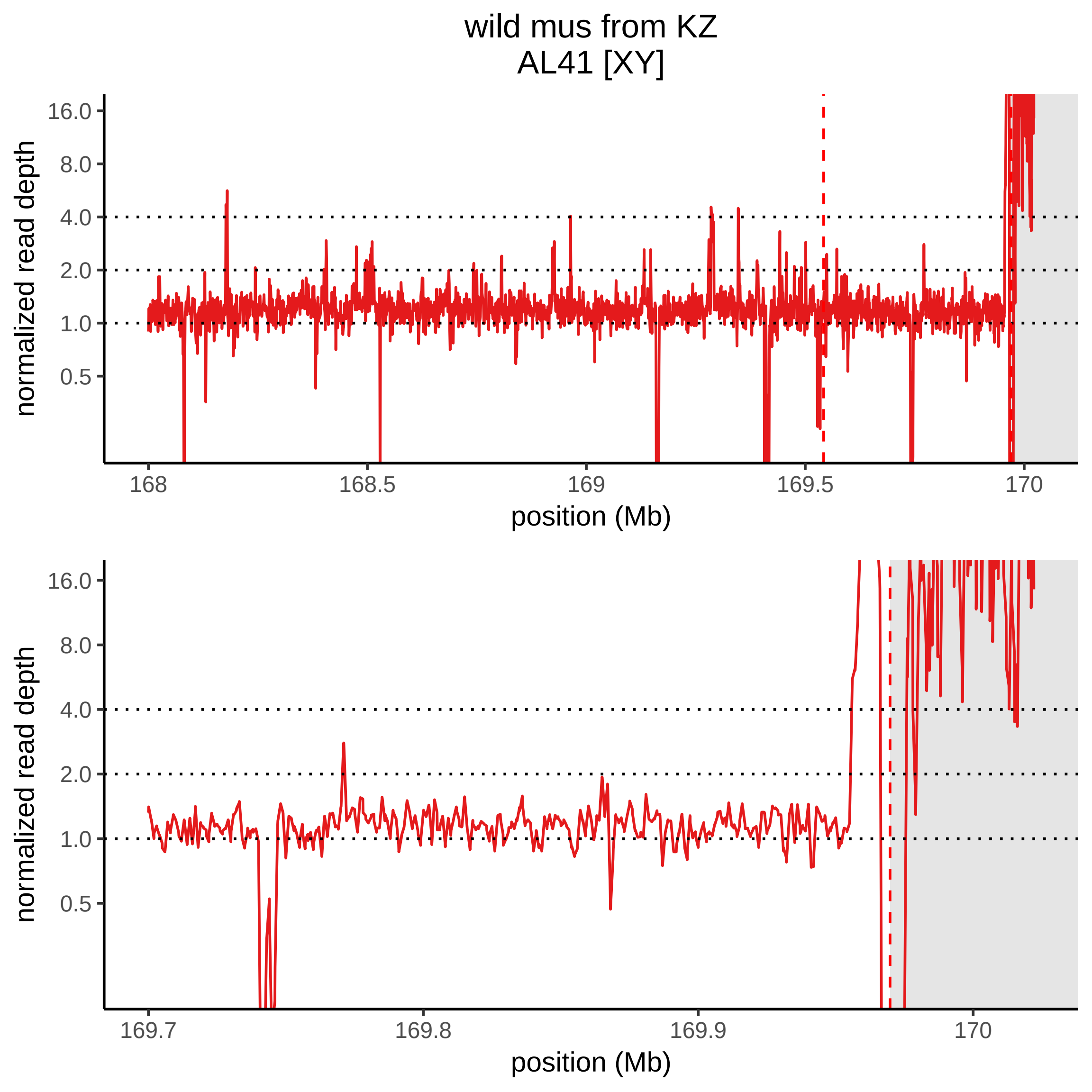
wild mus from KZ
AL41 [XY]
normalized read depth
16.0
8.0
4.0
2.0
1.0
0.5
168
168.5
169
169.5
170
position (Mb)
normalized read depth
16.0
8.0
4.0
2.0
1.0
0.5
169.7
169.8
169.9
170
position (Mb)

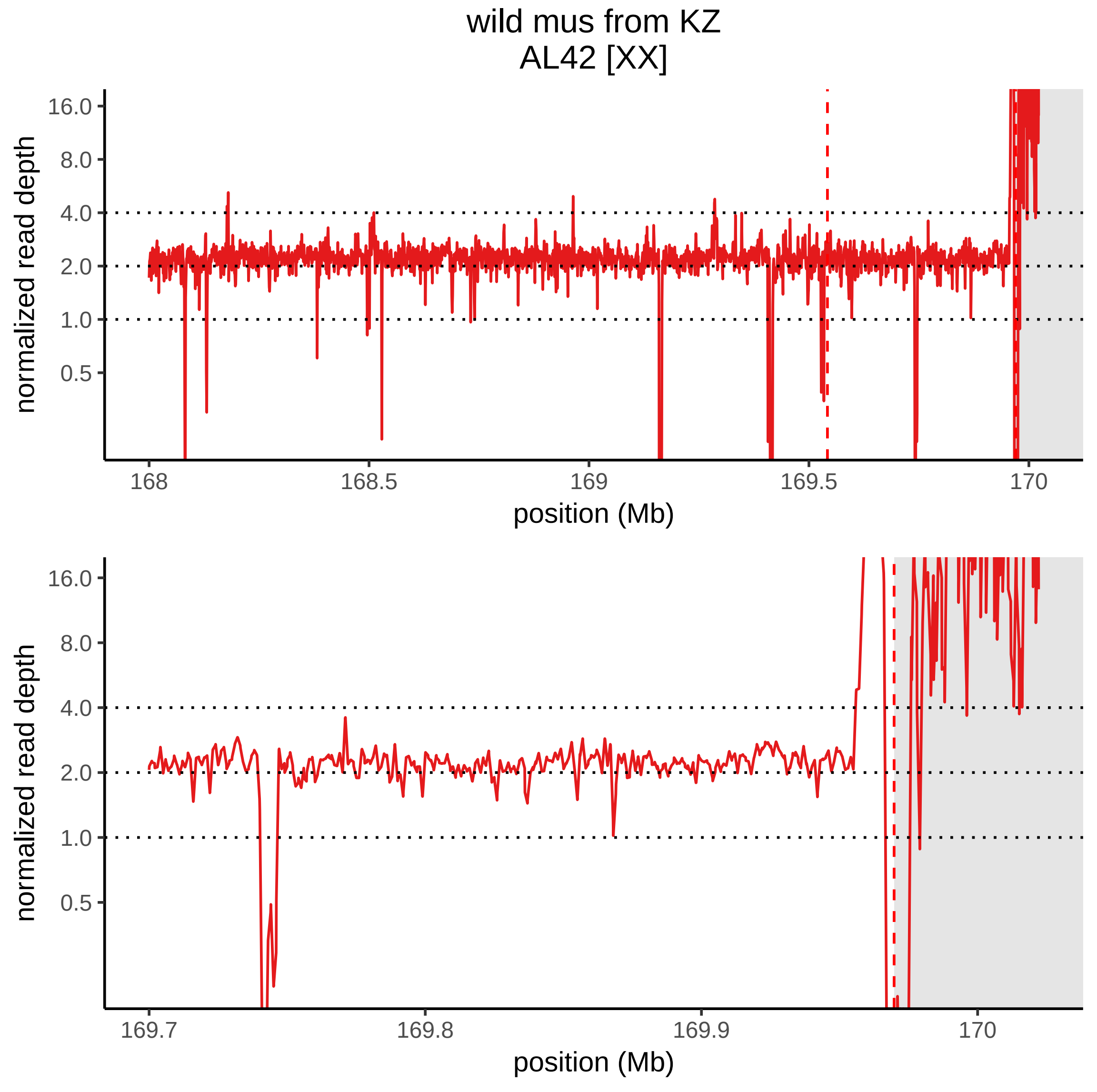

wild mus from KZ
AL42 [XX]
normalized read depth
16.0
8.0
4.0
2.0
1.0
0.5
168
168.5
169
169.5
170
position (Mb)
normalized read depth
16.0
8.0
4.0
2.0
1.0
0.5
169.7
169.8
169.9
170
position (Mb)

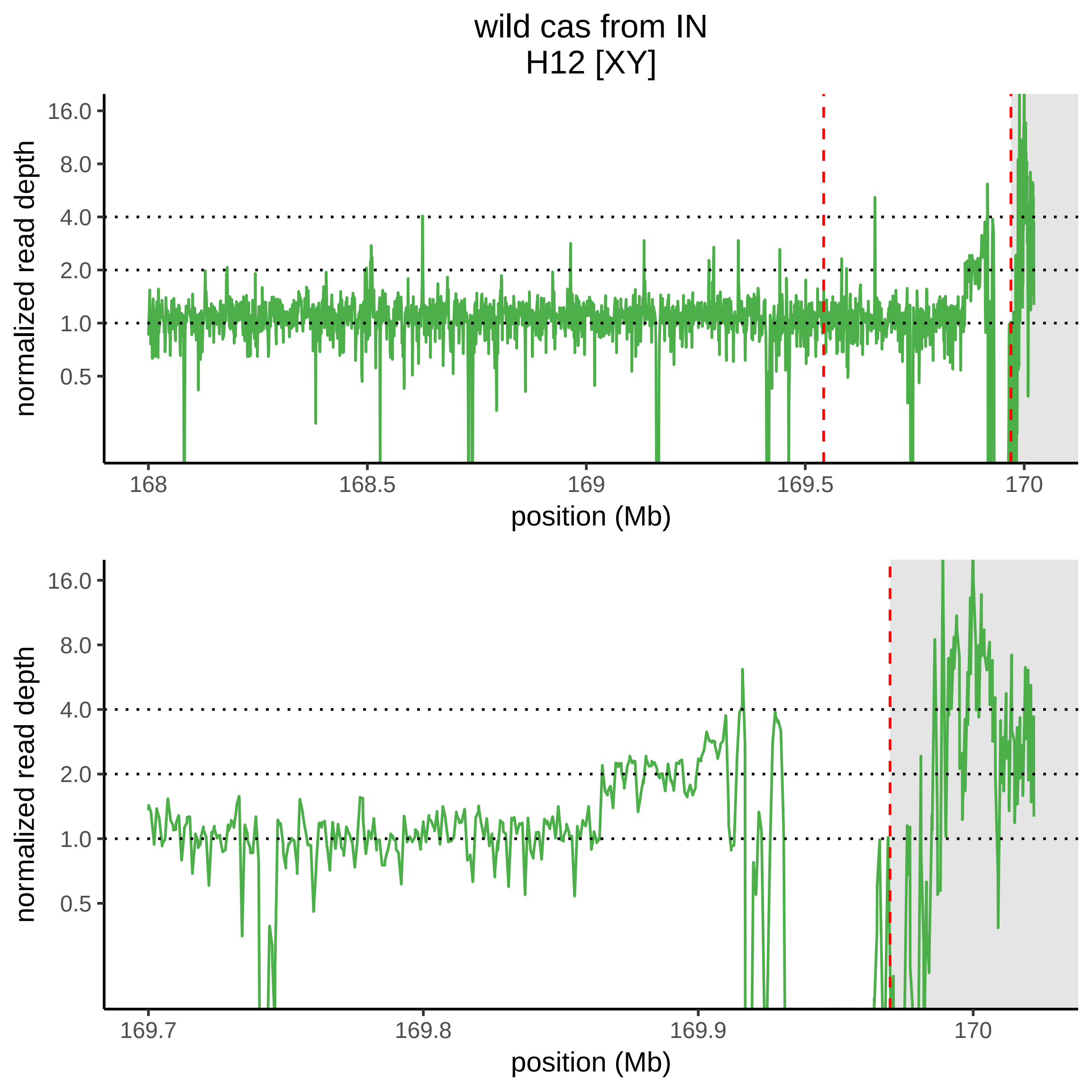
wild cas from IN
H12 [XY]
normalized read depth
16.0
8.0
4.0
2.0
1.0
0.5
168
168.5
169
169.5
170
position (Mb)
normalized read depth
16.0
8.0
4.0
2.0
1.0
0.5
169.7
169.8
169.9
170
position (Mb)

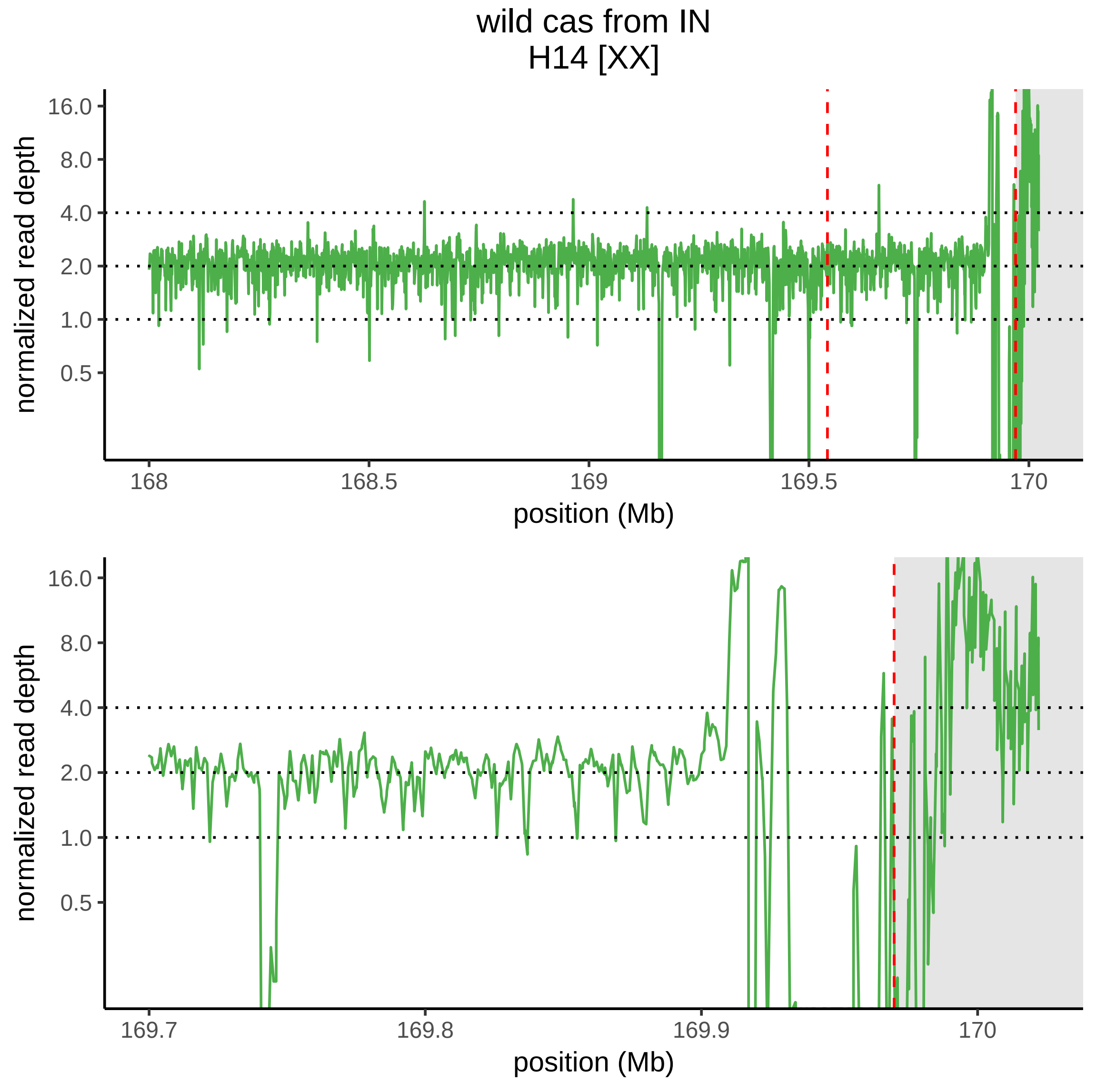

wild cas from IN
H14 [XX]
normalized read depth
16.0
8.0
4.0
2.0
1.0
0.5
168
168.5
169
169.5
170
position (Mb)
normalized read depth
16.0
8.0
4.0
2.0
1.0
0.5
169.7
169.8
169.9
170
position (Mb)

wild cas from IN
H15 [XX]

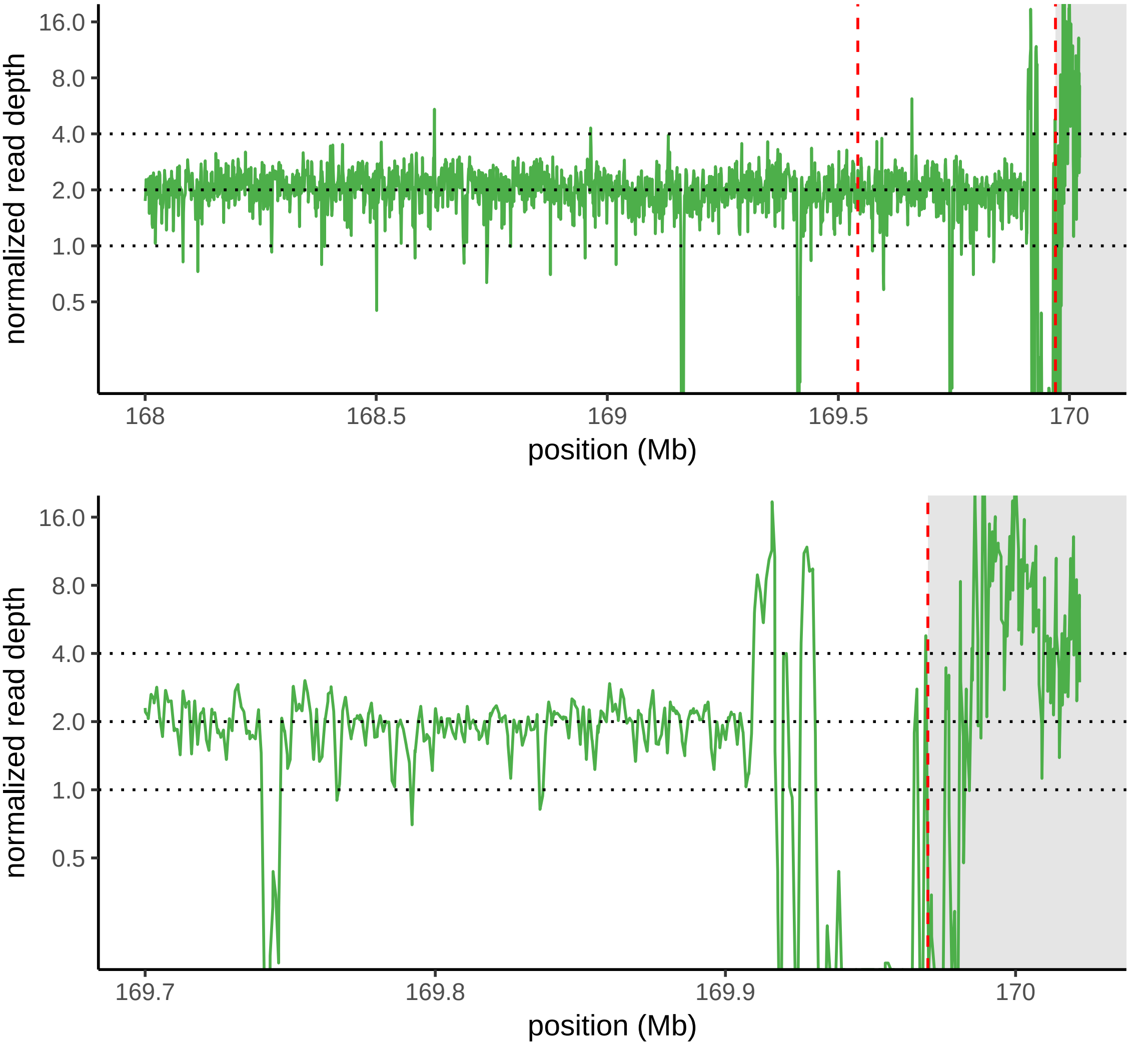

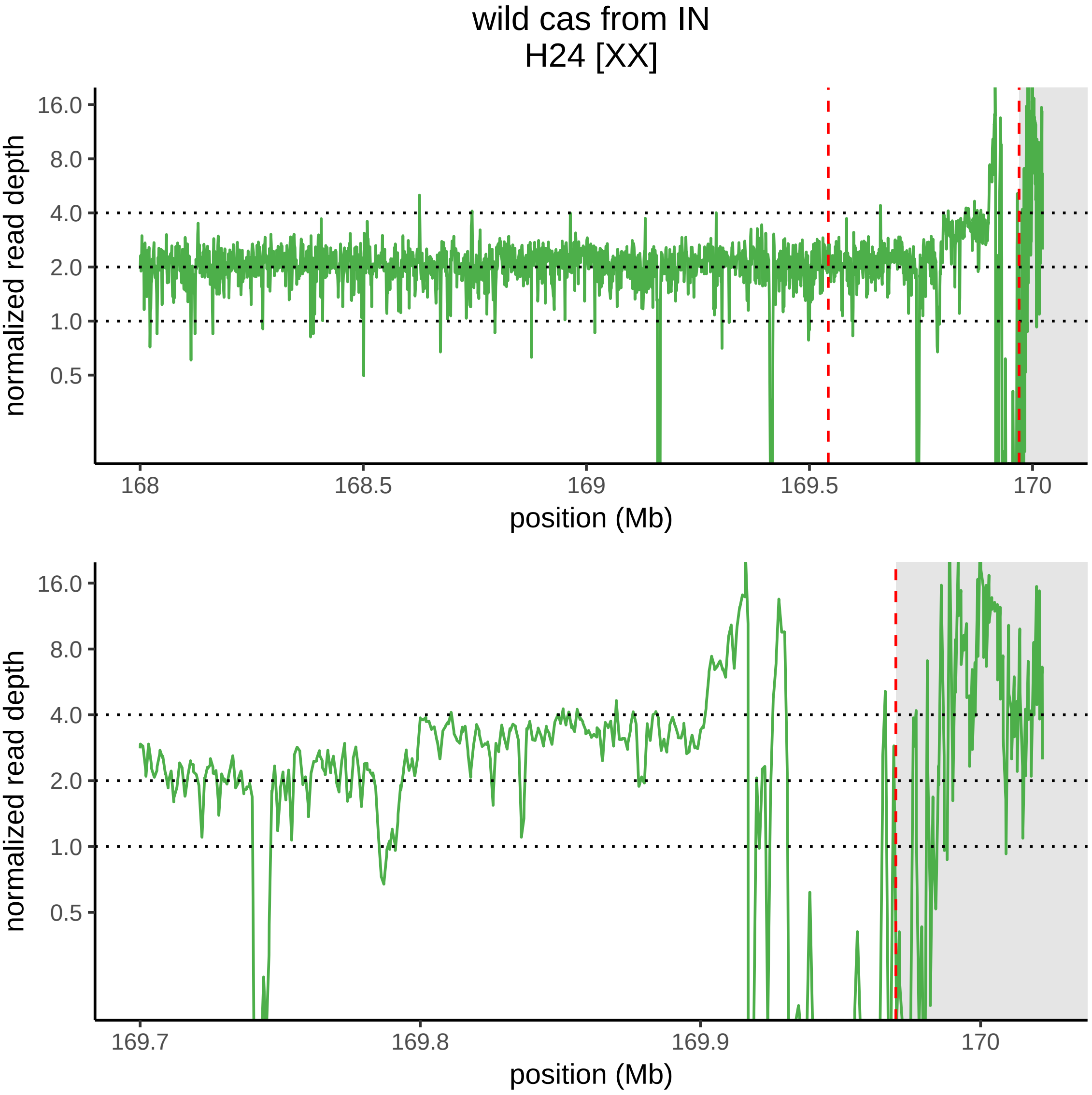

wild cas from IN
H24 [XX]
normalized read depth
16.0
8.0
4.0
2.0
1.0
0.5
168
168.5
169
169.5
170
position (Mb)
normalized read depth
16.0
8.0
4.0
2.0
1.0
0.5
169.7
169.8
169.9
170
position (Mb)

wild cas from IN
H26 [XX]

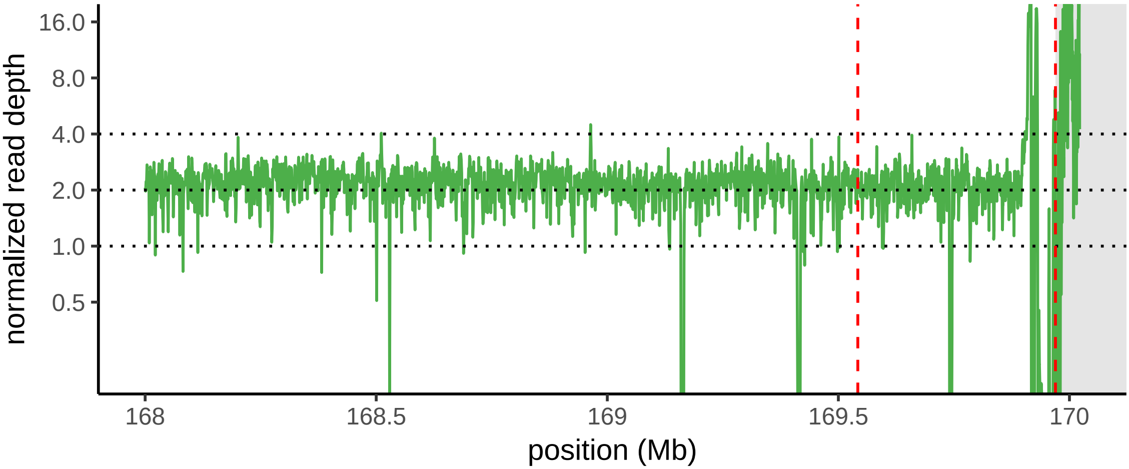


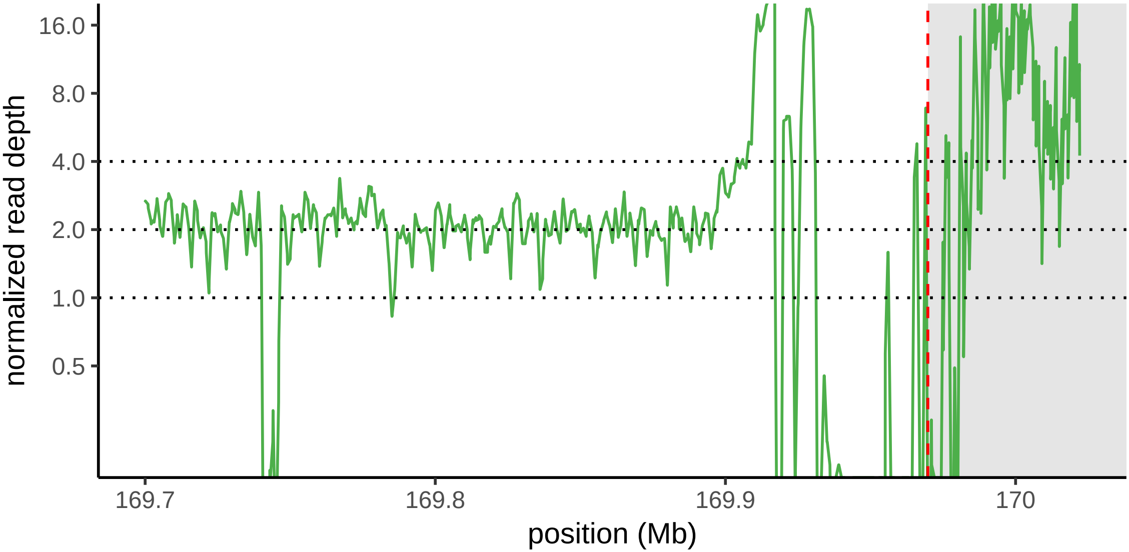

wild cas from IN
H27 [XX]

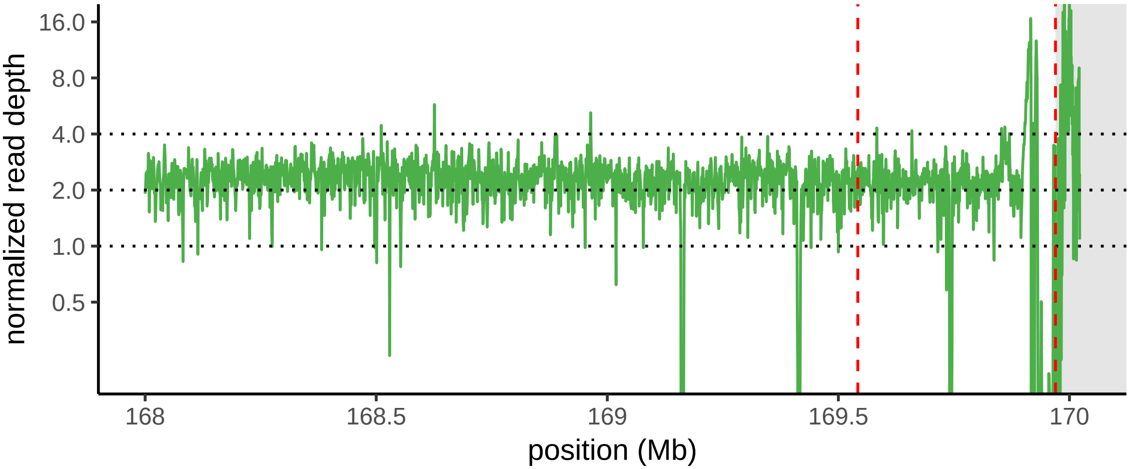


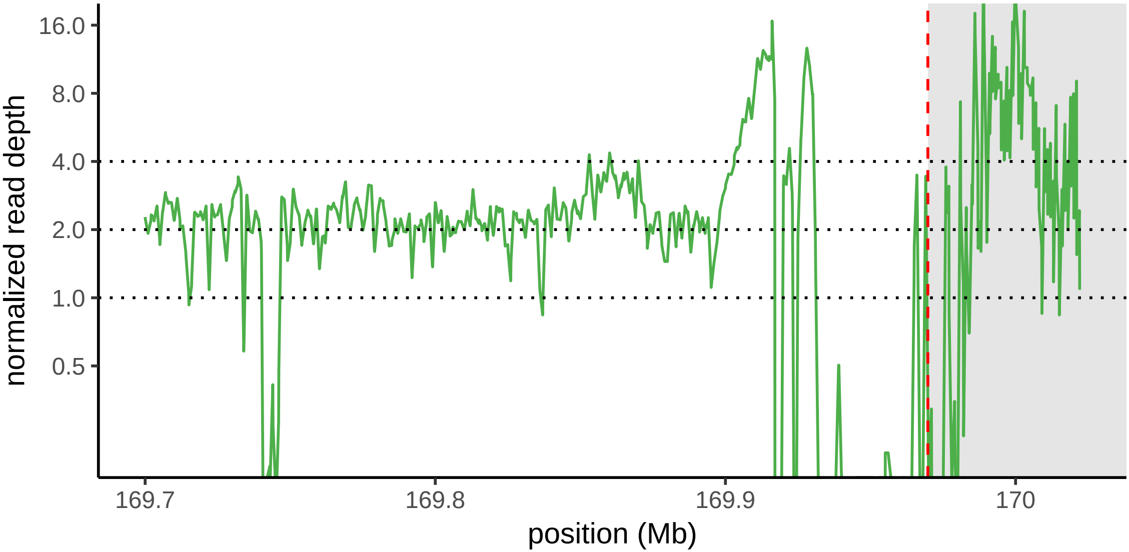

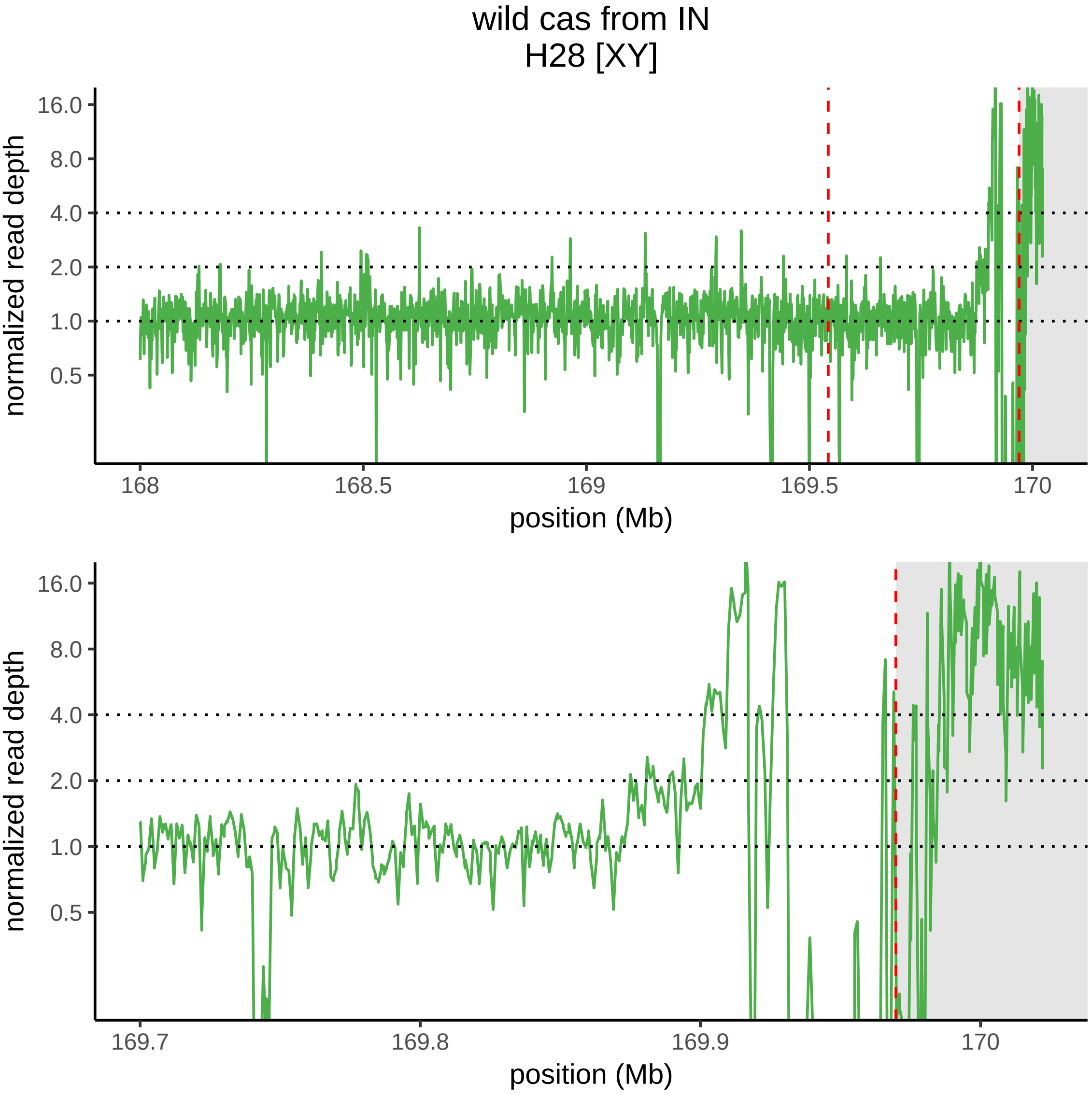
wild cas from IN
H28 [XY]
normalized read depth
16.0
8.0
4.0
2.0
1.0
0.5
168
168.5
169
169.5
170
position (Mb)
normalized read depth
16.0
8.0
4.0
2.0
1.0
0.5
169.7
169.8
169.9
170
position (Mb)

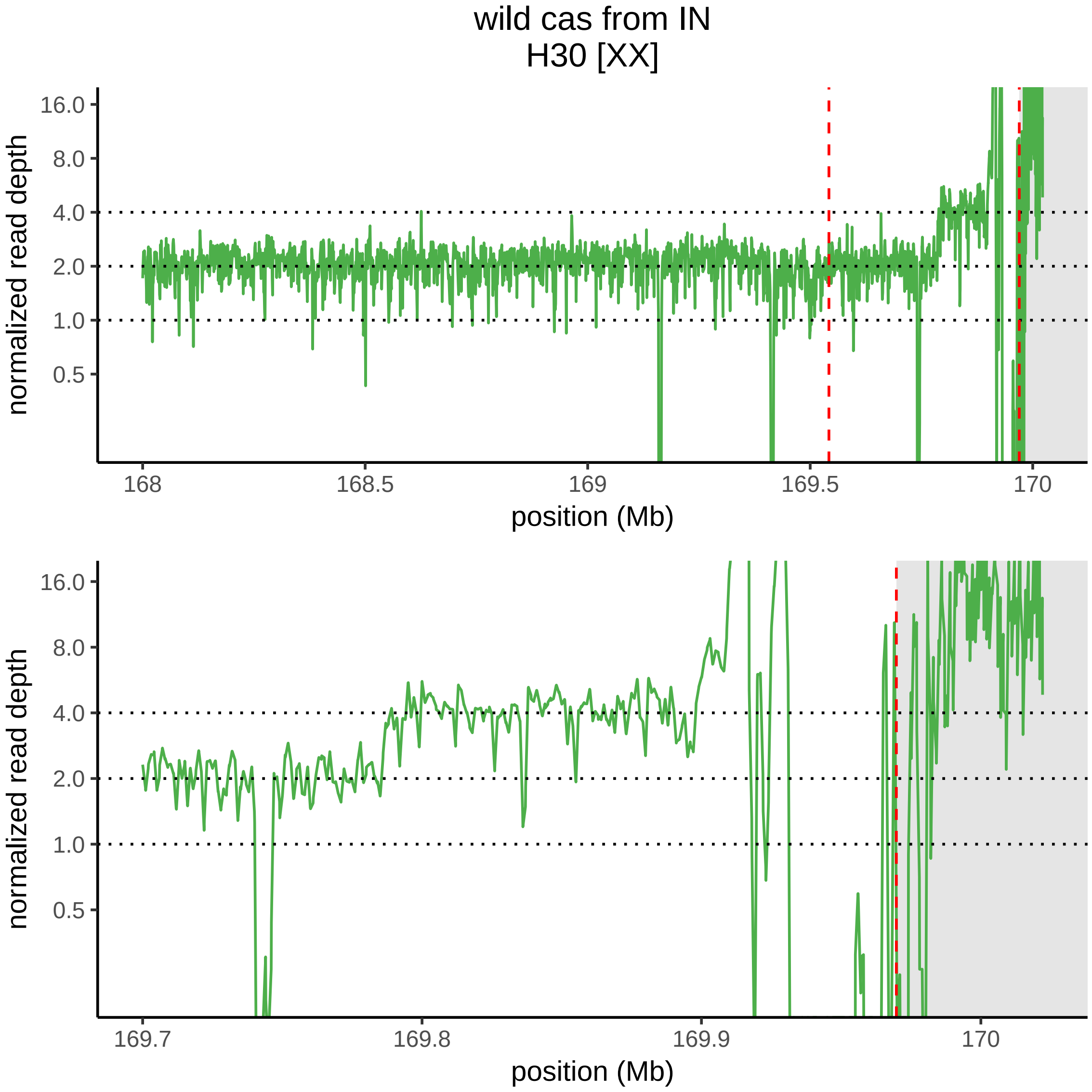
wild cas from IN
H30 [XX]
normalized read depth
16.0
8.0
4.0
2.0
1.0
0.5
168
168.5
169
169.5
170
position (Mb)
normalized read depth
16.0
8.0
4.0
2.0
1.0
0.5
169.7
169.8
169.9
170
position (Mb)

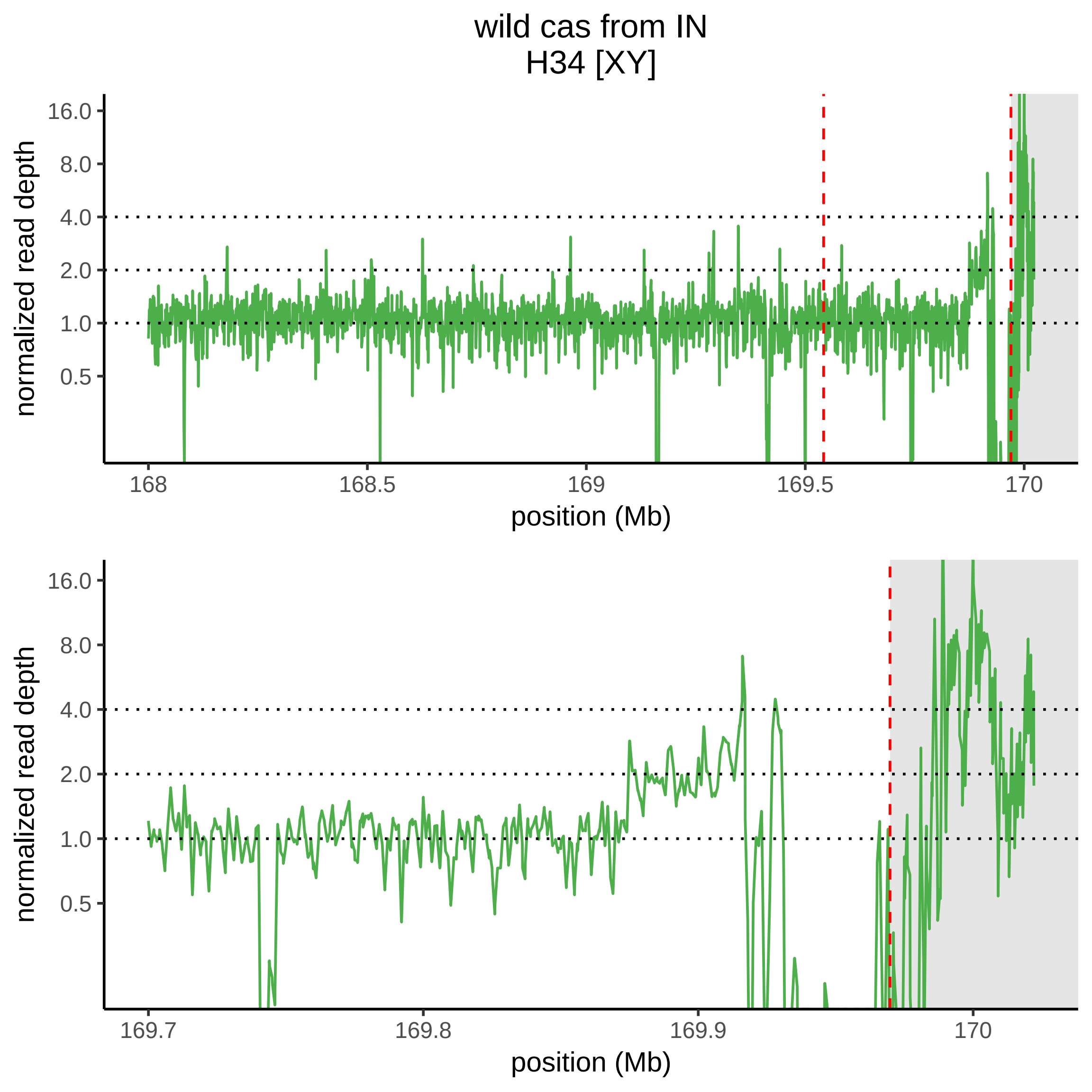
wild cas from IN
H34 [XY]
normalized read depth
16.0
8.0
4.0
2.0
1.0
0.5
168
168.5
169
169.5
170
position (Mb)
normalized read depth
16.0
8.0
4.0
2.0
1.0
0.5
169.7
169.8
169.9
170
position (Mb)

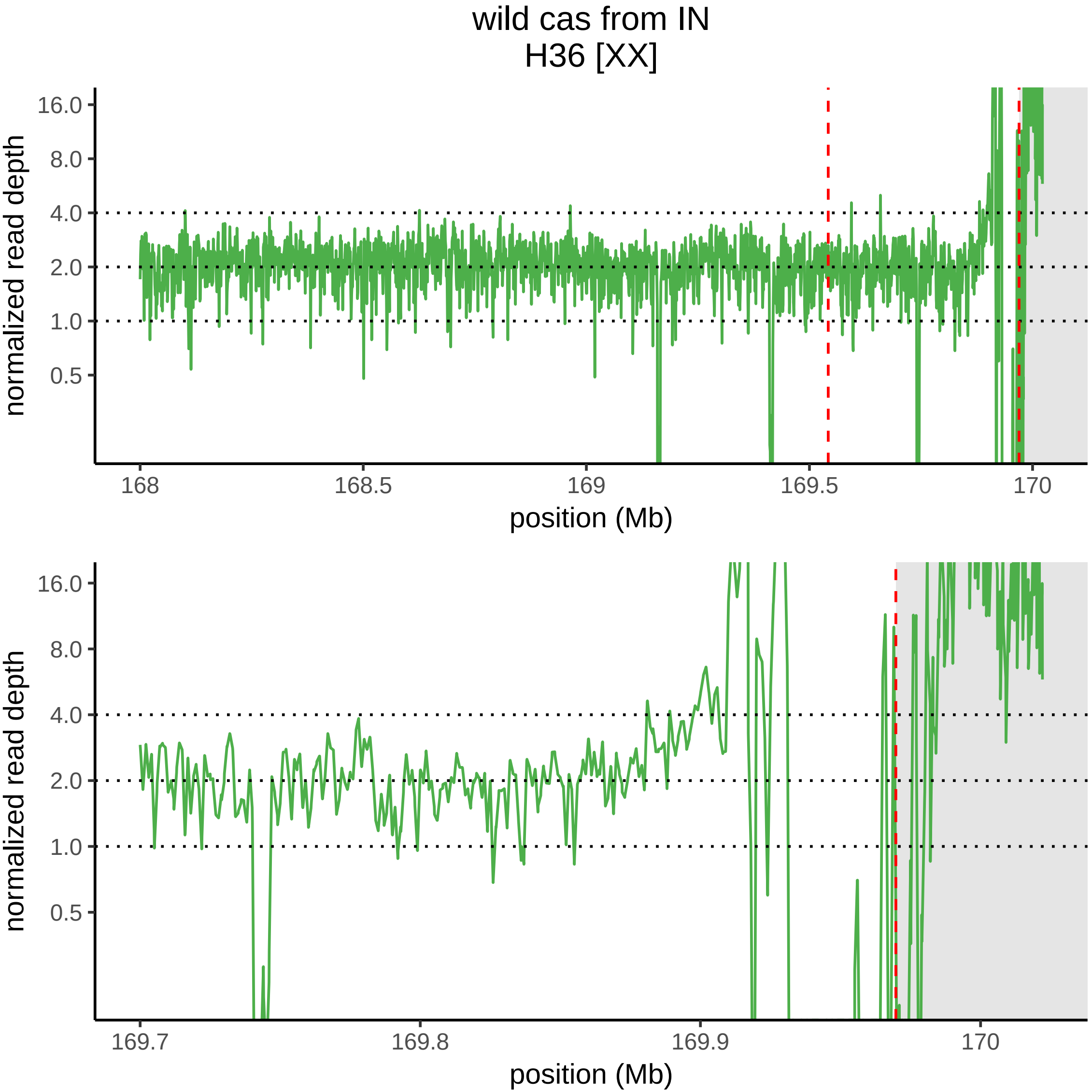

wild cas from IN
H36 [XX]
normalized read depth
16.0
8.0
4.0
2.0
1.0
0.5
168
168.5
169
169.5
170
position (Mb)
normalized read depth
16.0
8.0
4.0
2.0
1.0
0.5
169.7
169.8
169.9
170
position (Mb)

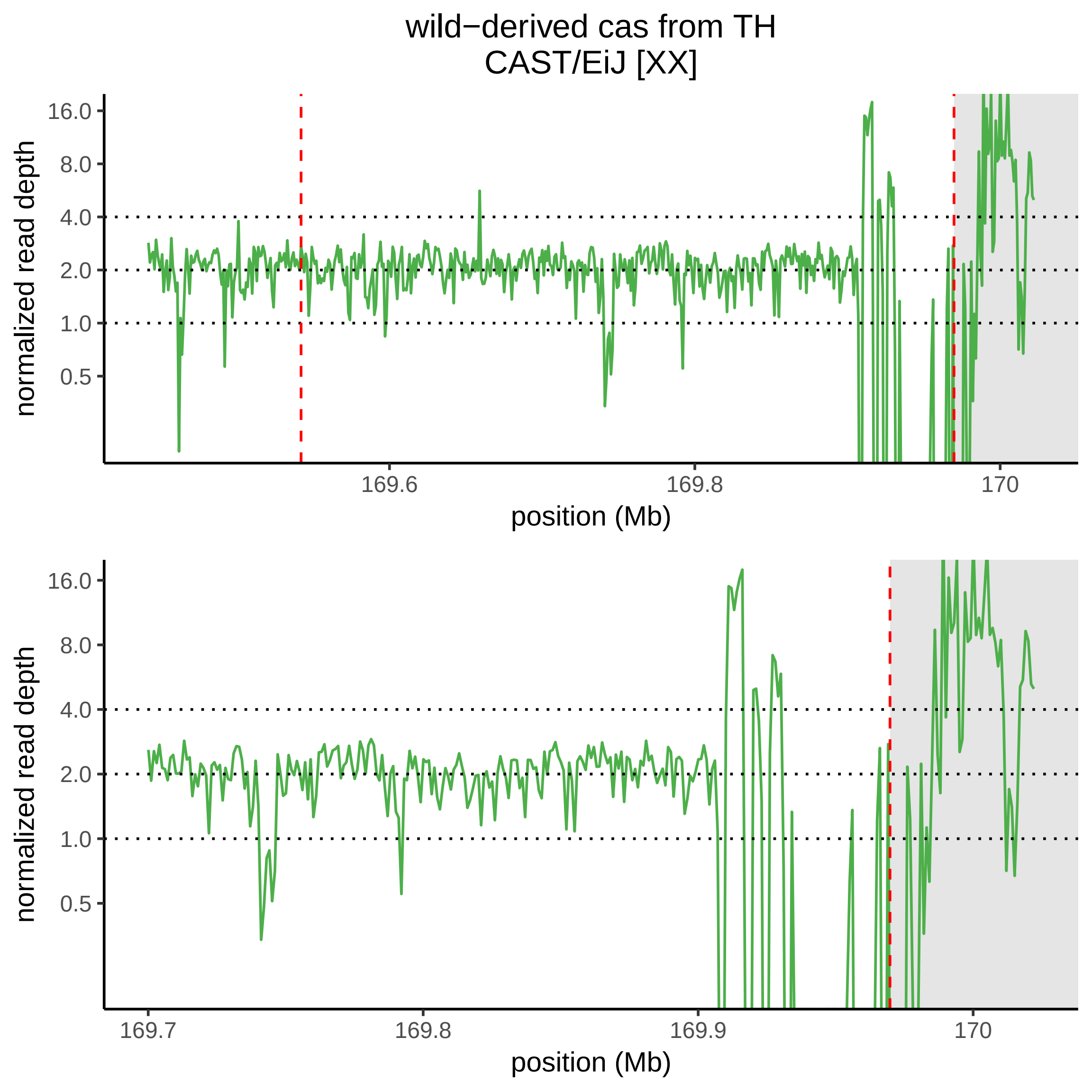

wild−derived cas from TH
CAST/EiJ [XX]
normalized read depth
16.0
8.0
4.0
2.0
1.0
0.5
169.6
169.8
170
position (Mb)
normalized read depth
16.0
8.0
4.0
2.0
1.0
0.5
169.7
169.8
169.9
170
position (Mb)

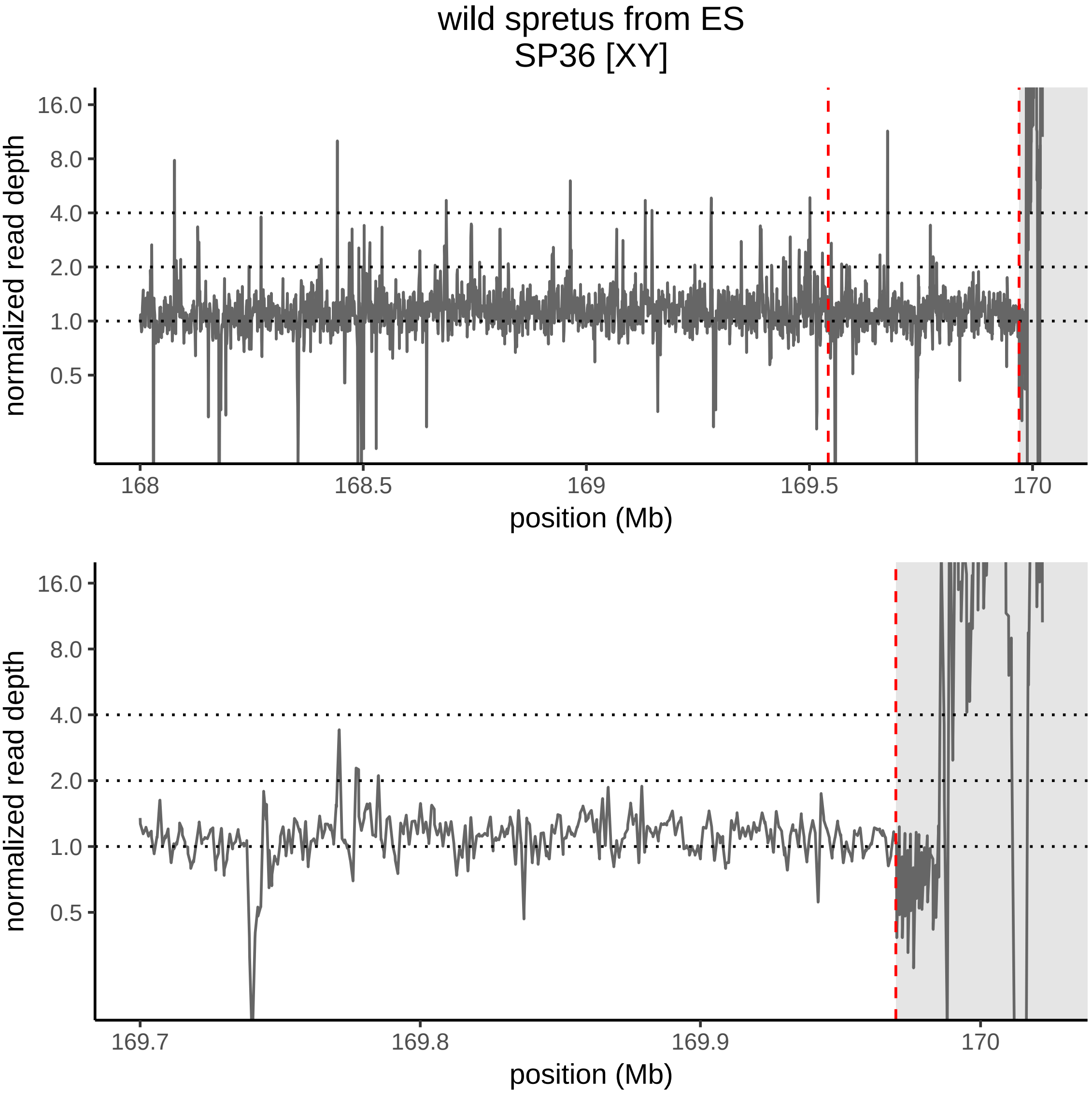
wild spretus from ES
SP36 [XY]
normalized read depth
16.0
8.0
4.0
2.0
1.0
0.5
168
168.5
169
169.5
170
position (Mb)
normalized read depth
16.0
8.0
4.0
2.0
1.0
0.5
169.7
169.8
169.9
170
position (Mb)

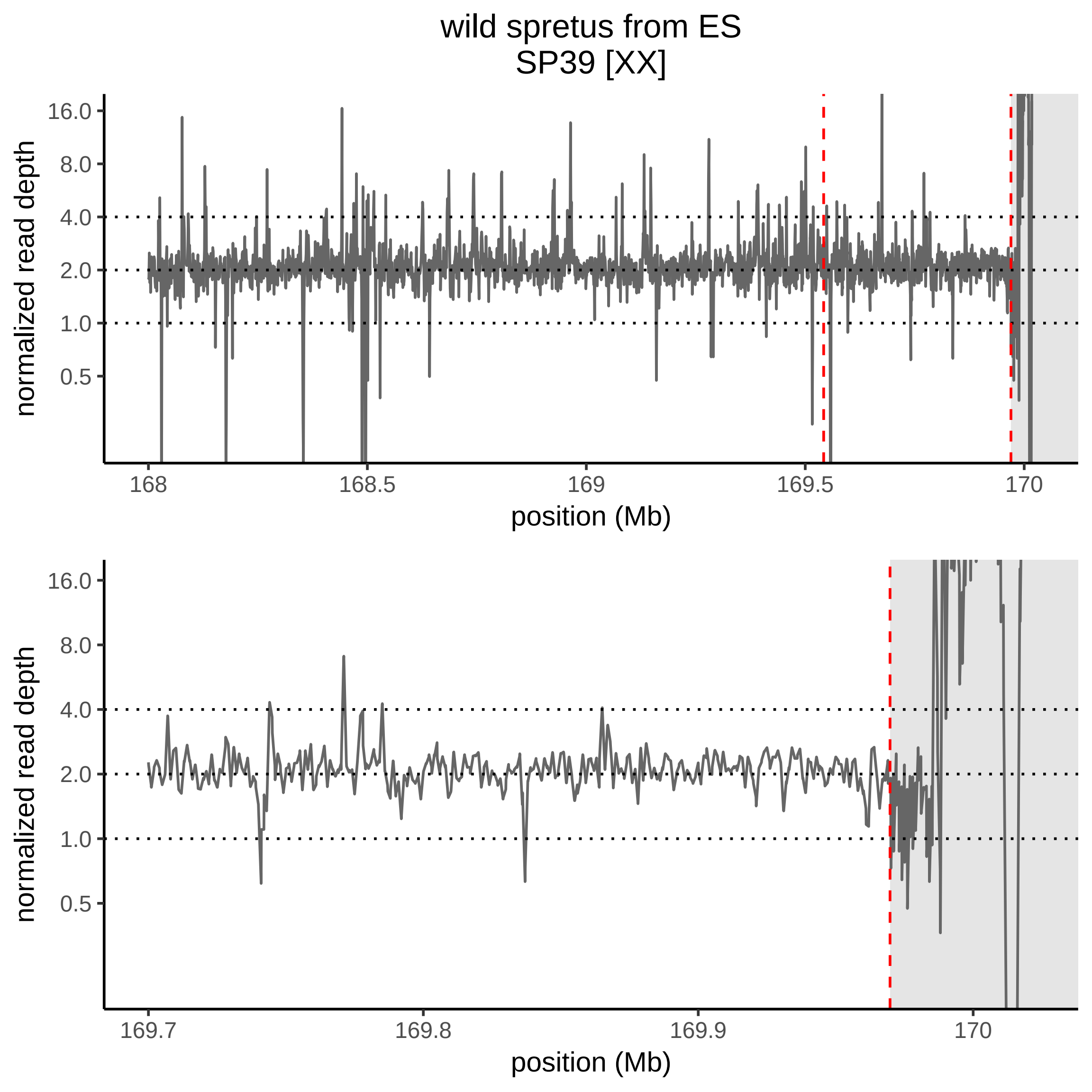
wild spretus from ES
SP39 [XX]
normalized read depth
16.0
8.0
4.0
2.0
1.0
0.5
168
168.5
169
169.5
170
position (Mb)
normalized read depth
16.0
8.0
4.0
2.0
1.0
0.5
169.7
169.8
169.9
170
position (Mb)

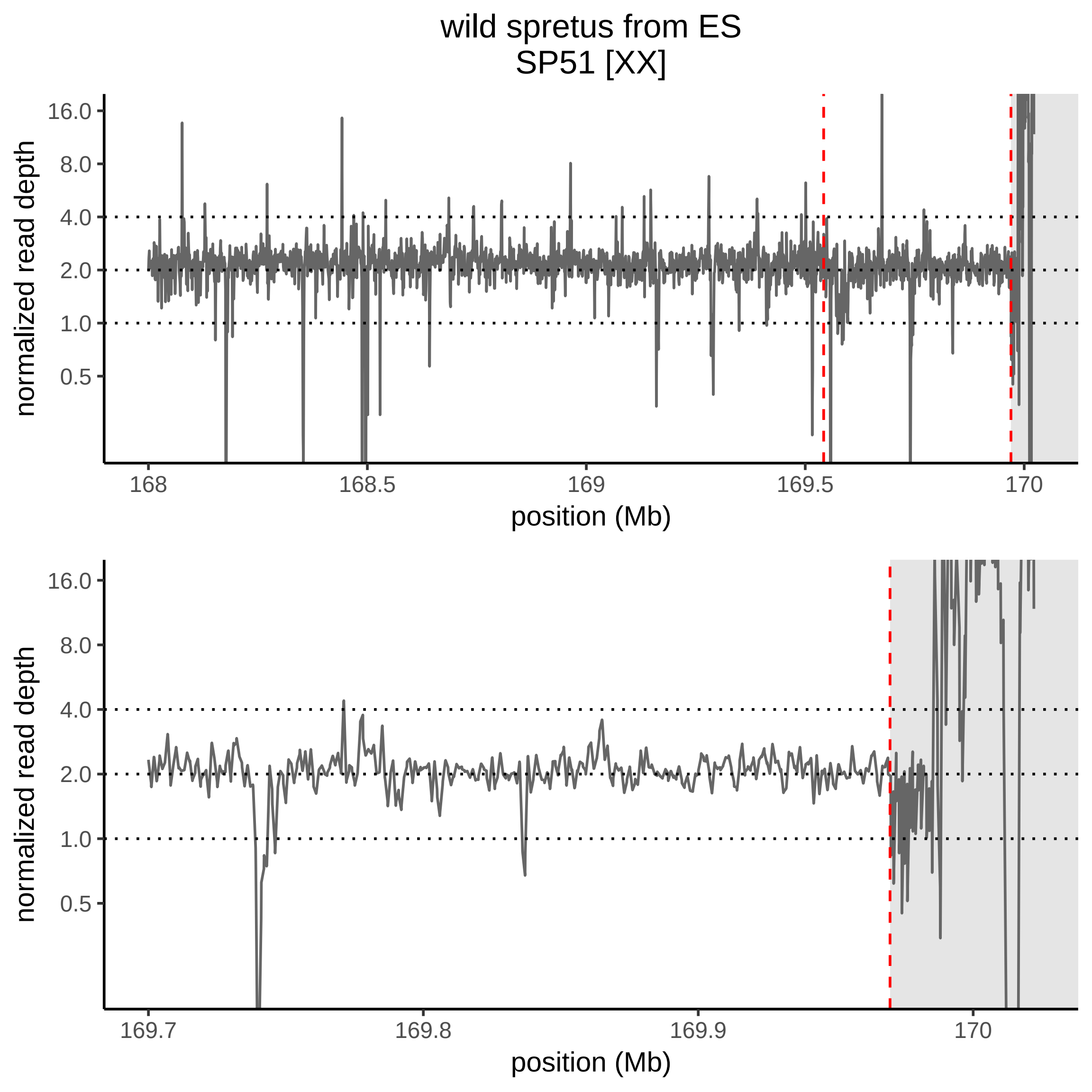
wild spretus from ES
SP51 [XX]
normalized read depth
16.0
8.0
4.0
2.0
1.0
0.5
168
168.5
169
169.5
170
position (Mb)
normalized read depth
16.0
8.0
4.0
2.0
1.0
0.5
169.7
169.8
169.9
170
position (Mb)

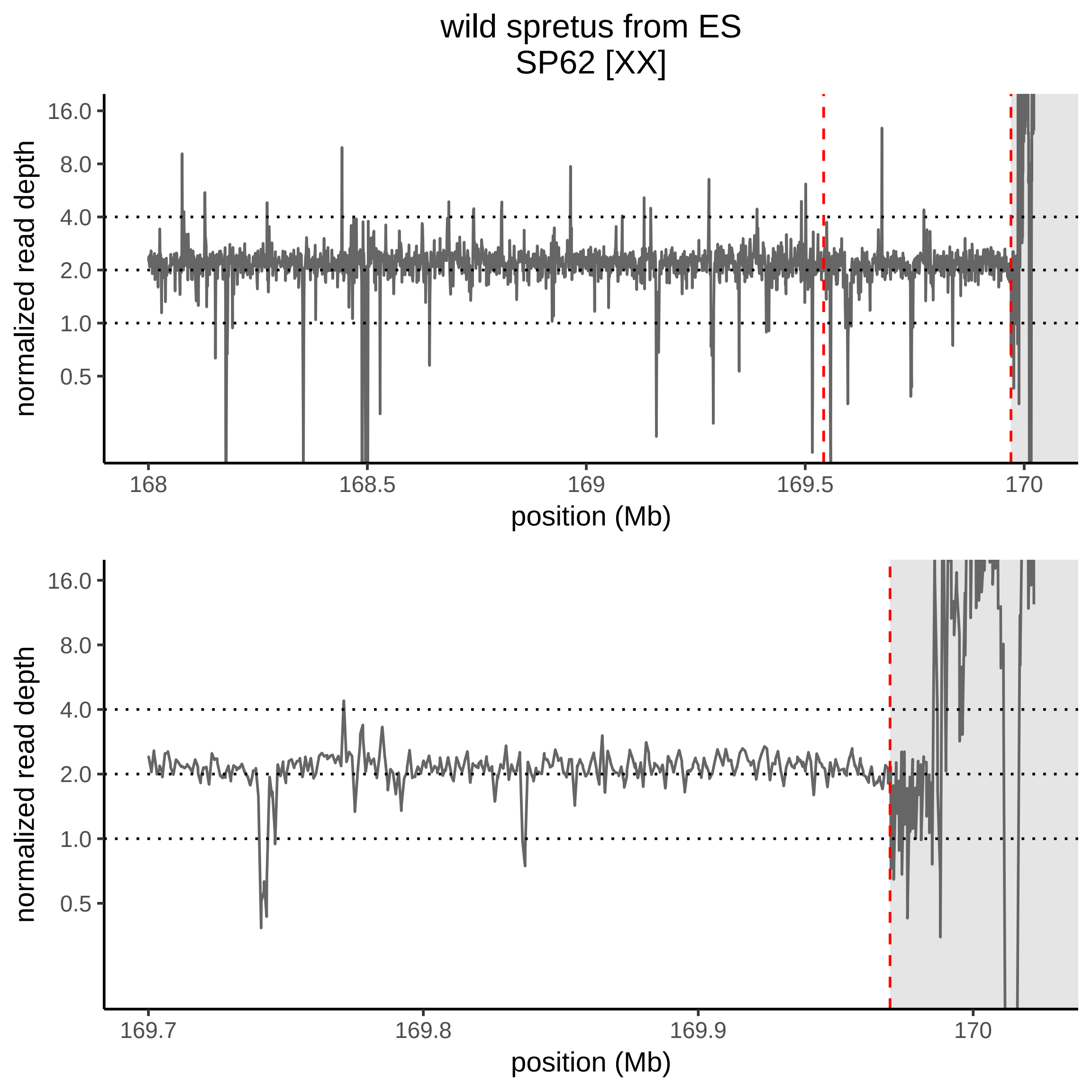

wild spretus from ES
SP62 [XX]
normalized read depth
16.0
8.0
4.0
2.0
1.0
0.5
168
168.5
169
169.5
170
position (Mb)
normalized read depth
16.0
8.0
4.0
2.0
1.0
0.5
169.7
169.8
169.9
170
position (Mb)

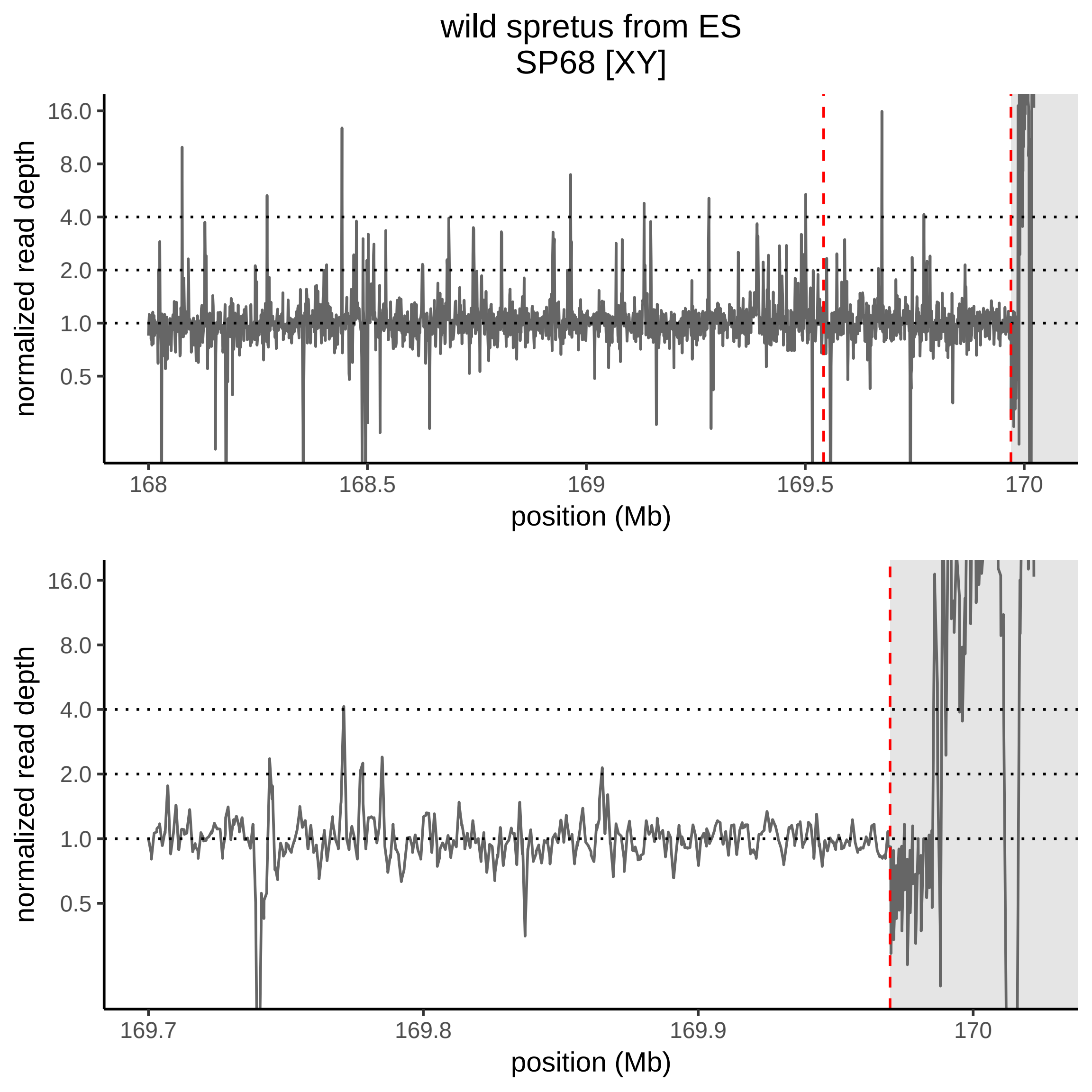

wild spretus from ES
SP68 [XY]
normalized read depth
16.0
8.0
4.0
2.0
1.0
0.5
168
168.5
169
169.5
170
position (Mb)
normalized read depth
16.0
8.0
4.0
2.0
1.0
0.5
169.7
169.8
169.9
170
position (Mb)

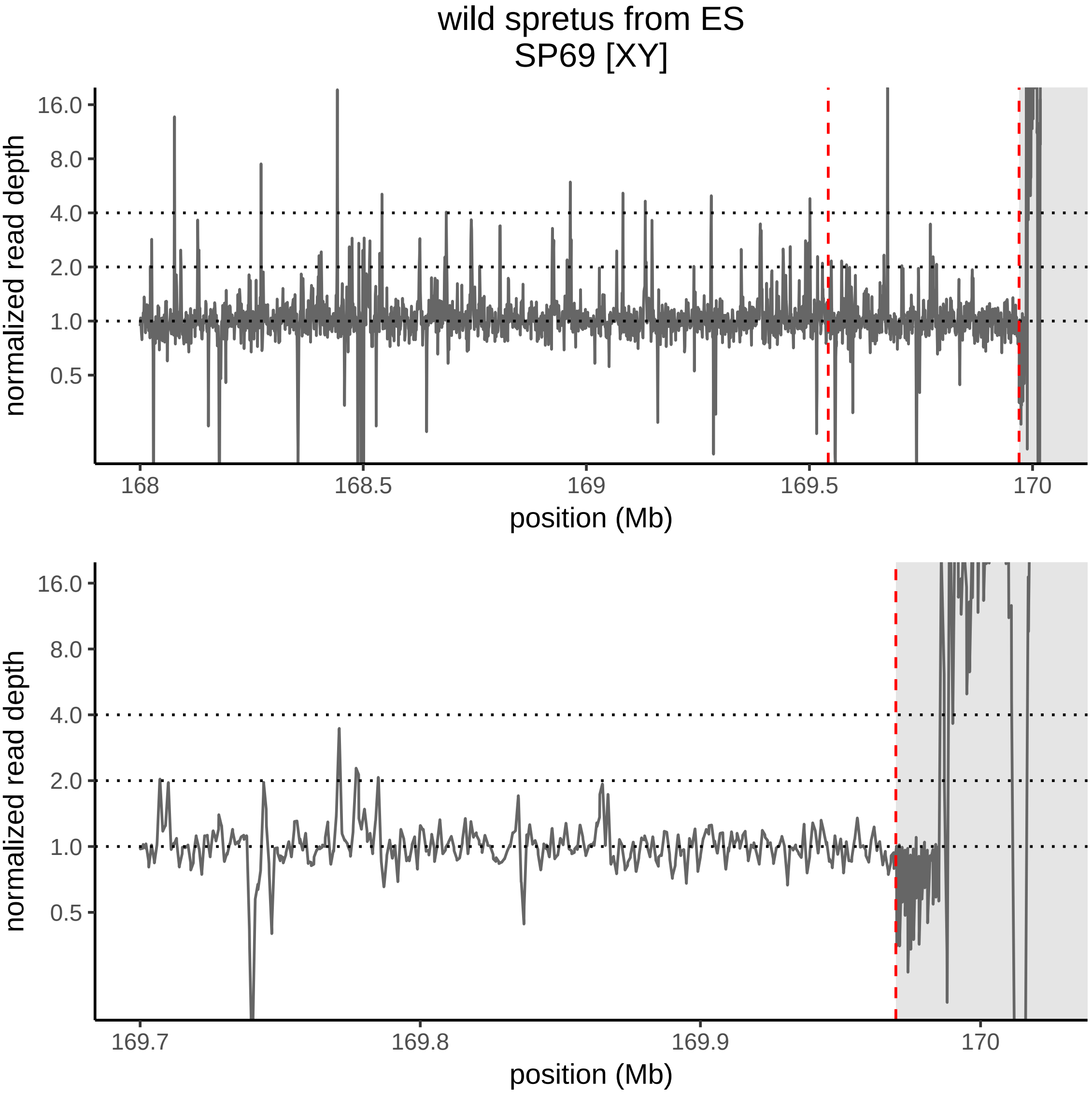

wild spretus from ES
SP69 [XY]
normalized read depth
16.0
8.0
4.0
2.0
1.0
0.5
168
168.5
169
169.5
170
position (Mb)
normalized read depth
16.0
8.0
4.0
2.0
1.0
0.5
169.7
169.8
169.9
170
position (Mb)

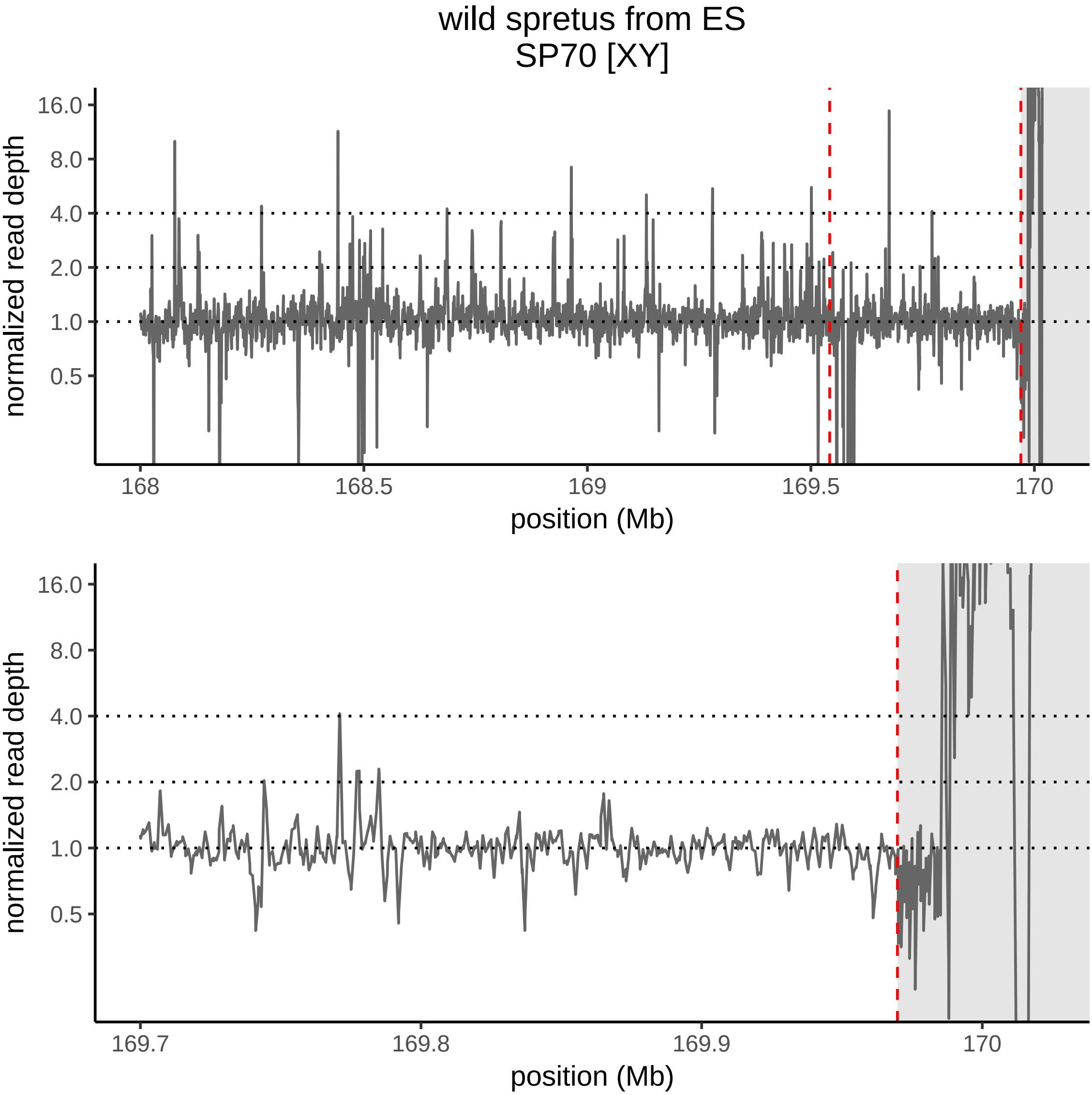
wild spretus from ES
SP70 [XY]
normalized read depth
16.0
8.0
4.0
2.0
1.0
0.5
168
168.5
169
169.5
170
position (Mb)
normalized read depth
16.0
8.0
4.0
2.0
1.0
0.5
169.7
169.8
169.9
170
position (Mb)

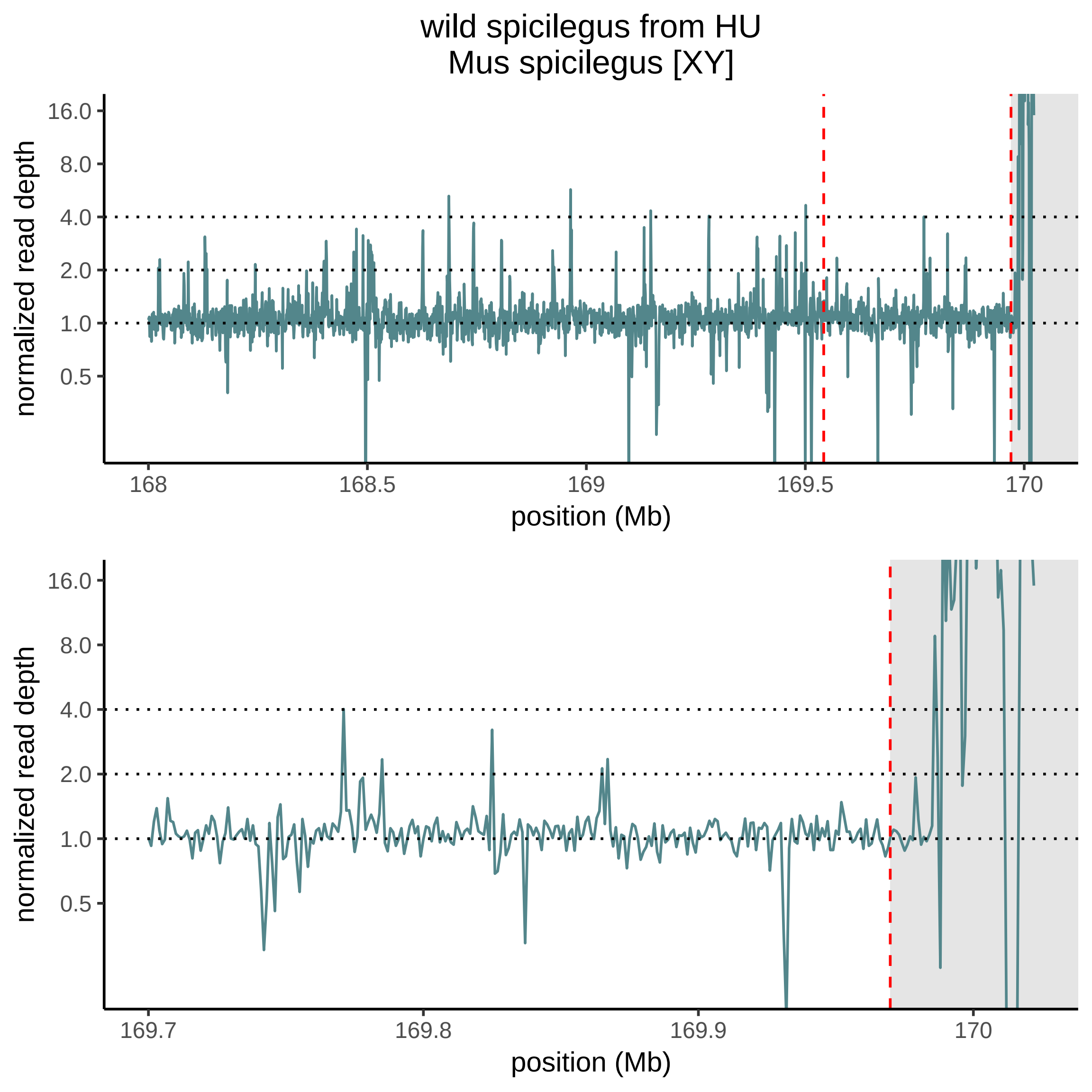
wild spicilegus from HU
Mus spicilegus [XY]
normalized read depth
16.0
8.0
4.0
2.0
1.0
0.5
168
168.5
169
169.5
170
position (Mb)
normalized read depth
16.0
8.0
4.0
2.0
1.0
0.5
169.7
169.8
169.9
170
position (Mb)

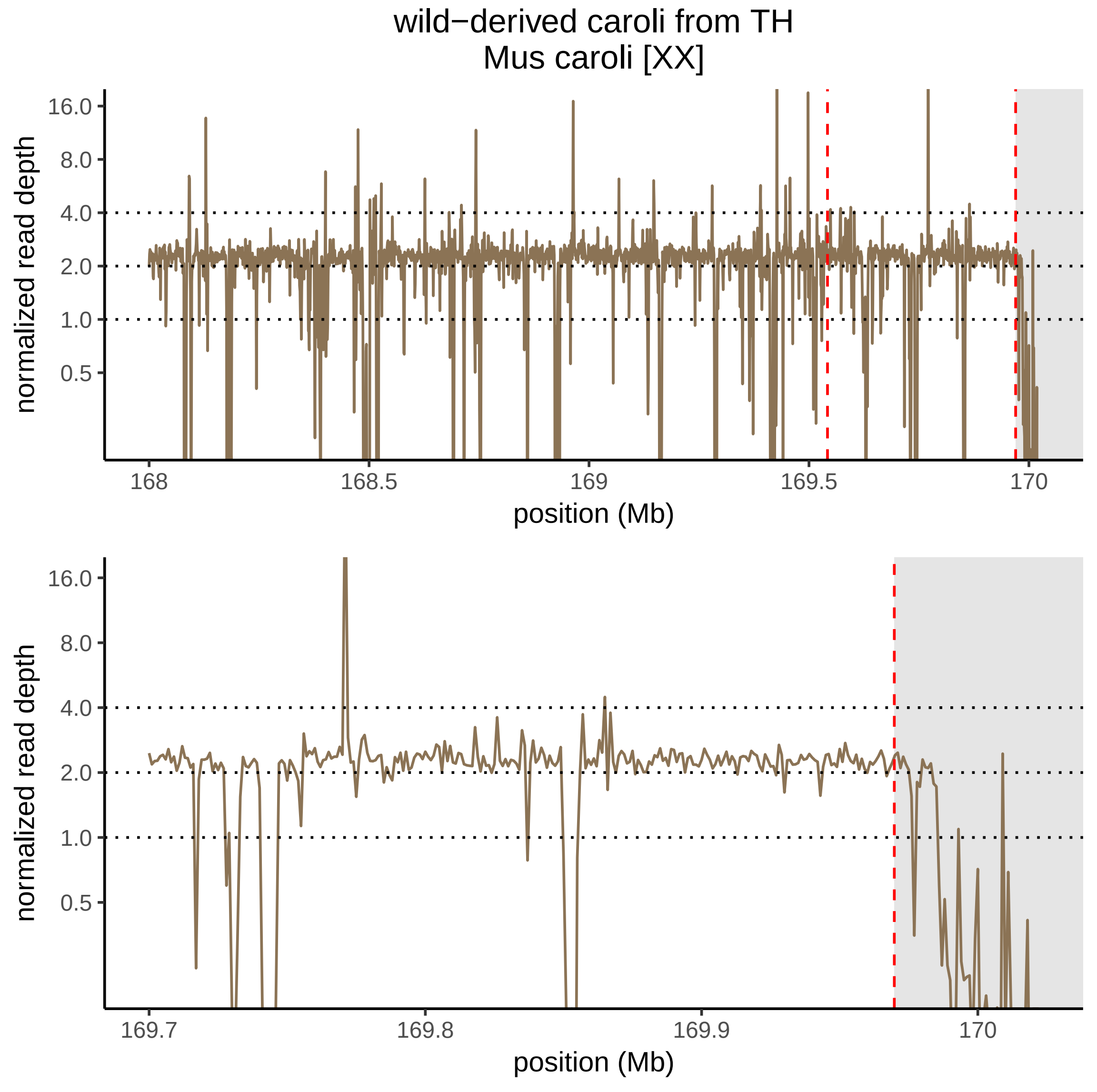

wild−derived caroli from TH
Mus caroli [XX]
normalized read depth
16.0
8.0
4.0
2.0
1.0
0.5
168
168.5
169
169.5
170
position (Mb)
normalized read depth
16.0
8.0
4.0
2.0
1.0
0.5
169.7
169.8
169.9
170
position (Mb)
